# Thermodynamic Parameter Estimation for Modified Oligonucleotides Using Molecular Dynamics Simulations

**DOI:** 10.1101/2024.11.29.626002

**Authors:** Soon Woo Park, Junehawk Lee, Jung Woo Park, Moon Ki Kim, Sangjae Seo

## Abstract

This study investigates the thermodynamic parameters of 1,300 RNA/DNA hybrid duplexes, including both natural and chemically modified forms, using molecular dynamics (MD) simulations. Modified duplexes consist of phosphorothioate (PS) backbones and 2′-O-methoxyethyl (MOE) modifications, both commonly used in therapeutic oligonucleotides. Hybridization enthalpy and entropy were calculated from MD trajectories using molecular mechanics Poisson-Boltzmann surface area (MMPBSA) and molecular mechanics generalized Born surface area (MMGBSA) approaches. To address discrepancies with experimental data, we established empirical relationships by comparing calculated values with known experimental results of natural hybrid duplexes, then extended these relationships to the entire dataset. The corrected parameters were subsequently used to generate nearest-neighbor (NN) models, allowing for experimentally reliable melting temperature predictions. In this process, MMGBSA demonstrated superior predictive performance with high convergence and consistency for both natural and modified duplexes. Specifically, MMGBSA captured the stabilizing effects of the MOE modification with minimal bias, while MMPBSA exhibited greater variability and limited reliability. These findings highlight the potential of MMGBSA for accurate thermodynamic modeling of both natural and modified nucleic acids, providing a robust framework and experimentally meaningful insights for applications in nucleic acid-based therapeutic design and biotechnology.

## 1. Introduction

Antisense oligonucleotides (ASOs) represent a unique class of modified nucleic acids (NAs), specifically engineered to bind complementary RNA sequences and modulate gene expression. ASOs are gaining significant attention as a therapeutic approach for treating diseases linked to aberrant gene expression, including cancer, genetic disorders, and viral infections. To improve their stability and therapeutic efficacy, ASOs are chemically modified with features such as phosphorothioate (PS) backbones,^1,2^ which replace one of the non-bridging oxygen atoms with sulfur. This modification increases their resistance to nucleases, thereby enhancing their stability in biological environments; however, it reduces the melting temperature, indicating a decrease in binding affinity to the target RNA.^1^ To address this issue, 2′-modifications such as 2′-O-methyl (2′-O-Me) and 2′-O-methoxyethyl (2′-O-MOE) are introduced.^3–5^ In this modification, they are incorporated into the sugar backbone to enhance binding affinity to target RNA and reduce off-target effects,^6,7^ further establishing ASOs as a powerful therapeutic tool for precise gene regulation.

These chemical modifications not only enhance the binding affinity and reduce off-target effects but also significantly impact the structural stability and dynamic properties, which are crucial for their therapeutic function. As a result, extensive research has been conducted in various scientific fields including molecular biology, medicine, and nanotechnology to better understand these properties.^8–10^ In particular, the thermodynamic properties of ASOs—enthalpy (Δ*H*), entropy (Δ*S*), and Gibbs free energy (Δ*G*)—directly influence their stability and provide insights into the molecular interactions and energy changes that occur during processes such as hybridization and conformational transitions. The melting temperature is another key parameter that indicates the temperature at which half of the DNA or RNA strands are in a double-helical state while the other half are single-stranded, making it essential for assessing the hybridization stability of oligonucleotide sequences. Since the melting temperature reflects both sequence-specific interactions and environmental conditions, it is widely employed in applications ranging from genotyping and molecular diagnostics to the design of therapeutics targeting NA sequences.^11–13^

One widely used predictive method for estimating the thermodynamic properties of NAs is the nearest-neighbor (NN) model,^14,15^ which enables calculations based on interactions between adjacent nucleotide pairs. The NN models are valued for their simplicity and effectiveness, providing a straightforward method to estimate melting temperatures and other thermodynamic properties based solely on sequence composition. For this reason, the conventional NN models have long provided essential reference data for understanding NA stability and hybridization properties.^14,16,17^ However, with the rise of chemically modified NAs in therapeutics and advanced research, the need for constructing new NN models that can accurately represent these modifications has grown. Simultaneously, advances in computing power have enabled the use of molecular dynamics (MD) simulations to directly derive thermodynamic properties for these modified structures, paving the way for enhanced precision and adaptability in NA analysis.

MD simulation is one of the most advanced theoretical techniques for exploring molecular systems, as it allows for the time-dependent evolution of molecular motions and samples the conformational space of a system. By numerically solving Newton’s equations of motion, MD simulation provides insights into structural dynamics and interactions, including structural flexibility, binding affinities, and conformational changes, at a resolution difficult to achieve through experimental methods alone. In recent years, numerous studies have utilized MD simulations to examine the hybridization and stability of DNA and RNA duplexes in relation to sequence composition, temperature, and chemical modification.^18–21^ Specifically, techniques such as molecular mechanics Poisson-Boltzmann surface area (MMPBSA), molecular mechanics generalized Born surface area (MMGBSA), and umbrella sampling are frequently applied in MD simulations to calculate binding free energies and assess interaction dynamics of NA duplexes.^22–24^ These simulations are increasingly applied to chemically modified NAs, thus enabling the evaluation of how specific modifications alter duplex stability and molecular affinity. For example, modified NAs, such as locked nucleic acids (LNAs),^25^ peptide nucleic acids (PNAs),^26^ and glycine morpholino oligomers (gMO),^27^ have been extensively studied using MD simulations.^28–31^ This approach through MD simulation has provided a robust framework for the rational design of more effective NA-based therapeutics.

In this study, we performed extensive MD simulations on a dataset of 1,300 RNA/DNA duplexes, including both natural and ASOs. For ASOs, particular attention was given to structures with PS backbones and PS combined with 2′-O-MOE modifications. Thermodynamic properties were calculated using MMPBSA and MMGBSA methods, and to reconcile the differences between the calculated and experimental values, we derived empirical relationships based on experimental data from natural duplexes. These relationships allowed us to correct the calculated thermodynamic parameters. With the corrected thermodynamic properties, the melting temperatures were determined and employed in the construction of an NN model, thereby providing experimentally relevant thermodynamic parameters that improve predictive accuracy for modified oligonucleotide systems. Finally, the obtained NN model was used to predict melting temperatures, which were compared with existing experimental melting temperatures to validate the accuracy of the NN model.

## 2. Materials and Methods

### 2.1. System Preparation

In this study, we generated a dataset by substituting thymine (T) with uracil (U) in the DNA sequence list presented in SantaLucia et al. (1996),^17^ SantaLucia et al. (1998),^14^ and Owczarzy et al. (2004)^16^ to create RNA sequence list. Each RNA sequence was paired with its complementary DNA strand, establishing a collection of RNA/DNA hybrid duplexes. Additionally, RNA/DNA hybrid duplexes from Sugimoto et al. (1995)^32^ were included, resulting in a final dataset of 260 non-redundant sequences. The database includes sequences ranging in length from 4 to 30 base pairs (bp), and all modified structures were generated using a custom script developed with the MDAnalysis Python library.^33,34^

For each of these 260 unique sequences, we constructed five cases of RNA/natural-DNA and RNA/modified-DNA hybrid duplexes, resulting in a total of 1,300 systems: (1) natural RNA/DNA duplexes (RNA/DNA); (2) RNA/DNA duplexes with a PS backbone modification in DNA (RNA/PSDNA); (3) RNA/DNA duplexes with both a PS backbone and 2′-O-MOE modification in DNA (RNA/PSMOE); (4) RNA/DNA duplexes with alternating PS and PSMOE modifications in DNA, starting with PS (RNA/PSDNA-PSMOE); and (5) RNA/DNA duplexes with alternating modifications starting with PSMOE (RNA/PSMOE-PSDNA). Figure 1 illustrates the structures of the PSDNA and PSMOE, providing a visual representation of these modified configurations.

**Figure 1.**
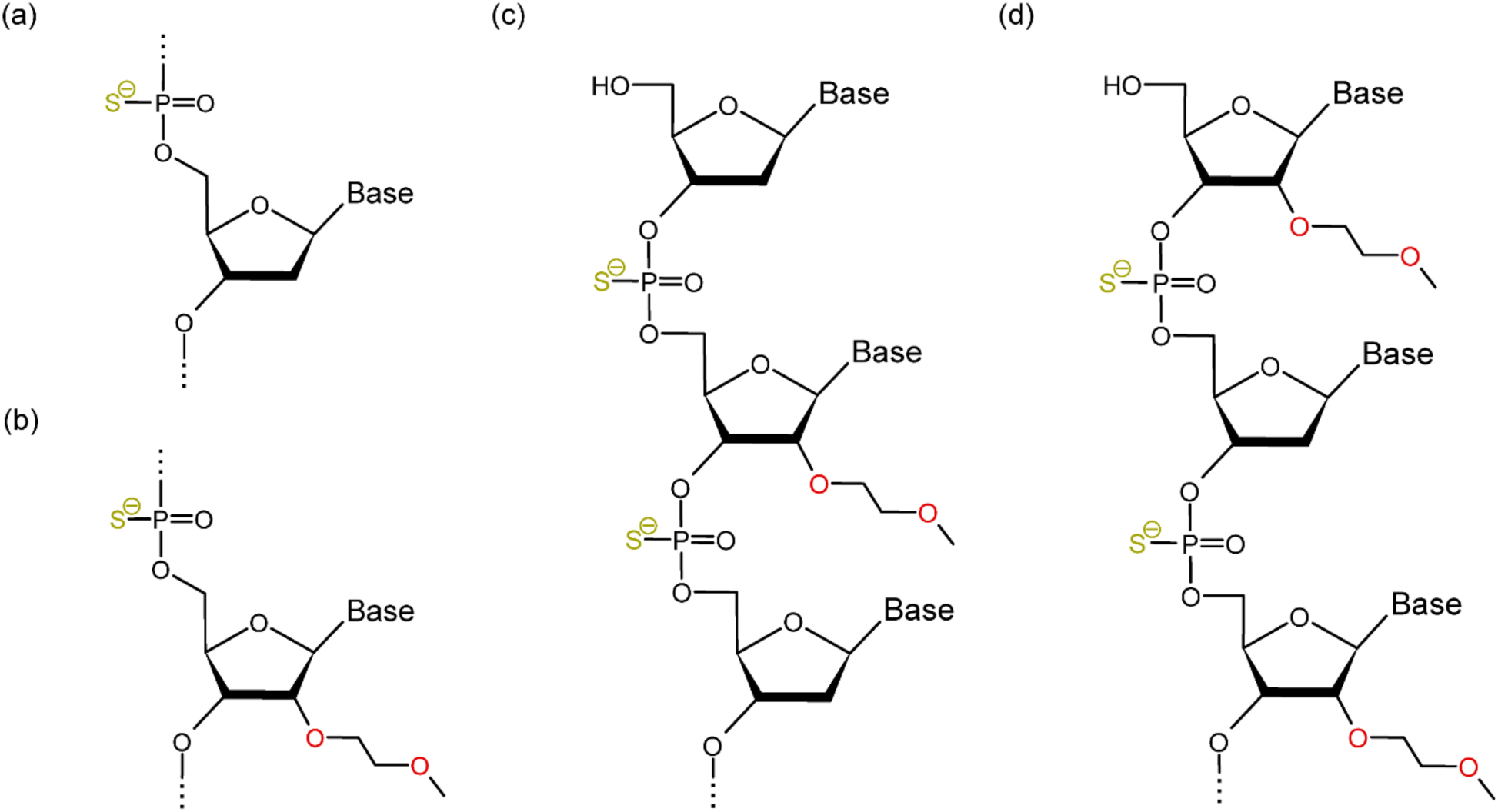
Schematic representation of chemical modifications in nucleotides. (a) Phosphorothioate (PS) backbone modification, where one of the non-bridging oxygen atoms in the phosphate group is replaced with sulfur (RNA/PSDNA). (b) 2′-O-methoxyethyl (MOE) modification, where a methoxyethyl group is attached at the 2′ position of the sugar ring, combined with the PS backbone (RNA/PSMOE). (c) RNA/DNA duplex with alternating PS and PSMOE modifications in DNA, starting with PS (RNA/PSDNA-PSMOE). (d) RNA/DNA duplex with alternating PS and PSMOE modifications in DNA, starting with PSMOE (RNA/PSMOE-PSDNA).

### 2.2. Molecular Dynamics Simulation

The duplex structures were generated as A-type helices using the Nucleic Acid Builder (NAB) tool^35^ and the Chimera.^36^ Each RNA/DNA duplex was constructed by combining the DNA and RNA single strands obtained by removing one strand from the generated duplex structure. The resulting RNA/DNA duplexes were solvated with the explicit TIP3P water model^37^ in a cubic box with a 10 Å edge distance using the xleap module in AmberTools23,^38^ and the systems were neutralized with sodium ions. For the natural structures, the OL15^39^ and OL3^40^ AMBER force fields were applied to DNA and RNA, respectively. For the modified structures, the geometries of PSDNA and PSMOE were optimized using the Hartree-Fock method with the 6-31G* basis set in Gaussian16,^41^ followed by restrained electrostatic potential (RESP) charge fitting^42^ using the Antechamber module^43^ included in AmberTools23. The resulting AMBER files were converted to GROMACS format using ParmEd package,^44^ and all MD simulations were subsequently performed using GROMACS.^45^

In MD simulations, all covalent bonds involving hydrogen atoms were constrained with the SHAKE algorithm^46^ to use a 2 fs time step. Long-range electrostatic interactions were calculated using the particle mesh Ewald (PME) method^47^ with fourth-order cubic interpolation and 0.16 nm grid spacing. A nonbonded cutoff of 10 Å was applied. Energy minimization was performed using the steepest descent method for 50,000 steps until the maximum force was less than 1,000 kJ/mol/nm. After minimization, equilibration was conducted in two stages. The system was first equilibrated under NVT conditions for 200 ps at 300 K using the V-rescale thermostat.^48^ Subsequently, NPT equilibration was performed for 1 ns at 300 K and 1 bar using the V-rescale thermostat and the Parrinello-Rahman barostat.^49^ Throughout the equilibration, harmonic restraints of 1,000 kJ/mol/nm^2^ were applied to the heavy atoms. Finally, a 100 ns production run was carried out at 300 K and 1 bar, employing the Nose-Hoover thermostat^50,51^ and Parrinello-Rahman barostat.

### 2.3. Calculation of Hybridization Enthalpy and Entropy

Hybridization enthalpy was calculated using both MMPBSA and MMGBSA approaches to analyze the MD trajectory. The obtained GROMACS trajectory was converted into AMBER format using the cpptraj module^52^ in AmberTools, and the hybridization enthalpy was subsequently calculated using the MMPBSA.py script.^53^ A total of 1,000 frames were extracted from the trajectory at 100 ps intervals for analysis. The binding free energy was calculated using the following equation:

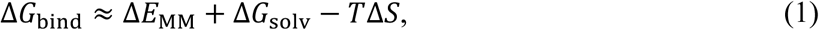

where Δ*E*_MM_ represents the molecular mechanics energy in a vacuum, which includes internal energy, electrostatic interactions, and van der Waals components. Δ*G*_solv_ is the solvation free energy, consisting of polar and non-polar contributions. The polar contribution is calculated using either the Poisson-Boltzmann (PB) or Generalized Born (GB) models, while the non-polar contribution is typically estimated based on the solvent-accessible surface area (SASA). Finally, *T*Δ*S* corresponds to the entropy contribution.

For the MMGBSA calculations, we used the igb = 5 setting, which corresponds to the OBC modified GB model developed by Onufriev and co-workers.^54^ This model is widely considered suitable for nucleic acids due to its improved accuracy in accounting for solvation effects. Additionally, PB calculations were performed, with the dielectric constants for the solvent and solute set to 80.0 and 1.0, respectively. The ionic strength was set to 0.1 M. For entropy calculations, the normal mode analysis (NMA) method implemented in MMPBSA.py was employed.

The enthalpy and entropy values derived from MD simulations generally align with the trends observed in experimental data, as reported in previous studies.^19,20^ However, direct numerical comparison with experimental data is often difficult due to inherent differences in simulation and experimental conditions, such as force fields and solvent models. To overcome this challenge, we aimed to establish an empirical relationship between the MD-derived and experimental thermodynamic values for natural RNA/DNA complexes with known experimental data. For this purpose, we constructed a dataset that incorporates both the enthalpy (Δ*H*) and entropy (Δ*S*) values as a pair of dependent variables. Using the SciPy library,^55^ a least-squares fitting was applied to the following linear relationship:

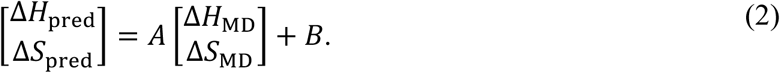

Here, Δ*H*_MD_ and Δ*S*_MD_ represent the values obtained from the MD simulations, and Δ*H*_pred_ and Δ*S*_pred_ are the predicted values fitted to minimize the discrepancy with respect to the experimental data. Therefore, the parameters *A* and *B* were determined by minimizing the sum of squared residuals between the experimental and predicted values. The derived relationship was subsequently extended to RNA/modified-DNA complexes, which allowed for the prediction of thermodynamic values that are consistent with experimental findings.

### 2.4. Estimation of Melting Temperature and Nearest-Neighbor Models

In this study, the corrected enthalpy and entropy values, obtained from MD simulations and subsequently fitted to experimental data, were used to calculate the Gibbs free energy and melting temperature of the RNA/DNA hybrid duplexes. The relationship between Gibbs free energy (Δ*G_T_*°) and the enthalpy and entropy is given by the following equation:

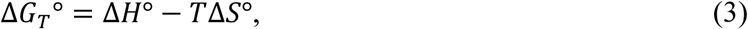

where *T* is the temperature in Kelvin, Δ*H*° is the enthalpy in units of cal/mol, and Δ*S*° is the entropy in units of cal/K/mol (commonly referred to as entropy units, e.u.). Using this equation, we calculated the free energy at various temperatures to assess the thermodynamic stability of the hybrid duplexes.

For the melting temperature (*T*_m_), we used the following equation:

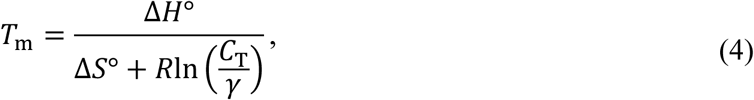

where *C*_T_ is the species concentration, *R* is the gas constant, and γ accounts for the duplex symmetry. In the case of non-self-complementary duplexes, γ equals 4, while for self-complementary duplexes, γ equals 1. As the focus of this study is on RNA/DNA hybrid duplexes, which are non-self-complementary by nature, we used γ = 4 in all calculations. This formula allows us to determine the melting temperature based on the thermodynamic properties of the duplex.

To construct the NN model for RNA/DNA hybrid duplexes, we employed the VarGibbs tool,^56^ which optimizes NN parameters using a multidimensional minimization approach. In VarGibbs, the algorithm minimizes the objective function, χ^2^, where the goal is to minimize the difference between measured melting temperatures and those predicted by the NN model.

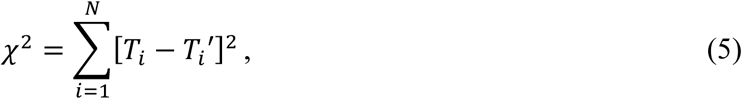

where *N* is the number of sequences, *T_i_* is the measured melting temperature for the *i* th sequence, and *T_i_*^′^ is the corresponding predicted temperature. The detailed procedure for parameter optimization is described in ref. 56.

## 3. Results and Discussion

### 3.1. Hybridization Enthalpy and Entropy of the Natural RNA/DNA Duplexes from MD Simulations

In this study, 100 ns MD simulations were performed on 1,300 systems of the natural and modified RNA/DNA duplexes. Enthalpy and entropy values were derived from the MD trajectories, and all the results were shown in Table S1. To compare these values with experimental data, we specifically examined the thermodynamic parameters for 64 natural RNA/DNA sequences experimentally revealed by Sugimoto et al. (1995).^32^ Figure 2 illustrates the correlation between the calculated and experimental values of hybridization enthalpy and entropy of these sequences.

**Figure 2.**
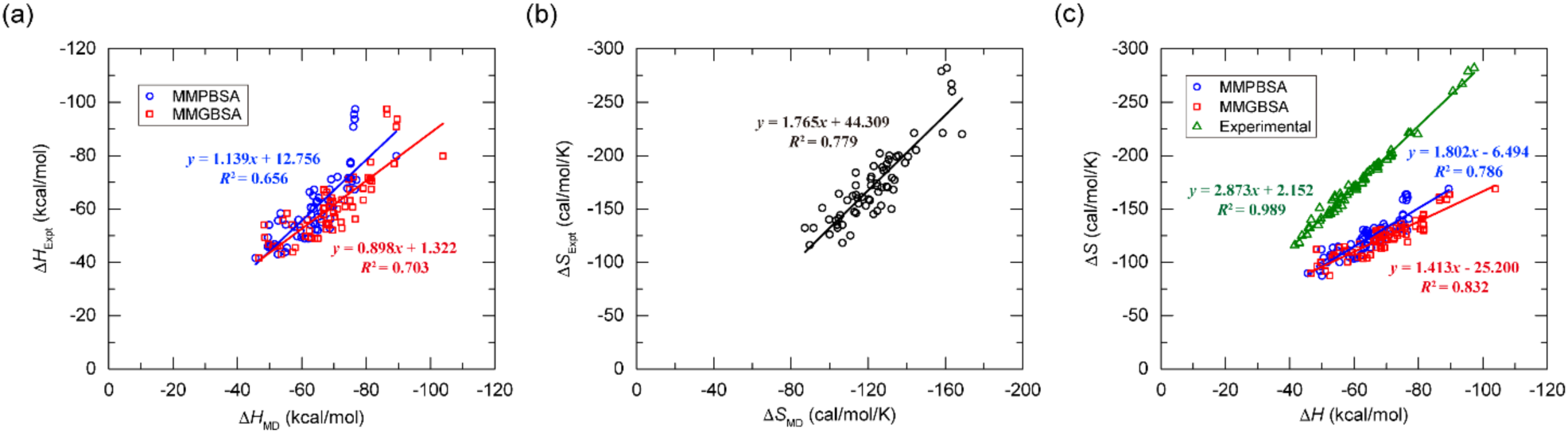
Correlation of thermodynamic parameters for 64 natural RNA/DNA duplexes with known experimental values. (a) Correlation of hybridization enthalpy (Δ*H*) values between MD-derived (MMPBSA and MMGBSA) and experimental data. (b) Correlation of hybridization entropy (Δ*S*) values between MD-derived and experimental data. (c) Enthalpy-entropy compensation, showing the linear relationship between enthalpy and entropy values for MMPBSA, MMGBSA, and experimental data.

The correlation of hybridization enthalpy values is shown in Figure 2a. MMPBSA showed an *R*^2^ of 0.66, where *R* is the Pearson correlation coefficient, with a slope of 1.14 and an offset of 12.76. This suggests that, while there is moderate correlation, MMPBSA tends to overestimate the enthalpy values compared to the experimental data. An absolute error was 6.85 kcal/mol, with a relative error of 11.56%. Here, the absolute error, 〈|Δ*H*_MD_−Δ*H*_Expt_|〉, is the average absolute difference between the calculated and experimental values, while the relative error, 〈|Δ*H*_MD_−Δ*H*_Expt_|/|Δ*H*_Expt_|〉, measures the magnitude of the error relative to the experimental values. In contrast, MMGBSA had a higher *R*^2^ of 0.70, with a slope of 0.90 and an offset of 1.32, indicating that MMGBSA provided better alignment with the experimental data. However, the absolute error for MMGBSA was 9.51 kcal/mol, which is larger than that of MMPBSA, with the relative error of 16.38%. These differences suggest that, while MMGBSA exhibited a stronger overall correlation, MMPBSA yielded more accurate absolute enthalpy values, albeit with a systematic overestimation. Both methods tended to overestimate the experimental enthalpy values, which could be attributed to inherent differences between MD and experimental setups, such as variations in force fields and solvent models. In addition, assumptions in the MMGBSA and MMPBSA methods, particularly regarding solvation effects, may contribute to these discrepancies. In Figure 2b, the correlation between entropy values from the NMA method included in MMPBSA.py and experimental data is shown. Despite the relatively high *R*^2^ value of 0.79, a slope of 1.76 and an offset of 44.31 suggested a significant shift from the experimental trend, especially with the absolute error of 49.82 cal/mol/K and the relative error of 27.80%. This implies that while the general trends are captured, the absolute entropy values are significantly overestimated, limiting their direct applicability in practical examples.

Figure 2c presents the linear relationship between the hybridization enthalpy and entropy values, illustrating enthalpy-entropy compensation. The experimental data (green triangles) showed strong linearity (*R*^2^ = 0.989), indicative of near-perfect compensation between these two thermodynamic quantities. The MMPBSA and MMGBSA data also showed reasonable correlation, *R*^2^ of 0.79 and 0.83, respectively. However, the slopes for MMPBSA (1.80) and MMGBSA (1.41), along with their offsets of -6.49 and -25.20, deviated significantly from the experimental slope of 2.87 and offset of 2.15. These deviations further highlight the tendency of MD-derived results to diverge from experimental thermodynamic parameters, particularly in absolute values. In the context of enthalpy-entropy compensation, it is clear that while MD methods were able to capture the relative trends, the discrepancies arise due to differences in force fields and other simulation-specific factors. To reduce these discrepancies and provide thermodynamically meaningful results, we performed a least-squares fit, establishing empirical relationships between the MD-derived and experimental values. This approach can help address the inherent differences between simulations and experimental data, ultimately leading to more experimentally relevant results for the RNA/DNA complexes.

### 3.2. Empirical Fit of Thermodynamic Parameters to Experimental Data

To refine the accuracy of the thermodynamic parameters derived from the MD simulations, fitting models were developed based on the experimental data. By comparing the hybridization enthalpy (Δ*H*) and entropy (Δ*S*) values obtained from the MMPBSA and MMGBSA methods with experimentally measured values, we established linear relationships aimed at bridging the gap between MD-derived predictions and experimentally measured values. The resulting fitting models for the MMPBSA and MMGBSA methods were as follows:

For MMPBSA,

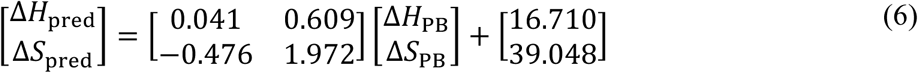

For MMGBSA,

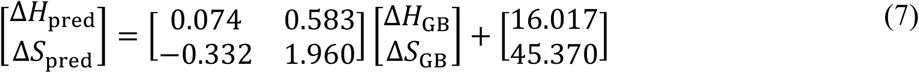

In both models, the slope coefficients represent the degree of scaling applied to both Δ*H* and Δ*S*, correcting the MD-derived values to better match experimental data. The positive offsets suggest additional corrections to account for consistent baseline deviations between simulations and experiments. The off-diagonal coefficients indicate the interaction between Δ*H* and Δ*S*. In MMPBSA, the values (0.609 and -0.476) suggest a moderate degree of coupling between Δ*H* and Δ*S*, whereas in MMGBSA, the off-diagonal coefficients (0.583 and -0.332) are slightly smaller in magnitude, indicating weaker coupling between Δ*H* and Δ*S* compared to MMPBSA. Despite these subtle differences, the interaction between Δ*H* and Δ*S* contributes notably to the final corrections in both cases, reflecting the complex interplay between these thermodynamic parameters in the fitting process.

By applying these corrections, the alignment between the predicted and experimental values was significantly improved. The effectiveness of the fitting models was demonstrated in Figure 3. In Figure 3a, the *R*^2^ for MMPBSA increased from 0.786 to 0.99, with the slope and offset shifted from 1.80 and -6.49 to 2.81 and -1.64, respectively. For MMGBSA, a similar trend was observed in Figure 3b, where the *R*^2^ value increased from 0.83 to 0.99 following the application of the fitting formula, indicating near-perfect correlation with the experimental data. The slope and offset shifted from 1.41 and -25.20 to 2.81 and -1.88, closely matching the experimental slope of 2.87 and offset of 2.15. Notably, the application of the fitting models significantly reduced both the absolute and relative errors for enthalpy and entropy. For MMPBSA, the absolute error in enthalpy decreased from 6.85 to 4.15 kcal/mol, with the relative error dropping from 11.56% to 6.78%. Similarly, for MMGBSA, the absolute error in enthalpy was reduced from 9.51 to 4.16 kcal/mol, and the relative error decreased from 16.38% to 6.79%. Entropy values also showed substantial improvements; the absolute error in entropy decreased from 49.82 to 13.12 cal/mol/K for MMPBSA and 13.14 cal/mol/K for MMGBSA, with corresponding reductions in the relative error from 27.80% to 7.64% in both cases. These results suggest that both the MMPBSA and MMGBSA methods, when combined with the appropriate post-simulation adjustments, can accurately reproduce experimental thermodynamic parameters for natural RNA/DNA duplexes. However, while these results demonstrate the reliability of the fitting models for 64 natural duplexes examined in this study, additional validation will be necessary for broader applications to other NA systems, particularly those involving modified NAs.

**Figure 3.**
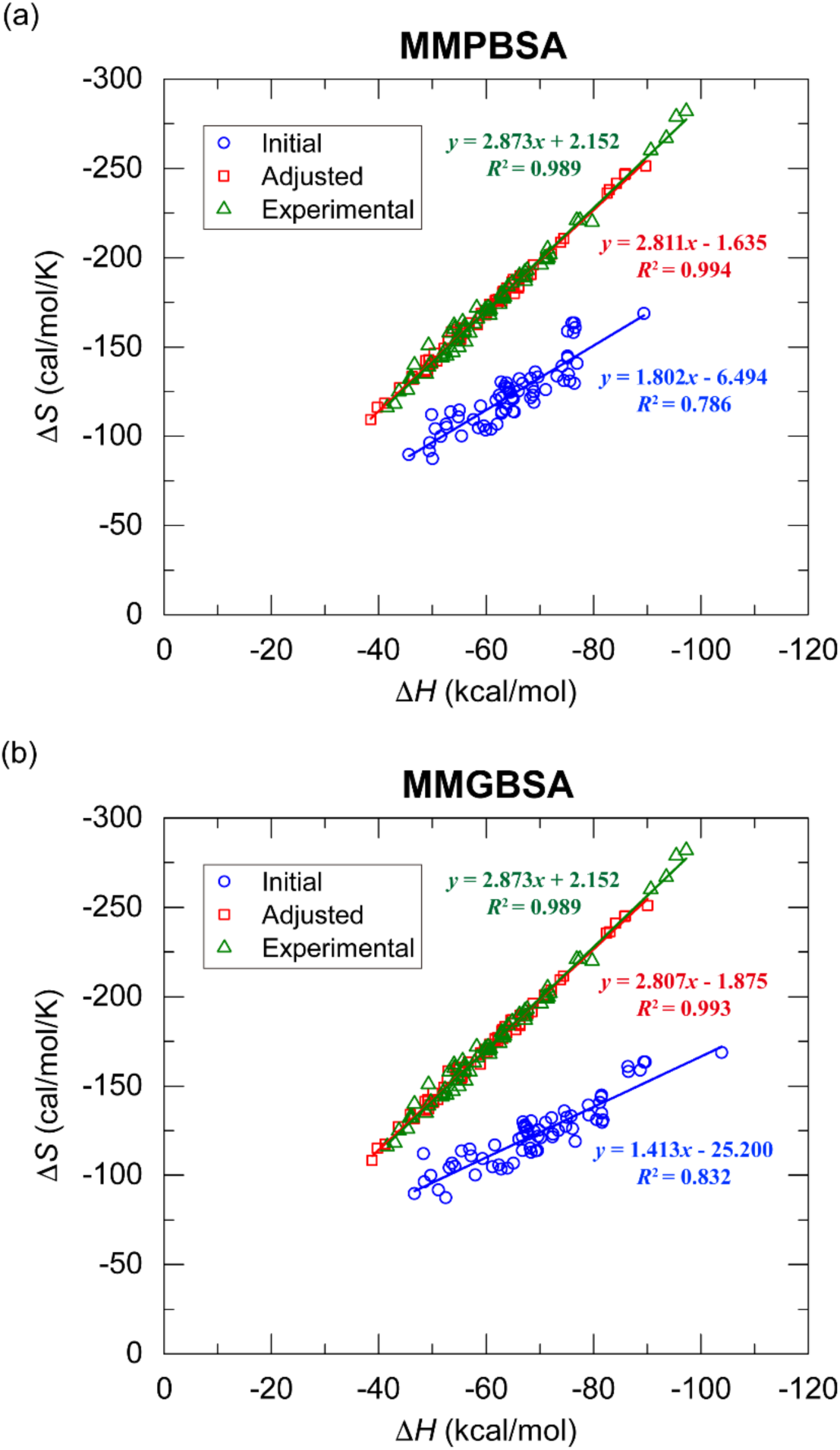
Thermodynamic parameters of 64 natural RNA/DNA duplexes adjusted using empirical relationships. (a) MMPBSA and (b) MMGBSA results. Initial values (blue circles) represent unadjusted MD-derived data, adjusted values (red squares) represent data corrected by the empirical formula, and experimental values (green triangles) are from experimental measurements.

### 3.3. Correction of MD-Derived Thermodynamic Parameters

In this subsection, the previously derived relationships (eqs 6 and 7) were extended beyond the initial 64 natural RNA/DNA duplexes to a total of 260 natural RNA/DNA duplexes, and further applied to four types of modified duplexes: RNA/PSDNA, RNA/PSMOE, RNA/PSDNA-PSMOE, and RNA/PSMOE-PSDNA. It is important to note that RNA/PSDNA-PSMOE and RNA/PSMOE-PSDNA share identical repeating PSDNA and PSMOE structures, differing only in the order of modifications. Given this structural similarity and the fact that the NN model is generated identically for both cases, these two cases were combined into a single dataset (RNA/PS+PSMOE), resulting in a total of 520 duplexes.

The results of applying the fitting equations derived from the natural RNA/DNA duplexes to the entire dataset are shown in Figure 4. A notable observation across all natural/modified duplexes is the significant improvement in the *R*^2^ values for both MMPBSA and MMGBSA, which increased to the range of 0.997-0.999 after fitting. Prior to correction, the MMPBSA *R*^2^ values for RNA/DNA, RNA/PSDNA, RNA/PSMOE, and RNA/PS+PSMOE were measured at 0.792, 0.767, 0.822, and 0.781, respectively, all of which were lower than the MMGBSA values of 0.951, 0.949, 0.962, and 0.953. However, upon applying the fitting procedure, these discrepancies were effectively resolved, demonstrating the robustness of the correction formula in aligning both methods with the experimental data. This result underscores the ability of the derived relationships, initially obtained from 64 natural duplexes, to generalize not only to the expanded set of 260 natural RNA/DNA duplexes (Figures 4a and 4b) but also to the 1,040 RNA/modified-DNA duplexes (Figures 4c-4h). The successful application of the fitting models across all cases highlights its capacity to accurately describe the enthalpy-entropy compensation in both natural and modified systems.

**Figure 4.**
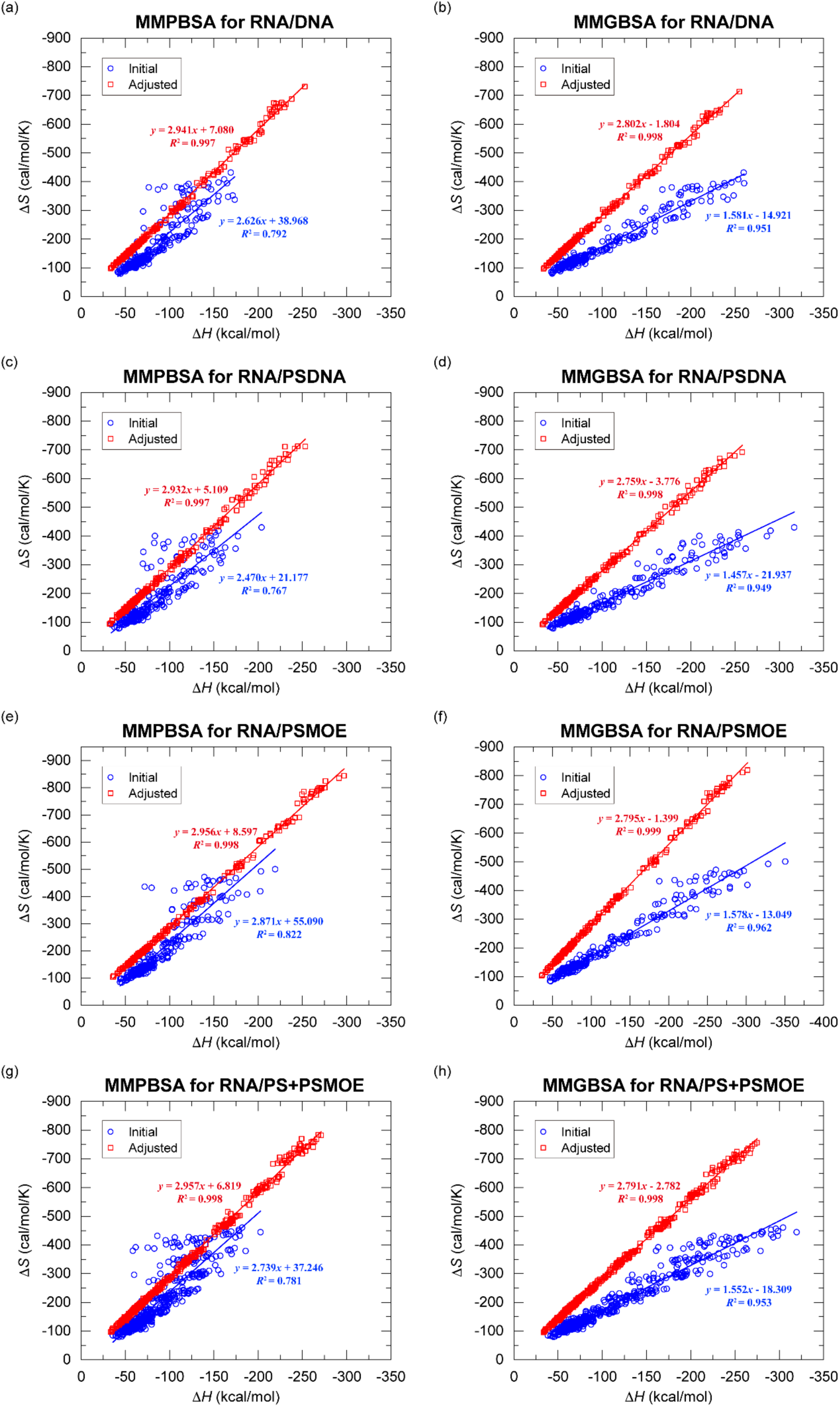
Enthalpy-entropy compensation for initial and adjusted thermodynamic parameters across different RNA/DNA duplexes. (a)-(h) show the Δ*H*-Δ*S* correlations with initial MD-derived values (blue circles) and adjusted values (red squares) corrected using empirical relationships. Each plot includes linear fits with the corresponding regression equations and *R*^2^ values. The panels represent: (a) MMPBSA for RNA/DNA, (b) MMGBSA for RNA/DNA, (c) MMPBSA for RNA/PSDNA, (d) MMGBSA for RNA/PSDNA, (e) MMPBSA for RNA/PSMOE, (f) MMGBSA for RNA/PSMOE, (g) MMPBSA for RNA/PS+PSMOE, and (h) MMGBSA for RNA/PS+PSMOE.

Based on these corrected enthalpy and entropy values, we then calculated the Gibbs free energy and melting temperatures for all RNA/DNA duplexes using eqs 3 and 4. The complete sets of the corrected thermodynamic parameters are provided in Supporting Information (Tables S2-S5). In the next subsection, we constructed the corresponding NN models based on these results and predicted the enthalpy and entropy values. To validate the accuracy of our entire workflow, we further compared the predicted melting temperatures from the NN models with the experimentally measured melting temperatures of several RNA/modified-DNA duplexes.

### 3.4. Development and Assessment of Nearest-Neighbor Models

Based on the corrected thermodynamic parameters (Tables S2-S5), we employed the VarGibbs tool^56^ to generate the NN models, with the results presented in Tables 1-4. One key observation is the more consistent and reasonable convergence of the MMGBSA values for all duplexes, whereas MMPBSA displayed a wider range in the predicted values. For example, in Table 1, the rGU/dCA sequence yields MMPBSA-derived values of Δ*H* = 6.019 kcal/mol and Δ*S* = 22.192 cal/mol/K, and the corresponding MMGBSA-derived values are Δ*H* = -2.765 kcal/mol and Δ*S* = -5.005 cal/mol/K. In contrast, the rUC/dAG sequence shows a much larger discrepancy in MMPBSA values (Δ*H* = -55.940 kcal/mol and Δ*S* = -171.688 cal/mol/K) compared to MMGBSA (Δ*H* = - 20.135 kcal/mol and Δ*S* = -59.740 cal/mol/K). This significant range in MMPBSA-derived enthalpy (ranging from -55.940 to 6.019 kcal/mol) and entropy (ranging from -171.688 to 22.192 cal/mol/K) suggests problematic convergence, which is consistently observed across different modified duplexes. Despite our efforts to address this issue, including adjusting the characteristic length in the downhill simplex method from 1 to 0.01 to limit the search range of the NN parameters, MMPBSA-derived values still exhibited substantial variability.

**Table 1.** Nearest-Neighbor Thermodynamic Parameters for Natural RNA/DNA Duplexes.

| Propagation Sequence | MMPBSA |  | MMGBSA |  |
| --- | --- | --- | --- | --- |
| | $\Delta H^\circ$ | $\Delta S^\circ$ | $\Delta H^\circ$ | $\Delta S^\circ$ |
|  | (kcal/mol) | (cal/mol/K) | (kcal/mol) | (cal/mol/K) |
| rAA/dTT | -12.236 | -36.300 | -9.177 | -26.829 |
| rAU/dTA | 3.294 | 11.905 | -3.683 | -9.928 |
| rAG/dTC | -17.006 | -49.972 | -14.141 | -40.854 |
| rAC/dTG | -10.756 | -29.223 | -10.592 | -29.163 |
| rUA/dAT | -18.341 | -55.154 | -9.161 | -26.622 |
| rUU/dAA | -26.644 | -83.120 | -11.066 | -33.188 |
| rUG/dAC | -30.594 | -92.543 | -15.597 | -44.868 |
| rUC/dAG | -55.940 | -171.688 | -20.135 | -59.740 |
| rGA/dCT | -17.010 | -51.058 | -10.611 | -30.532 |
| rGU/dCA | 6.019 | 22.192 | -2.765 | -5.005 |
| rGG/dCC | -11.963 | -31.961 | -11.362 | -29.464 |
| rGC/dCG | -8.332 | -20.515 | -10.802 | -28.878 |
| rCA/dGT | -18.797 | -57.060 | -11.571 | -33.046 |
| rCU/dGA | 5.456 | 19.116 | -4.408 | -11.399 |
| rCG/dGC | -13.567 | -38.480 | -12.430 | -34.503 |
| rCC/dGG | -8.070 | -19.702 | -8.159 | -19.940 |

**Table 2.** Nearest-Neighbor Thermodynamic Parameters for RNA/PSDNA Duplexes.

| Propagation Sequence<sup>a</sup> | MMPBSA |  | MMGBSA |  |
| --- | --- | --- | --- | --- |
| | $\Delta H^\circ$ | $\Delta S^\circ$ | $\Delta H^\circ$ | $\Delta S^\circ$ |
|  | (kcal/mol) | (cal/mol/K) | (kcal/mol) | (cal/mol/K) |
| rAA/dT'T' | -6.091 | -16.675 | -8.150 | -23.342 |
| rAU/dT'A' | -6.383 | -18.981 | -8.865 | -26.104 |
| rAG/dT'C' | -7.104 | -18.350 | -7.559 | -19.385 |
| rAC/dT'G' | -3.533 | -6.880 | -8.798 | -23.663 |
| rUA/dA'T' | -20.848 | -63.795 | -11.029 | -32.379 |
| rUU/dA'A' | -38.461 | -121.043 | -15.313 | -46.085 |
| rUG/dA'C' | -8.095 | -22.343 | -7.848 | -20.841 |
| rUC/dA'G' | -31.017 | -92.759 | -11.094 | -30.628 |
| rGA/dC'T' | -29.441 | -89.887 | -14.074 | -40.292 |
| rGU/dC'A' | -2.984 | -6.345 | -6.746 | -17.463 |
| rGG/dC'C' | -16.651 | -46.982 | -13.077 | -34.563 |
| rGC/dC'G' | -6.219 | -13.504 | -8.691 | -21.702 |
| rCA/dG'T' | -23.283 | -70.976 | -12.038 | -34.242 |
| rCU/dG'A' | -3.254 | -9.235 | -4.740 | -12.973 |
| rCG/dG'C' | -14.337 | -41.409 | -13.202 | -36.854 |
| rCC/dG'G' | -7.841 | -19.454 | -8.421 | -21.186 |

<sup>a</sup> The apostrophe in sequence represents the PS backbone modification in DNA strand.

**Table 3.** Nearest-Neighbor Thermodynamic Parameters for RNA/PSMOE Duplexes.

| Propagation<br>Sequence<sup>a</sup> | MMPBSA |  | MMGBSA |  |
| --- | --- | --- | --- | --- |
| | $\Delta H^\circ$ | $\Delta S^\circ$ | $\Delta H^\circ$ | $\Delta S^\circ$ |
|  | (kcal/mol) | (cal/mol/K) | (kcal/mol) | (cal/mol/K) |
| rAA/dT*T* | -13.197 | -38.522 | -10.197 | -29.334 |
| rAU/dT*A* | 0.748 | 5.169 | -7.178 | -19.883 |
| rAG/dT*C* | -13.836 | -40.640 | -12.951 | -37.043 |
| rAC/dT*G* | 1.174 | 8.139 | -8.103 | -21.419 |
| rUA/dA*T* | -36.988 | -114.552 | -14.766 | -44.017 |
| rUU/dA*A* | -27.926 | -87.554 | -11.969 | -35.803 |
| rUG/dA*C* | -30.144 | -92.846 | -14.672 | -42.225 |
| rUC/dA*G* | -54.601 | -167.433 | -16.967 | -50.091 |
| rGA/dC*T* | -34.886 | -104.971 | -16.299 | -46.473 |
| rGU/dC*A* | 7.400 | 27.243 | -5.001 | -11.724 |
| rGG/dC*C* | -18.257 | -51.179 | -14.229 | -37.762 |
| rGC/dC*G* | -8.251 | -19.359 | -12.111 | -31.949 |
| rCA/dG*T* | -29.389 | -88.678 | -14.538 | -40.775 |
| rCU/dG*A* | -2.458 | -3.483 | -7.767 | -20.284 |
| rCG/dG*C* | -13.357 | -37.507 | -11.354 | -30.247 |
| rCC/dG*G* | -11.177 | -29.053 | -11.102 | -28.826 |

<sup>a</sup> The asterisk in sequence represents the PS + 2'-O-MOE modification in DNA strand.

**Table 4.**
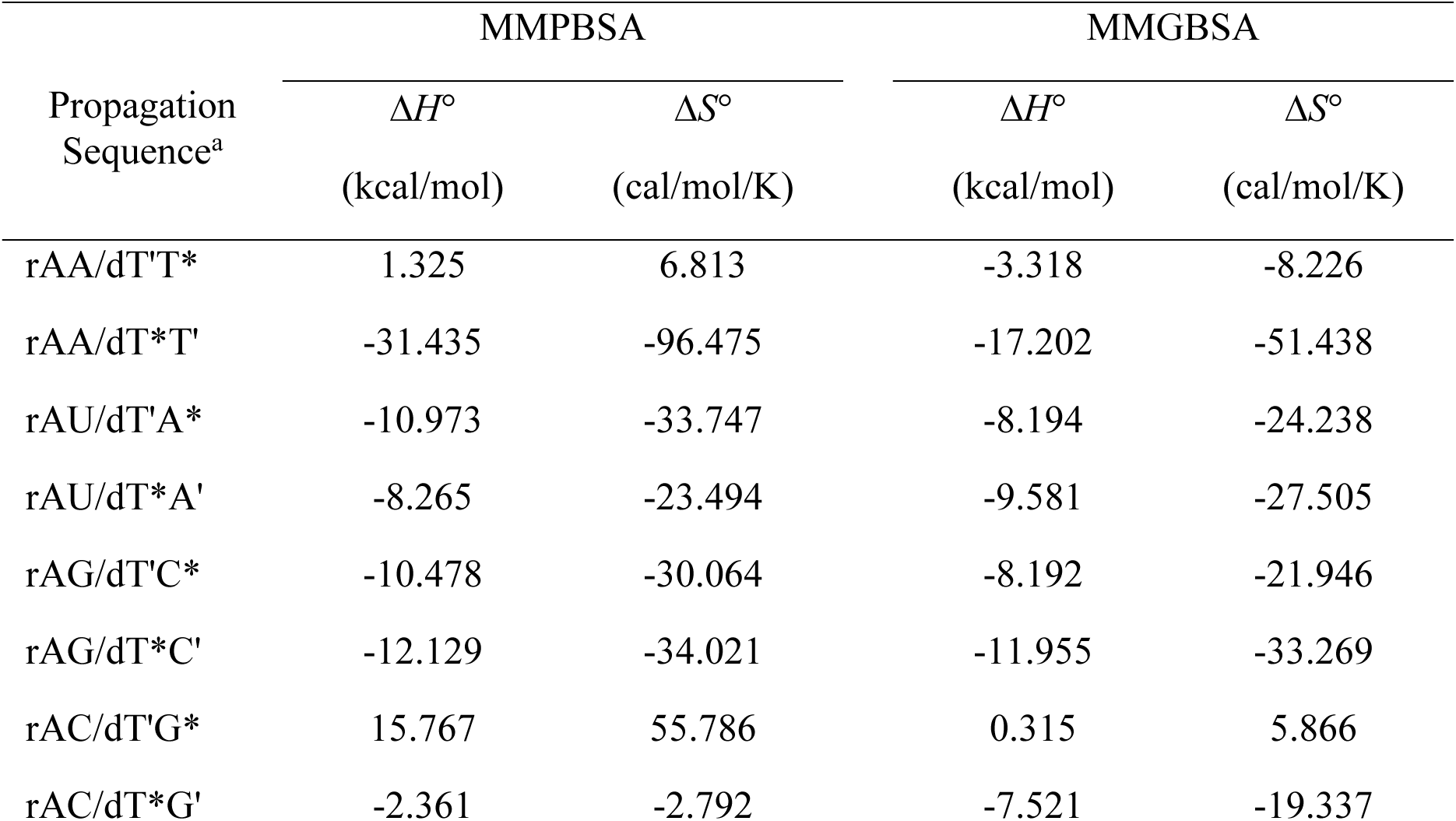
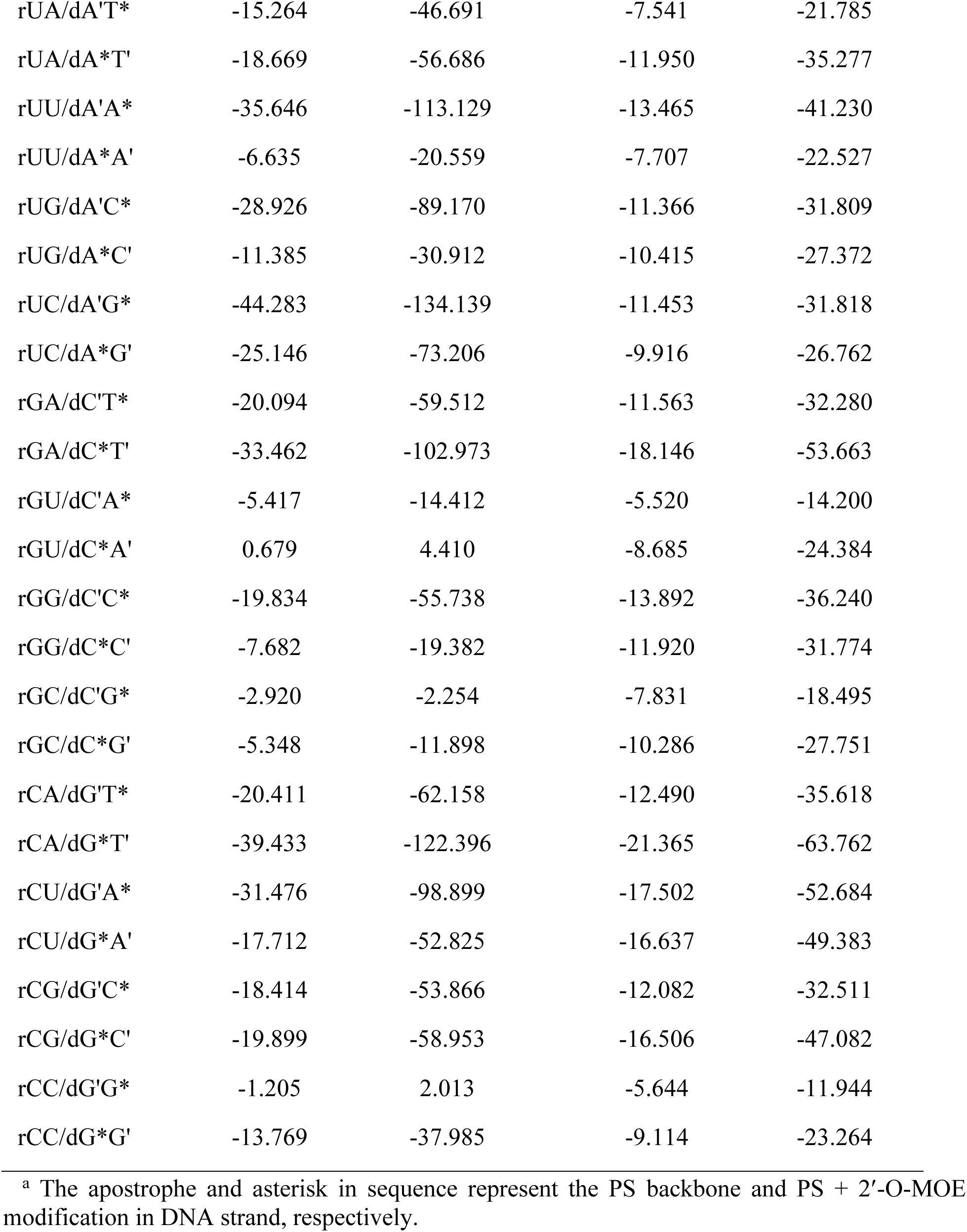
Nearest-Neighbor Thermodynamic Parameters for RNA/PS+PSMOE Duplexes.

This variability was also reflected in the predictive accuracy of the melting temperatures. Figure 5 illustrates the correlation between the melting temperatures predicted by the NN models and those calculated from the MD simulations. Across all RNA/DNA duplexes, MMGBSA consistently outperformed MMPBSA, as indicated by both the slope and *R*^2^ values. For example, in the natural RNA/DNA duplexes, MMGBSA achieved a slope of 0.943 and an *R*^2^ of 0.944, compared to MMPBSA’s slope of 0.829 and *R*^2^ of 0.835. This trend was observed across the RNA/modified-DNA duplexes as well, with MMGBSA yielding slopes of 0.947, 0.956, and 0.949 for RNA/PSDNA, RNA/PSMOE, and RNA/PS+PSMOE, respectively, along with corresponding *R*^2^ values of 0.951, 0.957, and 0.951. In comparison, MMPBSA showed significantly lower slopes (ranging from 0.837 to 0.847) and *R*^2^ values (ranging from 0.842 to 0.851) across these modified duplexes. The fact that MMGBSA produced slopes close to 1 highlights its minimal bias in predicting melting temperatures, and the high *R*^2^ values indicate a strong linear relationship between the predicted and MD-calculated values. These results reinforce the robustness of the MMGBSA method, particularly in handling both natural and modified duplexes. Additionally, the absolute error in the predicted melting temperatures was consistently lower for MMGBSA across all cases. For RNA/DNA duplexes, the absolute error with MMGBSA was 2.816°C, compared to 4.131°C with MMPBSA. Similarly, MMGBSA outperformed MMPBSA for RNA/PSDNA (2.963°C vs. 4.322°C), RNA/PSMOE (2.242°C vs. 3.298°C), and RNA/PS+PSMOE (2.698°C vs. 3.959°C). These results further confirm the superior predictive performance of MMGBSA in estimating melting temperatures using NN model.

**Figure 5.**
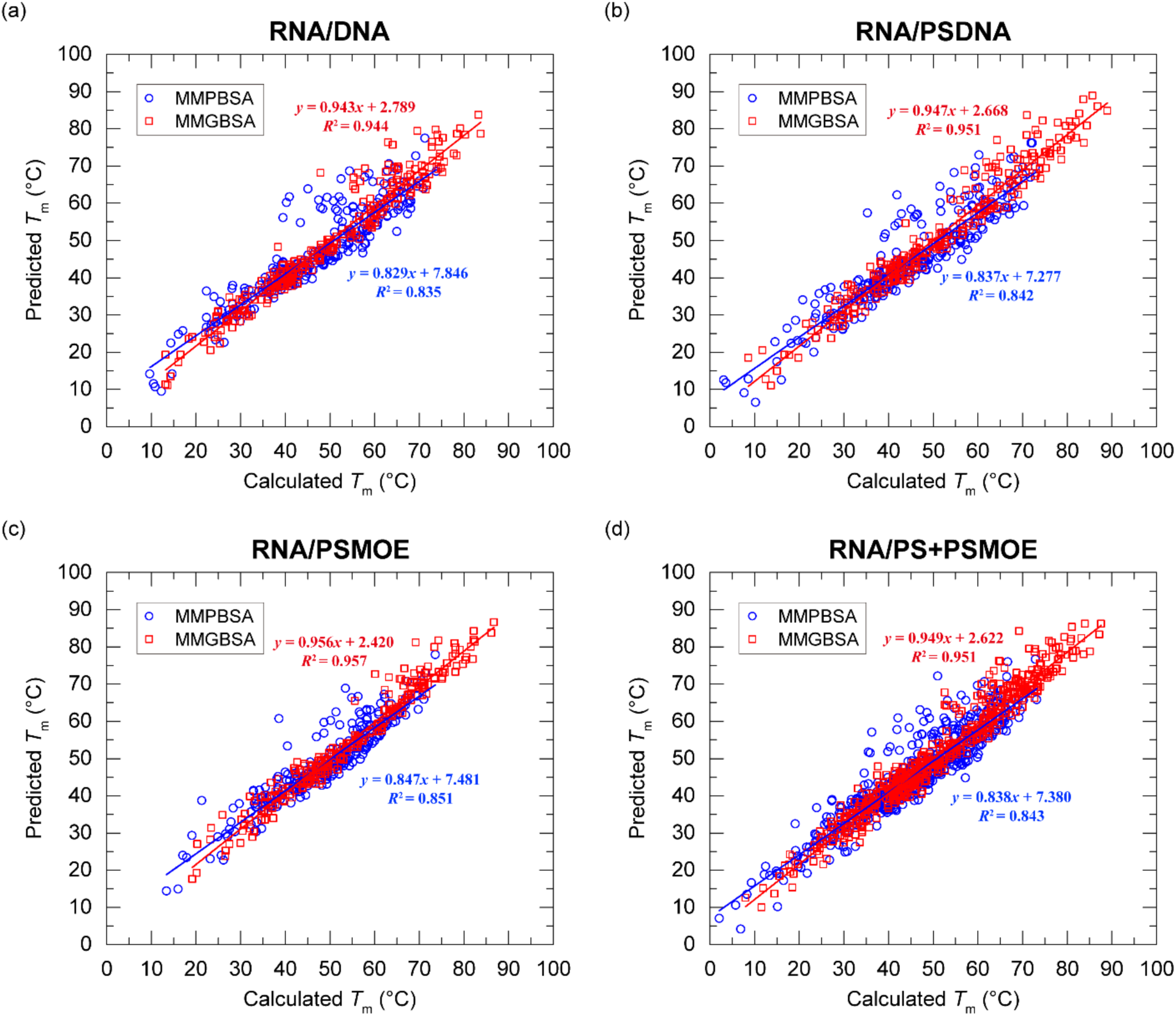
Comparison of predicted and calculated melting temperatures (*T*_m_) for the natural and modified RNA/DNA duplexes. (a)-(d) show the correlation between predicted *T*_m_ values obtained from MMPBSA (blue circles) and MMGBSA (red squares) against calculated *T*_m_ values for each duplex type: (a) RNA/DNA, (b) RNA/PSDNA, (c) RNA/PSMOE, and (d) RNA/PS+PSMOE. Each plot includes linear fits for MMPBSA and MMGBSA data with the corresponding regression equations and *R*^2^ values, illustrating the predictive accuracy of each method across different duplex configurations.

To further assess the reliability of the obtained NN models, we compared the predicted melting temperatures with experimentally determined values from several studies.^57,58^ In the literature by Masaki et al. (2018),^58^ the experimental melting temperature of the PS-d(^m^C_MOE_T_MOE_G_MOE_^m^C_MOE_T_MOE_AG^m^C^m^CT^m^CTGGAT_MOE_T_MOE_T_MOE_G_MOE_A_MOE_) sequence was 72°C. For this sequence, the MMGBSA-derived NN models predicted a melting temperature of 71.580°C, showing strong alignment with the experimental value. Another sequence, PS-d(TCCCGCCTGTGACATGCATT), which features the only PS backbone modification, the experimental melting temperature was 62°C, as reported in Altmann et al. (1996).^57^ However, the MMGBSA-derived NN models overestimated the melting temperature to 74.614°C. This overestimation reflects the limitations in our approach using MD simulations in capturing the destabilizing effects of the PS backbone, which is known to reduce melting temperature by altering the backbone’s flexibility and electrostatic interactions.^59^ Force fields of MD simulation may not fully represent the unique solvation and electrostatic properties of the PS modification, leading to an overestimation of duplex stability in PS backbone structures.

In contrast, PS-d(T_MOE_C_MOE_C_MOE_C_MOE_G_MOE_C_MOE_CTGTGACAT_MOE_G_MOE_C_MOE_A_MOE_T_MOE_T_MOE_), also from Altmann et al. (1996),^57^ had an experimental melting temperature of 76°C. The MMGBSA-derived NN model prediction of 74.764°C closely matched the experimental value, highlighting MMGBSA’s ability to account for the stabilizing influence of the MOE modification. Indeed, the 2′-O-MOE modification increases the melting temperature by further stabilizing the A-form conformation of the ribose backbone, primarily by reinforcing the C3′-endo sugar pucker.^60^ This stabilizing effect of the MOE modification is likely better represented in current MD force fields, which could explain the close approximation of the two MMGBSA-predicted melting temperatures in the RNA/PS+PSMOE duplexes. While the destabilizing effect of the PS backbone might not be fully captured, the prominent stabilizing influence of the MOE modification appears to be well-modeled in the NN prediction.

Given the convergence issue in the MMPBSA-derived NN models, their predictions exhibited significant discrepancies. For the first sequence, the NN models underestimated the melting temperature to 61.460°C, and for the second and third sequences (PS and PS+PSMOE), it predicted values of 64.280°C and 65.653°C, respectively. Although the NN prediction for the PS backbone duplex (64.280°C) seemed close to the experimental value (62°C), this result likely stems from the MMPBSA-derived NN model’s general tendency to underestimate melting temperatures, rather than a true reflection of the PS backbone’s destabilizing effect.

In addition, some of the discrepancies in the PB/GB-predicted melting temperature values can be attributed to differences between the experimental structures and the models used in the MD simulations. For example, the first experimental sequences include modifications such as 5-methylcytosine (5mC or ^m^C), which were not accounted for in our simulations where cytosine was modeled instead. These mismatches in sequence composition, along with other experimental conditions not fully captured in the simulations, likely contributed to the deviations observed between the predicted and experimental melting temperatures. Nonetheless, the NN model based on MMGBSA demonstrated highly accurate predictions for the PS+PSMOE modifications, with predicted melting temperatures of 71.580°C and 74.764°C closely matching the experimental values of 72°C and 76°C. These results of our approach imply that, as experimental conditions are more precisely incorporated into MD simulations, predictions could become even more reliable for a wider range of modified duplexes.

## 4. Conclusions

This study demonstrates the effectiveness of MD simulations in modeling the thermodynamic parameters of RNA/DNA hybrid duplexes when combined with MMPBSA and MMGBSA methods. The duplexes studied included both natural and modified forms, specifically those with PS backbone and 2′-O-MOE modifications, resulting in a comprehensive dataset of 1,300 systems. MD simulations were performed for 100 ns for all systems, followed by MMPBSA and MMGBSA analyses on the resulting MD trajectories. Correction models were then developed through empirical fitting to experimental data, significantly enhancing the predictive reliability of both MMPBSA and MMGBSA results and improving alignment with experimental values. The integration of corrected thermodynamic parameters into NN models allowed for robust prediction of melting temperatures across various duplex configurations. NN models derived from MMGBSA showed high accuracy and minimal bias, with predictive melting temperatures closely matching experimental data, particularly for modifications with stabilizing effects like MOE. In contrast, MMPBSA models displayed greater variability, likely due to inherent limitations in convergence and stability within the NN framework.

The study also revealed that certain structural modifications, such as the PS backbone, introduce complexities not fully captured by current MD simulation. This limitation led to discrepancies in melting temperature predictions, as the expected decrease in melting temperature for PS modification was not fully reflected. Conversely, the stabilizing effect of MOE modification was well-captured, resulting in an accurately predicted increase in melting temperature. These findings imply the need for further refinement of MD simulation protocols and force field parameters to ensure accurate predictions across various modified NAs, accommodating both well-matched and challenging modifications. In summary, this study provides a validated framework for integrating MD simulations with NN models, establishing a foundation for accurate thermodynamic predictions in both natural and chemically modified NAs, with promising applications in therapeutic oligonucleotide design and NA-based technologies.

## Supporting Information

Thermodynamic parameters, including enthalpy, entropy, Gibbs free energy, and melting temperature, for all natural/modified hybrid duplexes (PDF)

## Acknowledgments

This work was supported by Basic Science Research Program through the National Research Foundation of Korea (NRF) funded by the Ministry of Education (2021R1A6A1A03039696). This work was also supported by the NRF grant funded by the Korea government (MSIT) (No. RS-2023-00257666). This work was supported by the National Supercomputing Center with supercomputing resources including technical support (KSC-2023-CRE-0526).

## References

(1) Stein, C. A.; Subasinghe, C.; Shinozuka, K.; Cohen, J. S. Physicochemical Properties of Phospborothioate Oligodeoxynucleotides. Nucleic Acids Res. 1988, 16 (8), 3209–3221. 10.1093/nar/16.8.3209.

(2) Matsukura, M.; Shinozuka, K.; Zon, G.; Mitsuya, H.; Reitz, M.; Cohen, J. S.; Broder, S. Phosphorothioate Analogs of Oligodeoxynucleotides: Inhibitors of Replication and Cytopathic Effects of Human Immunodeficiency Virus. Proc. Natl. Acad. Sci. 1987, 84 (21), 7706–7710.

(3) Martin, P. Ein Neuer Zugang Zu 2′-*O*-Alkylribonucleosiden Und Eigenschaften Deren Oligonucleotide. Helv. Chim. Acta 1995, 78 (2), 486–504.

(4) Manoharan, M. 2′-Carbohydrate Modifications in Antisense Oligonucleotide Therapy: Importance of Conformation, Configuration and Conjugation. Biochim. Biophys. Acta BBA -Gene Struct. Expr. 1999, 1489 (1), 117–130.

(5) Teplova, M.; Minasov, G.; Tereshko, V.; Inamati, G. B.; Cook, P. D.; Manoharan, M.; Egli, M. Crystal Structure and Improved Antisense Properties of 2’-O-(2-Methoxyethyl)-RNA. Nat. Struct. Biol. 1999, 6 (6), 535–539.

(6) Bennett, C. F.; Swayze, E. E. RNA Targeting Therapeutics: Molecular Mechanisms of Antisense Oligonucleotides as a Therapeutic Platform. Annu. Rev. Pharmacol. Toxicol. 2010, 50 (1), 259–293.

(7) Jackson, A. L.; Burchard, J.; Leake, D.; Reynolds, A.; Schelter, J.; Guo, J.; Johnson, J. M.; Lim, L.; Karpilow, J.; Nichols, K.; Marshall, W.; Khvorova, A.; Linsley, P. S. Position-Specific Chemical Modification of siRNAs Reduces “off-Target” Transcript Silencing. RNA 2006, 12 (7), 1197–1205.

(8) Yingling, Y. G.; Shapiro, B. A. Computational Design of an RNA Hexagonal Nanoring and an RNA Nanotube. Nano Lett. 2007, 7 (8), 2328–2334.

(9) Yan, H. Nucleic Acid Nanotechnology. Science 2004, 306 (5704), 2048–2049.

(10) Burnett, J. C.; Rossi, J. J. RNA-Based Therapeutics: Current Progress and Future Prospects. Chem. Biol. 2012, 19 (1), 60–71.

(11) Schallon, A.; Synatschke, C. V.; Pergushov, D. V.; Jérôme, V.; Müller, A. H. E.; Freitag, R. DNA Melting Temperature Assay for Assessing the Stability of DNA Polyplexes Intended for Nonviral Gene Delivery. Langmuir 2011, 27 (19), 12042–12051.

(12) Reed, G. H.; Kent, J. O.; Wittwer, C. T. High-Resolution DNA Melting Analysis for Simple and Efficient Molecular Diagnostics. Pharmacogenomics 2007, 8 (6), 597–608.

(13) Owczarzy, R.; Tataurov, A. V.; Wu, Y.; Manthey, J. A.; McQuisten, K. A.; Almabrazi, H. G.; Pedersen, K. F.; Lin, Y.; Garretson, J.; McEntaggart, N. O.; Sailor, C. A.; Dawson, R. B.; Peek, A. S. IDT SciTools: A Suite for Analysis and Design of Nucleic Acid Oligomers. Nucleic Acids Res. 2008, 36 (Web Server), W163–W169.

(14) SantaLucia, J. A Unified View of Polymer, Dumbbell, and Oligonucleotide DNA Nearest-Neighbor Thermodynamics. Proc. Natl. Acad. Sci. 1998, 95 (4), 1460–1465.

(15) Breslauer, K. J.; Frank, R.; Blöcker, H.; Marky, L. A. Predicting DNA Duplex Stability from the Base Sequence. Proc. Natl. Acad. Sci. U. S. A. 1986, 83 (11), 3746.

(16) Owczarzy, R.; You, Y.; Moreira, B. G.; Manthey, J. A.; Huang, L.; Behlke, M. A.; Walder, J. A. Effects of Sodium Ions on DNA Duplex Oligomers: Improved Predictions of Melting Temperatures. Biochemistry 2004, 43 (12), 3537–3554.

(17) SantaLucia, J.; Allawi, H. T.; Seneviratne, P. A. Improved Nearest-Neighbor Parameters for Predicting DNA Duplex Stability. Biochemistry 1996, 35 (11), 3555–3562.

(18) Dowerah, D.; V. N. Uppuladinne, M.; Sarma, P. J.; Biswakarma, N.; Sonavane, U. B.; Joshi, R. R.; Ray, S. K.; Namsa, N. D.; Deka, R. Ch. Design of LNA Analogues Using a Combined Density Functional Theory and Molecular Dynamics Approach for RNA Therapeutics. ACS Omega 2023, 8 (25), 22382–22405.

(19) Lomzov, A. A.; Vorobjev, Y. N.; Pyshnyi, D. V. Evaluation of the Gibbs Free Energy Changes and Melting Temperatures of DNA/DNA Duplexes Using Hybridization Enthalpy Calculated by Molecular Dynamics Simulation. J. Phys. Chem. B 2015, 119 (49), 15221–15234.

(20) Golyshev, V. M.; Pyshnyi, D. V.; Lomzov, A. A. Calculation of Energy for RNA/RNA and DNA/RNA Duplex Formation by Molecular Dynamics Simulation. Mol. Biol. 2021, 55 (6), 927–940.

(21) Shen, L.; Johnson, T. L.; Clugston, S.; Huang, H.; Butenhof, K. J.; Stanton, R. V. Molecular Dynamics Simulation and Binding Energy Calculation for Estimation of Oligonucleotide Duplex Thermostability in RNA-Based Therapeutics. J. Chem. Inf. Model. 2011, 51 (8), 1957–1965.

(22) Torrie, G. M.; Valleau, J. P. Nonphysical Sampling Distributions in Monte Carlo Free-Energy Estimation: Umbrella Sampling. J. Comput. Phys. 1977, 23 (2), 187–199.

(23) Kollman, P. A.; Massova, I.; Reyes, C.; Kuhn, B.; Huo, S.; Chong, L.; Lee, M.; Lee, T.; Duan, Y.; Wang, W.; Donini, O.; Cieplak, P.; Srinivasan, J.; Case, D. A.; Cheatham, T. E. Calculating Structures and Free Energies of Complex Molecules: Combining Molecular Mechanics and Continuum Models. Acc. Chem. Res. 2000, 33 (12), 889–897.

(24) Srinivasan, J.; Cheatham, T. E.; Cieplak, P.; Kollman, P. A.; Case, D. A. Continuum Solvent Studies of the Stability of DNA, RNA, and Phosphoramidate−DNA Helices. J. Am. Chem. Soc. 1998, 120 (37), 9401–9409.

(25) Vester, B.; Wengel, J. LNA (Locked Nucleic Acid): High-Affinity Targeting of Complementary RNA and DNA. Biochemistry 2004, 43 (42), 13233–13241.

(26) Egholm, M.; Buchardt, O.; Nielsen, P. E.; Berg, R. H. Peptide Nucleic Acids (PNA). Oligonucleotide Analogs with an Achiral Peptide Backbone. J. Am. Chem. Soc. 1992, 114 (5), 1895–1897.

(27) Golyshev, V. M.; Abramova, T. V.; Pyshnyi, D. V.; Lomzov, A. A. Structure and Hybridization Properties of Glycine Morpholine Oligomers in Complexes with DNA and RNA: Experimental and Molecular Dynamics Studies. J. Phys. Chem. B 2019, 123 (50), 10571–10581.

(28) Ivanova, A.; Rösch, N. The Structure of LNA:DNA Hybrids from Molecular Dynamics Simulations: The Effect of Locked Nucleotides. J. Phys. Chem. A 2007, 111 (38), 9307–9319.

(29) Shields, G. C.; Laughton, C. A.; Orozco, M. Molecular Dynamics Simulation of a PNA·DNA·PNA Triple Helix in Aqueous Solution. J. Am. Chem. Soc. 1998, 120 (24), 5895–5904.

(30) Sen, S.; Nilsson, L. Molecular Dynamics of Duplex Systems Involving PNA: Structural and Dynamical Consequences of the Nucleic Acid Backbone. J. Am. Chem. Soc. 1998, 120 (4), 619–631.

(31) Pande, V.; Nilsson, L. Insights into Structure, Dynamics and Hydration of Locked Nucleic Acid (LNA) Strand-Based Duplexes from Molecular Dynamics Simulations. Nucleic Acids Res. 2008, 36 (5), 1508–1516.

(32) Sugimoto, N.; Nakano, S.; Katoh, M.; Matsumura, A.; Nakamuta, H.; Ohmichi, T.; Yoneyama, M.; Sasaki, M. Thermodynamic Parameters To Predict Stability of RNA/DNA Hybrid Duplexes. Biochemistry 1995, 34 (35), 11211–11216.

(33) Michaud-Agrawal, N.; Denning, E. J.; Woolf, T. B.; Beckstein, O. MDAnalysis: A Toolkit for the Analysis of Molecular Dynamics Simulations. J. Comput. Chem. 2011, 32 (10), 2319– 2327.

(34) Gowers, R.; Linke, M.; Barnoud, J.; Reddy, T.; Melo, M.; Seyler, S.; Domański, J.; Dotson, D.; Buchoux, S.; Kenney, I.; Beckstein, O. MDAnalysis: A Python Package for the Rapid Analysis of Molecular Dynamics Simulations; Los Alamos National Laboratory (LANL), Los Alamos, NM (United States), 2019; pp 98–105.

(35) Macke, T. J.; Case, D. A. Modeling Unusual Nucleic Acid Structures. In Molecular Modeling of Nucleic Acids; Leontis, N. B., SantaLucia, J., Jr., Eds.; American Chemical Society: Washington, DC, 1998; pp 379–393.

(36) Pettersen, E. F.; Goddard, T. D.; Huang, C. C.; Couch, G. S.; Greenblatt, D. M.; Meng, E. C.; Ferrin, T. E. UCSF Chimera—A Visualization System for Exploratory Research and Analysis. J. Comput. Chem. 2004, 25 (13), 1605–1612.

(37) Jorgensen, W. L.; Chandrasekhar, J.; Madura, J. D.; Impey, R. W.; Klein, M. L. Comparison of Simple Potential Functions for Simulating Liquid Water. J. Chem. Phys. 1983, 79 (2), 926–935.

(38) Case, D. A.; Aktulga, H. M.; Belfon, K.; Cerutti, D. S.; Cisneros, G. A.; Cruzeiro, V. W. D.; Forouzesh, N.; Giese, T. J.; Götz, A. W.; Gohlke, H.; Izadi, S.; Kasavajhala, K.; Kaymak, M. C.; King, E.; Kurtzman, T.; Lee, T.-S.; Li, P.; Liu, J.; Luchko, T.; Luo, R.; Manathunga, M.; Machado, M. R.; Nguyen, H. M.; O’Hearn, K. A.; Onufriev, A. V.; Pan, F.; Pantano, S.; Qi, R.; Rahnamoun, A.; Risheh, A.; Schott-Verdugo, S.; Shajan, A.; Swails, J.; Wang, J.; Wei, H.; Wu, X.; Wu, Y.; Zhang, S.; Zhao, S.; Zhu, Q.; Cheatham, T. E. I.; Roe, D. R.; Roitberg, A.; Simmerling, C.; York, D. M.; Nagan, M. C.; Merz, K. M. Jr. AmberTools. J. Chem. Inf. Model. 2023, 63 (20), 6183–6191.

(39) Zgarbová, M.; Šponer, J.; Otyepka, M.; Cheatham, T. E. I.; Galindo-Murillo, R.; Jurečka, P. Refinement of the Sugar–Phosphate Backbone Torsion Beta for AMBER Force Fields Improves the Description of Z-and B-DNA. J. Chem. Theory Comput. 2015, 11 (12), 5723–5736.

(40) Zgarbová, M.; Otyepka, M.; Šponer, J.; Mládek, A.; Banáš, P.; Cheatham, T. E. I.; Jurečka, P. Refinement of the Cornell et al. Nucleic Acids Force Field Based on Reference Quantum Chemical Calculations of Glycosidic Torsion Profiles. J. Chem. Theory Comput. 2011, 7 (9), 2886–2902.

(41) Frisch, M. J.; Trucks, G. W.; Schlegel, H. B.; Scuseria, G. E.; Robb, M. A.; Cheeseman, J. R.; Scalmani, G.; Barone, V.; Petersson, G. A.; Nakatsuji, H.; Li, X.; Caricato, M.; Marenich, A. V.; Bloino, J.; Janesko, B. G.; Gomperts, R.; Mennucci, B.; Hratchian, H. P.; Ortiz, J. V.; Izmaylov, A. F.; Sonnenberg, J. L.; Williams; Ding, F.; Lipparini, F.; Egidi, F.; Goings, J.; Peng, B.; Petrone, A.; Henderson, T.; Ranasinghe, D.; Zakrzewski, V. G.; Gao, J.; Rega, N.; Zheng, G.; Liang, W.; Hada, M.; Ehara, M.; Toyota, K.; Fukuda, R.; Hasegawa, J.; Ishida, M.; Nakajima, T.; Honda, Y.; Kitao, O.; Nakai, H.; Vreven, T.; Throssell, K.; Montgomery Jr., J. A.; Peralta, J. E.; Ogliaro, F.; Bearpark, M. J.; Heyd, J. J.; Brothers, E. N.; Kudin, K. N.; Staroverov, V. N.; Keith, T. A.; Kobayashi, R.; Normand, J.; Raghavachari, K.; Rendell, A. P.; Burant, J. C.; Iyengar, S. S.; Tomasi, J.; Cossi, M.; Millam, J. M.; Klene, M.; Adamo, C.; Cammi, R.; Ochterski, J. W.; Martin, R. L.; Morokuma, K.; Farkas, O.; Foresman, J. B.; Fox, D. J. Gaussian 16 Rev. C.01, 2016.

(42) Bayly, C. I.; Cieplak, P.; Cornell, W.; Kollman, P. A. A Well-Behaved Electrostatic Potential Based Method Using Charge Restraints for Deriving Atomic Charges: The RESP Model. J. Phys. Chem. 1993, 97 (40), 10269–10280.

(43) Wang, J.; Wang, W.; Kollman, P. A.; Case, D. A. Automatic Atom Type and Bond Type Perception in Molecular Mechanical Calculations. J. Mol. Graph. Model. 2006, 25 (2), 247–260.

(44) Shirts, M. R.; Klein, C.; Swails, J. M.; Yin, J.; Gilson, M. K.; Mobley, D. L.; Case, D. A.; Zhong, E. D. Lessons Learned from Comparing Molecular Dynamics Engines on the SAMPL5 Dataset. J. Comput. Aided Mol. Des. 2017, 31 (1), 147–161.

(45) Berendsen, H. J. C.; van der Spoel, D.; van Drunen, R. GROMACS: A Message-Passing Parallel Molecular Dynamics Implementation. Comput. Phys. Commun. 1995, 91 (1), 43–56.

(46) Ryckaert, J.-P.; Ciccotti, G.; Berendsen, H. J. C. Numerical Integration of the Cartesian Equations of Motion of a System with Constraints: Molecular Dynamics of *n*-Alkanes. J. Comput. Phys. 1977, 23 (3), 327–341.

(47) Darden, T.; York, D.; Pedersen, L. Particle Mesh Ewald: An N⋅log(N) Method for Ewald Sums in Large Systems. J. Chem. Phys. 1993, 98 (12), 10089–10092.

(48) Bussi, G.; Donadio, D.; Parrinello, M. Canonical Sampling through Velocity Rescaling. J. Chem. Phys. 2007, 126 (1), 014101.

(49) Parrinello, M.; Rahman, A. Polymorphic Transitions in Single Crystals: A New Molecular Dynamics Method. J. Appl. Phys. 1981, 52 (12), 7182–7190.

(50) Hoover, W. G. Canonical Dynamics: Equilibrium Phase-Space Distributions. Phys. Rev. A 1985, 31 (3), 1695–1697.

(51) Nosé, S. A Unified Formulation of the Constant Temperature Molecular Dynamics Methods. J. Chem. Phys. 1984, 81 (1), 511–519.

(52) Roe, D. R.; Cheatham, T. E. I. PTRAJ and CPPTRAJ: Software for Processing and Analysis of Molecular Dynamics Trajectory Data. J. Chem. Theory Comput. 2013, 9 (7), 3084– 3095.

(53) Miller, B. R.; McGee, T. D.; Swails, J. M.; Homeyer, N.; Gohlke, H.; Roitberg, A. E. *MMPBSA.Py* : An Efficient Program for End-State Free Energy Calculations. J. Chem. Theory Comput. 2012, 8 (9), 3314–3321.

(54) Onufriev, A.; Bashford, D.; Case, D. A. Exploring Protein Native States and Large-Scale Conformational Changes with a Modified Generalized Born Model. Proteins Struct. Funct. Bioinforma. 2004, 55 (2), 383–394.

(55) Virtanen, P.; Gommers, R.; Oliphant, T. E.; Haberland, M.; Reddy, T.; Cournapeau, D.; Burovski, E.; Peterson, P.; Weckesser, W.; Bright, J.; van der Walt, S. J.; Brett, M.; Wilson, J.; Millman, K. J.; Mayorov, N.; Nelson, A. R. J.; Jones, E.; Kern, R.; Larson, E.; Carey, C. J.; Polat, İ.; Feng, Y.; Moore, E. W.; VanderPlas, J.; Laxalde, D.; Perktold, J.; Cimrman, R.; Henriksen, I.; Quintero, E. A.; Harris, C. R.; Archibald, A. M.; Ribeiro, A. H.; Pedregosa, F.; van Mulbregt, P. SciPy 1.0: Fundamental Algorithms for Scientific Computing in Python. Nat. Methods 2020, 17 (3), 261–272.

(56) Weber, G. Optimization Method for Obtaining Nearest-Neighbour DNA Entropies and Enthalpies Directly from Melting Temperatures. Bioinformatics 2015, 31 (6), 871–877.

(57) Altmann, K.-H.; Dean, N. M.; Fabbro, D.; Freier, S. M.; Geiger, T.; Häner, R.; Hüsken, D.; Martin, P.; Monia, B. P.; Müller, M.; Natt, F.; Nicklin, P.; Phillips, J.; Pieles, U.; Sasmor, H.; Moser, H. E. Second Generation of Antisense Oligonucleotides: From Nuclease Resistance to Biological Efficacy in Animals. Chimia 1996, 50 (4), 168.

(58) Masaki, Y.; Iriyama, Y.; Nakajima, H.; Kuroda, Y.; Kanaki, T.; Furukawa, S.; Sekine, M.; Seio, K. Application of 2′-O-(2-N-Methylcarbamoylethyl) Nucleotides in RNase H-Dependent Antisense Oligonucleotides. Nucleic Acid Ther. 2018, 28 (5), 307–311.

(59) Boczkowska, M.; Guga, P.; Stec, W. J. Stereodefined Phosphorothioate Analogues of DNA: Relative Thermodynamic Stability of the Model PS-DNA/DNA and PS-DNA/RNA Complexes. Biochemistry 2002, 41 (41), 12483–12487.

(60) Lind, K. E.; Ferguson, D. M.; Mohan, V.; Manoharan, M. Structural Characteristics of 2’-O-(2-Methoxyethyl)-Modified Nucleic Acids from Molecular Dynamics Simulations. Nucleic Acids Res. 1998, 26 (16), 3694–3699.

