## Supporting Information for "Thermodynamic Parameter Estimation for Modified Oligonucleotides Using Molecular Dynamics Simulations"

**Table S1.** Hybridization Enthalpy and Entropy from Experimental Data and MD Simulations for Natural and Modified RNA/DNA Duplexes

| RNA Sequence[a] | Experimental[b] | | RNA/DNA | | | RNA/PSDNA | | | RNA/PSMOE | | |
|---|---|---|---|---|---|---|---|---|---|---|---|
| | $\Delta H°$ (kcal/mol) | $\Delta S°$ (cal/mol/K) | $\Delta H°_{PB}$ (kcal/mol) | $\Delta H°_{GB}$ (kcal/mol) | $\Delta S°$ (cal/mol/K) | $\Delta H°_{PB}$ (kcal/mol) | $\Delta H°_{GB}$ (kcal/mol) | $\Delta S°$ (cal/mol/K) | $\Delta H°_{PB}$ (kcal/mol) | $\Delta H°_{GB}$ (kcal/mol) | $\Delta S°$ (cal/mol/K) |
| CCGG | | | -42.98 | -43.72 | -81.29 | -42.78 | -44.61 | -77.52 | -50.12 | -52.51 | -90.83 |
| CGCG | | | -44.65 | -43.49 | -79.61 | -43.39 | -43.54 | -79.10 | -46.24 | -47.75 | -83.53 |
| GCGC | | | -42.05 | -42.40 | -85.44 | -40.74 | -42.31 | -82.19 | -44.04 | -46.96 | -84.95 |
| AGCCG | -41.6 | -116 | -45.68 | -46.65 | -89.75 | -47.04 | -49.62 | -91.23 | -53.29 | -56.63 | -100.09 |
| CACAG | | | -44.73 | -44.67 | -84.95 | -43.46 | -44.56 | -91.06 | -51.36 | -52.57 | -91.56 |
| CGUGC | | | -50.69 | -51.58 | -93.50 | -37.50 | -41.71 | -82.98 | -51.33 | -55.78 | -101.41 |
| GCACG | | | -43.57 | -45.58 | -95.94 | -47.77 | -50.24 | -94.38 | -52.69 | -55.82 | -94.83 |
| CGGCU | -45.8 | -132 | -49.41 | -51.16 | -91.72 | -47.69 | -52.43 | -95.01 | -48.10 | -52.08 | -94.94 |
| GGUGG | -46 | -132 | -50.07 | -52.52 | -87.26 | -48.96 | -53.55 | -90.14 | -51.89 | -57.49 | -100.19 |
| CCGCGG | | | -65.44 | -67.22 | -111.01 | -65.54 | -70.44 | -106.55 | -73.24 | -79.38 | -116.63 |
| CGAUCG | | | -50.85 | -50.41 | -93.43 | -53.81 | -56.15 | -109.56 | -60.61 | -63.57 | -112.11 |
| CGCGCG | | | -61.30 | -63.16 | -101.43 | -63.82 | -67.61 | -107.81 | -70.29 | -75.53 | -116.09 |
| CGGCCG | | | -62.05 | -65.05 | -99.43 | -63.84 | -70.26 | -109.96 | -69.02 | -75.62 | -119.91 |
| CGUACG | | | -56.36 | -57.27 | -105.69 | -53.80 | -57.70 | -101.10 | -60.14 | -65.34 | -108.77 |
| GACGUC | | | -52.20 | -54.61 | -104.41 | -46.03 | -50.53 | -98.73 | -58.78 | -64.94 | -117.25 |
| GCAUGC | | | -54.45 | -55.71 | -101.22 | -54.16 | -58.18 | -104.65 | -57.09 | -61.74 | -110.13 |
| GCCGGC | | | -65.15 | -69.11 | -115.21 | -64.51 | -71.25 | -107.73 | -68.58 | -76.62 | -122.01 |
| GCGAGC | | | -59.66 | -61.14 | -111.01 | -58.31 | -63.04 | -107.07 | -63.15 | -68.88 | -122.20 |
| GCGCGC | | | -62.71 | -64.83 | -113.68 | -62.79 | -68.75 | -111.62 | -66.77 | -73.90 | -115.94 |
| GCUAGC | | | -54.79 | -54.96 | -107.08 | -54.65 | -58.04 | -102.70 | -58.49 | -62.20 | -116.64 |
| GGAUCC | | | -51.93 | -54.24 | -94.90 | -50.72 | -55.82 | -105.06 | -57.48 | -63.38 | -110.55 |
| GGCGCC | | | -62.72 | -66.77 | -107.16 | -56.92 | -63.99 | -106.89 | -68.13 | -76.38 | -117.04 |
| GGGACC | | | -59.06 | -63.86 | -103.90 | -54.25 | -61.79 | -107.46 | -64.03 | -72.07 | -113.10 |
| GUGAAC | | | -50.00 | -51.80 | -99.37 | -48.96 | -53.25 | -100.61 | -53.10 | -58.12 | -106.14 |
| UCAUGA | | | -43.63 | -44.50 | -90.90 | -43.44 | -46.79 | -96.29 | -50.91 | -53.52 | -101.53 |
| UGAUCA | | | -41.42 | -42.49 | -84.73 | -34.07 | -37.76 | -91.25 | -44.33 | -47.23 | -96.58 |
| CCCGGG | | | -69.93 | -72.83 | -106.91 | -69.98 | -75.48 | -106.34 | -75.60 | -82.70 | -122.91 |
| ACCGCA | -45.4 | -126 | -55.48 | -58.01 | -100.06 | -53.89 | -57.74 | -101.08 | -56.94 | -61.80 | -101.48 |
| AGUUGC | | | -47.87 | -49.25 | -102.75 | -49.55 | -53.59 | -97.99 | -51.96 | -57.30 | -110.71 |
| AUGCGC | | | -54.05 | -54.96 | -98.64 | -52.22 | -56.25 | -99.09 | -56.21 | -61.68 | -109.05 |
| CCAACG | | | -53.11 | -54.36 | -100.35 | -53.70 | -56.72 | -108.69 | -57.26 | -59.54 | -111.02 |
| CGGACG | | | -60.31 | -62.38 | -104.60 | -59.96 | -64.93 | -100.45 | -66.56 | -72.18 | -113.28 |
| CGGUGC | -48.9 | -135 | -60.98 | -64.00 | -103.94 | -59.85 | -65.83 | -108.04 | -64.56 | -72.00 | -115.41 |
| CGUGCC | -49.3 | -139 | -59.94 | -62.83 | -103.34 | -57.76 | -63.28 | -107.11 | -61.63 | -69.28 | -113.62 |
| CGUUGA | | | -52.69 | -53.02 | -97.55 | -51.75 | -55.11 | -96.23 | -45.78 | -49.91 | -97.78 |
| CGUUGC | | | -57.07 | -57.82 | -105.06 | -56.40 | -60.14 | -102.42 | -58.36 | -63.45 | -110.28 |
| CGUUGU | | | -50.86 | -50.93 | -99.99 | -44.68 | -48.32 | -95.48 | -52.34 | -56.59 | -102.67 |
| GCCUGC | | | -60.24 | -62.20 | -99.79 | -59.42 | -63.94 | -103.14 | -62.50 | -68.29 | -114.19 |
| UGCGCA | | | -53.21 | -55.66 | -102.40 | -51.89 | -56.67 | -94.66 | -56.57 | -62.44 | -101.56 |
| UGUUGC | | | -49.04 | -50.57 | -99.51 | -49.52 | -53.46 | -98.64 | -44.88 | -49.76 | -97.04 |
| CAAUCG | -46.7 | -140 | -51.60 | -49.72 | -99.84 | -40.81 | -44.46 | -103.96 | -57.39 | -58.95 | -106.75 |
| CACGGC | -48.9 | -135 | -62.08 | -65.10 | -106.76 | -61.79 | -67.40 | -104.42 | -65.73 | -72.41 | -115.07 |
| CGAUUG | -49.3 | -151 | -49.57 | -48.50 | -96.20 | -51.33 | -53.01 | -101.52 | -57.20 | -59.25 | -112.59 |
| GCACCG | -49.7 | -141 | -58.58 | -61.20 | -104.70 | -57.05 | -61.70 | -110.46 | -65.30 | -70.52 | -111.95 |
| GGCACG | -51.8 | -144 | -59.64 | -62.38 | -105.56 | -57.61 | -63.17 | -108.76 | -64.68 | -71.11 | -112.53 |
| CCCAGGG | | | -76.69 | -80.80 | -113.61 | -75.58 | -83.96 | -118.00 | -82.09 | -89.80 | -136.89 |
| CAAAAAG | | | -52.51 | -51.23 | -107.85 | -54.31 | -55.28 | -107.95 | -61.81 | -61.76 | -117.10 |
| AAUACCG | -54.1 | -160 | -55.07 | -57.02 | -114.55 | -48.34 | -54.36 | -106.89 | -62.76 | -67.84 | -120.07 |
| AGCCGUG | | | -65.38 | -68.72 | -119.83 | -57.09 | -65.08 | -114.92 | -62.18 | -71.80 | -126.21 |
| AGCUUCA | -55.6 | -164 | -52.68 | -54.23 | -104.91 | -48.98 | -54.90 | -108.94 | -56.31 | -62.11 | -122.54 |
| GGACUUA | -43.8 | -125 | -54.85 | -57.26 | -110.81 | -51.18 | -57.15 | -99.58 | -58.60 | -64.76 | -121.48 |
| ACGUAUG | -58.3 | -172 | -53.41 | -55.45 | -113.67 | -51.37 | -57.09 | -116.52 | -53.12 | -59.23 | -120.04 |
| CACGGCU | -53.9 | -147 | -62.89 | -68.44 | -112.90 | -66.01 | -74.21 | -117.85 | -69.15 | -77.75 | -129.10 |

| | | | | | | | | | | | |
|---|---|---|---|---|---|---|---|---|---|---|---|
| CAUACGU | -53.2 | -158 | -57.71 | -59.34 | -109.29 | -56.19 | -61.05 | -112.70 | -62.38 | -68.15 | -115.89 |
| UAAGUCC | -46.2 | -132 | -50.54 | -53.19 | -104.17 | -51.96 | -57.33 | -108.68 | -60.01 | -65.76 | -117.32 |
| UGAAGCU | -43 | -118 | -52.59 | -53.71 | -106.66 | -51.65 | -57.01 | -106.46 | -57.80 | -62.91 | -119.54 |
| AAAAAAAA | -54 | -162 | -49.85 | -48.37 | -112.10 | -42.94 | -49.10 | -108.38 | -56.72 | -56.89 | -115.23 |
| UAGAUCUA | | | -48.24 | -51.38 | -111.65 | -50.46 | -57.48 | -116.43 | -62.20 | -68.64 | -132.34 |
| UCUAUAGA | | | -55.31 | -57.06 | -112.02 | -56.13 | -64.47 | -117.88 | -62.34 | -68.73 | -132.68 |
| CAAAAAAG | | | -60.99 | -59.42 | -116.60 | -59.03 | -61.66 | -120.75 | -68.11 | -69.38 | -130.32 |
| CAAGCUUG | | | -68.57 | -69.16 | -124.27 | -68.74 | -74.00 | -125.50 | -75.44 | -80.98 | -140.50 |
| CAUCGAUG | | | -67.45 | -69.87 | -116.12 | -65.11 | -72.39 | -118.30 | -74.22 | -81.75 | -141.26 |
| CGAUAUCG | | | -65.12 | -67.24 | -117.81 | -63.09 | -70.23 | -127.26 | -71.05 | -78.75 | -138.01 |
| CGUCGACG | | | -75.64 | -80.52 | -136.74 | -66.10 | -76.28 | -131.54 | -78.42 | -89.20 | -142.19 |
| GAAGCUUC | | | -60.50 | -62.22 | -121.47 | -63.36 | -69.65 | -122.42 | -68.55 | -75.82 | -142.12 |
| GAUCGAUC | | | -60.65 | -64.35 | -122.10 | -57.08 | -65.47 | -121.67 | -67.77 | -76.98 | -134.12 |
| GAUGCAUC | | | -61.89 | -66.34 | -118.40 | -56.94 | -65.95 | -126.86 | -66.75 | -76.21 | -133.65 |
| GGAAUUCC | | | -63.42 | -67.05 | -115.03 | -57.68 | -65.54 | -125.21 | -68.08 | -77.70 | -134.81 |
| GGACGUCC | | | -71.54 | -79.28 | -125.24 | -68.73 | -80.79 | -133.28 | -79.69 | -93.32 | -142.21 |
| GGAGCUCC | | | -72.90 | -77.92 | -118.89 | -72.04 | -81.74 | -125.93 | -77.59 | -88.83 | -142.09 |
| GGUAUACC | | | -62.86 | -69.77 | -120.18 | -60.33 | -71.28 | -133.97 | -67.35 | -80.18 | -140.31 |
| GUACGUAC | | | -63.74 | -70.39 | -126.45 | -60.70 | -71.87 | -128.67 | -68.60 | -80.52 | -136.12 |
| GUAGCUAC | | | -62.83 | -66.80 | -128.62 | -60.22 | -68.48 | -130.27 | -68.39 | -78.12 | -138.70 |
| GUUGCAAC | | | -63.87 | -69.14 | -123.90 | -62.03 | -71.78 | -119.57 | -67.80 | -77.57 | -131.39 |
| AAGCGUAG | -67.2 | -192 | -63.82 | -66.86 | -129.69 | -62.83 | -70.99 | -122.23 | -74.10 | -82.65 | -138.08 |
| AAUCCAGU | -55.9 | -161 | -59.05 | -61.69 | -116.92 | -56.66 | -64.00 | -114.64 | -65.11 | -72.40 | -130.65 |
| ACAUAUGU | | | -55.58 | -57.77 | -114.89 | -54.57 | -61.42 | -118.64 | -59.55 | -66.33 | -126.48 |
| ACCUAGUC | -56 | -159 | -61.94 | -66.13 | -120.27 | -58.19 | -66.46 | -136.61 | -69.38 | -78.60 | -135.69 |
| ACGACCUC | -55 | -150 | -66.23 | -72.22 | -132.04 | -66.08 | -76.05 | -129.03 | -74.39 | -85.22 | -143.96 |
| AGAGAGAG | | | -66.29 | -68.06 | -117.58 | -68.07 | -75.13 | -114.70 | -72.87 | -81.03 | -144.31 |
| AGAGCUCU | | | -61.71 | -64.69 | -124.26 | -62.51 | -69.75 | -119.96 | -67.03 | -75.57 | -141.37 |
| AGCGUAAG | -60.1 | -170 | -63.79 | -67.15 | -127.06 | -61.75 | -70.17 | -124.20 | -71.38 | -81.37 | -135.29 |
| AGUCCUGA | -54.7 | -155 | -63.71 | -68.24 | -115.06 | -60.97 | -70.30 | -122.36 | -67.30 | -78.49 | -134.74 |
| AUGCGCAU | | | -63.20 | -67.67 | -122.82 | -61.83 | -71.00 | -121.88 | -66.79 | -76.43 | -129.81 |
| CACGGCUC | -71.6 | -200 | -74.51 | -80.66 | -131.05 | -73.32 | -83.89 | -134.67 | -80.92 | -92.56 | -150.49 |
| CCAUAUGG | | | -68.99 | -72.64 | -114.30 | -68.10 | -75.93 | -120.84 | -75.34 | -83.24 | -140.47 |
| CGCGUAUA | | | -66.29 | -70.25 | -123.89 | -61.19 | -69.73 | -127.67 | -61.98 | -72.36 | -132.87 |
| CGCUGUAA | -60.4 | -172 | -62.94 | -66.78 | -121.46 | -65.64 | -73.68 | -125.91 | -71.75 | -80.84 | -139.05 |
| CGUCGUCC | | | -69.36 | -75.74 | -128.50 | -65.88 | -76.17 | -127.76 | -77.76 | -90.12 | -136.74 |
| CUAGUGGA | -63.4 | -178 | -68.31 | -72.43 | -121.41 | -65.36 | -73.57 | -120.94 | -74.72 | -84.45 | -135.76 |
| CUCACGGC | -70.3 | -196 | -75.60 | -81.79 | -130.76 | -71.43 | -81.03 | -131.90 | -81.31 | -92.60 | -144.29 |
| CUGAGUCC | -60.8 | -171 | -68.95 | -73.48 | -124.86 | -67.12 | -75.92 | -127.06 | -75.06 | -85.15 | -141.12 |
| GAAUAUUC | | | -53.08 | -54.50 | -111.52 | -47.64 | -53.53 | -113.83 | -58.57 | -64.60 | -126.94 |
| GACUAGUC | | | -61.83 | -66.54 | -119.07 | -61.24 | -69.85 | -123.74 | -68.28 | -77.88 | -141.36 |
| GAGUACUC | | | -59.78 | -65.06 | -121.12 | -60.61 | -69.93 | -125.34 | -66.84 | -77.08 | -137.06 |
| GAUUAAUC | | | -52.73 | -54.45 | -108.74 | -50.28 | -56.73 | -119.83 | -56.76 | -63.21 | -129.59 |
| GCAUAUGC | | | -64.73 | -68.47 | -126.38 | -62.45 | -71.23 | -122.78 | -67.90 | -76.95 | -135.89 |
| GCCAGUUA | -62.9 | -180 | -64.90 | -69.65 | -121.61 | -58.98 | -68.07 | -120.10 | -70.08 | -79.37 | -135.34 |
| GGACCUCG | | | -75.44 | -80.59 | -126.27 | -71.87 | -81.69 | -123.50 | -79.56 | -90.63 | -145.07 |
| GGAGCACG | | | -75.45 | -80.52 | -126.98 | -73.59 | -83.83 | -127.71 | -80.62 | -91.99 | -138.49 |
| GGUGCCAA | | | -70.46 | -78.33 | -122.92 | -67.18 | -79.63 | -126.38 | -73.95 | -86.23 | -136.08 |
| GUCGAACA | | | -64.13 | -69.84 | -122.17 | -60.59 | -70.59 | -133.65 | -68.07 | -77.46 | -129.88 |
| GUCUAGAC | | | -62.80 | -67.37 | -120.21 | -61.81 | -71.32 | -121.54 | -68.60 | -78.50 | -136.84 |
| UAGGCCUA | | | -60.96 | -69.05 | -132.23 | -62.89 | -73.14 | -128.52 | -73.78 | -83.63 | -146.20 |
| UAUGCAUA | | | -55.07 | -58.04 | -119.47 | -52.90 | -60.19 | -123.88 | -59.88 | -67.62 | -131.35 |
| UGAGCUCA | | | -60.95 | -64.37 | -114.83 | -60.32 | -68.19 | -114.34 | -64.17 | -72.46 | -133.37 |
| AAUGUCGC | -64.9 | -186 | -63.56 | -67.21 | -127.95 | -61.20 | -69.91 | -125.96 | -69.21 | -78.87 | -136.07 |
| ACUGGAUU | -57.8 | -163 | -62.93 | -66.84 | -113.78 | -61.35 | -70.87 | -125.35 | -59.17 | -67.70 | -129.84 |
| CAACAGCA | -52.7 | -145 | -68.80 | -72.46 | -122.84 | -66.29 | -74.18 | -125.30 | -67.44 | -74.35 | -129.82 |
| CUACGCUU | -54.3 | -153 | -64.25 | -67.57 | -126.76 | -61.73 | -69.46 | -126.66 | -68.98 | -77.16 | -135.29 |
| CUUACGCU | -52.3 | -148 | -64.37 | -67.66 | -124.72 | -63.17 | -70.60 | -125.13 | -70.02 | -78.28 | -142.37 |

| | | | | | | | | | | | |
|---|---|---|---|---|---|---|---|---|---|---|---|
| GACUAGGU | -57 | -158 | -65.16 | -70.19 | -121.03 | -65.47 | -75.17 | -125.00 | -69.20 | -79.02 | -142.24 |
| GAGCCGUG | -67.3 | -187 | -76.49 | -81.65 | -129.49 | -70.97 | -81.47 | -126.71 | -82.68 | -93.98 | -148.53 |
| GAGGUCGU | -71.9 | -202 | -71.18 | -76.09 | -126.05 | -71.83 | -81.44 | -121.99 | -73.32 | -84.96 | -129.19 |
| GCCGUGAG | -71.4 | -199 | -75.38 | -81.52 | -134.66 | -74.13 | -85.31 | -132.92 | -80.90 | -92.70 | -147.20 |
| GCGACAUU | -60.6 | -170 | -62.84 | -68.44 | -130.19 | -64.85 | -74.70 | -130.49 | -64.79 | -74.30 | -132.16 |
| UAACUGGC | -62.6 | -174 | -64.55 | -69.04 | -124.47 | -65.40 | -74.19 | -121.11 | -71.88 | -81.25 | -132.74 |
| UCCACUAG | -64.1 | -184 | -65.31 | -69.30 | -113.59 | -60.28 | -68.40 | -116.81 | -72.98 | -80.70 | -130.70 |
| UGCUGUUG | -52 | -146 | -65.06 | -69.63 | -113.82 | -63.59 | -73.24 | -116.76 | -69.52 | -78.88 | -136.28 |
| UGUUCGAC | -66.2 | -190 | -62.47 | -67.87 | -122.98 | -59.40 | -69.18 | -118.83 | -67.79 | -78.84 | -135.67 |
| UUACAGCG | -58.4 | -166 | -66.04 | -69.85 | -125.27 | -61.73 | -69.81 | -126.99 | -73.37 | -81.48 | -137.88 |
| UUGGCACC | -56.2 | -153 | -68.90 | -76.67 | -118.79 | -64.89 | -76.33 | -128.25 | -74.87 | -87.25 | -137.73 |
| CAAAAAAAG | | | -65.15 | -64.88 | -129.83 | -64.76 | -68.44 | -125.65 | -73.28 | -76.52 | -141.78 |
| CAAACAAAG | | | -67.61 | -70.91 | -135.36 | -65.24 | -72.70 | -138.38 | -77.55 | -83.50 | -144.57 |
| CAAAUAAAG | | | -62.85 | -64.17 | -126.29 | -60.95 | -66.75 | -136.05 | -72.24 | -77.14 | -144.97 |
| CAAAGAAAG | | | -70.87 | -71.76 | -132.92 | -70.56 | -76.00 | -130.62 | -77.68 | -83.38 | -143.75 |
| AAAAAAAAA | | | -56.01 | -55.18 | -122.09 | -53.18 | -56.59 | -122.27 | -62.18 | -64.10 | -129.55 |
| AUAACUGGC | -60.7 | -168 | -69.70 | -75.84 | -133.07 | -69.44 | -80.17 | -133.31 | -74.74 | -86.81 | -140.95 |
| AUCUAUCCG | -59.9 | -171 | -66.81 | -71.12 | -129.60 | -63.85 | -73.12 | -131.53 | -72.04 | -82.90 | -147.07 |
| CGCUGUUAC | -71.5 | -205 | -75.17 | -81.57 | -144.75 | -74.18 | -85.19 | -138.42 | -77.42 | -90.77 | -154.95 |
| GCCAGUUAA | -63.1 | -179 | -68.35 | -75.04 | -132.38 | -66.12 | -76.18 | -128.41 | -77.28 | -88.81 | -144.67 |
| CAACAGCAA | -63.3 | -177 | -73.28 | -79.22 | -133.67 | -67.27 | -79.36 | -142.07 | -75.73 | -85.53 | -142.91 |
| CAACAGCAU | -71 | -200 | -69.19 | -74.62 | -135.82 | -68.64 | -80.66 | -139.29 | -75.90 | -86.20 | -142.28 |
| CGCUGUUAG | -67.7 | -193 | -74.16 | -79.06 | -139.10 | -76.60 | -86.17 | -135.97 | -81.46 | -92.95 | -157.85 |
| CUAACAGCG | -70.8 | -199 | -76.97 | -81.25 | -140.78 | -75.09 | -84.40 | -141.12 | -79.41 | -89.71 | -150.17 |
| GUAACAGCG | -77.5 | -221 | -75.20 | -81.45 | -143.76 | -73.50 | -84.75 | -140.56 | -81.54 | -93.45 | -155.50 |
| UUAACUGGC | -67.3 | -189 | -68.65 | -74.83 | -127.39 | -68.89 | -80.05 | -133.04 | -76.32 | -88.37 | -145.58 |
| AUGAGCUCAU | | | -71.90 | -78.92 | -140.03 | -66.68 | -79.71 | -139.71 | -76.88 | -91.22 | -168.49 |
| GCGAAUUCGC | | | -81.85 | -89.77 | -154.15 | -80.13 | -94.30 | -154.11 | -86.81 | -102.63 | -173.43 |
| GCGAAAAGCG | | | -84.85 | -90.96 | -160.87 | -83.99 | -95.99 | -152.19 | -91.96 | -105.41 | -167.28 |
| AUCAAUCAUA | | | -62.05 | -68.43 | -136.12 | -60.66 | -70.92 | -148.26 | -65.67 | -76.01 | -150.25 |
| UUGUAGUCAU | | | -67.24 | -75.93 | -139.32 | -60.73 | -74.68 | -137.99 | -72.98 | -88.29 | -159.88 |
| GAAAUGAAAG | | | -70.48 | -75.22 | -141.63 | -70.26 | -81.74 | -143.49 | -78.38 | -88.46 | -159.94 |
| CCAACUUCUU | | | -74.24 | -80.38 | -145.27 | -70.14 | -81.21 | -145.15 | -79.64 | -90.63 | -166.80 |
| AUCGUCUGGA | | | -77.48 | -87.17 | -138.80 | -75.19 | -91.84 | -142.70 | -82.34 | -99.02 | -163.18 |
| AGCGUAAGUC | | | -75.91 | -85.14 | -147.15 | -75.57 | -90.42 | -145.59 | -82.83 | -99.72 | -164.06 |
| CGAUCUGCGA | | | -79.19 | -88.86 | -138.80 | -79.19 | -92.54 | -143.44 | -83.88 | -100.02 | -168.36 |
| UGGCGAGCAC | | | -82.82 | -94.76 | -149.20 | -85.41 | -102.75 | -146.81 | -92.30 | -112.64 | -176.41 |
| GAUGCGCUCG | | | -84.09 | -93.34 | -155.02 | -84.08 | -99.28 | -152.20 | -91.96 | -109.21 | -169.89 |
| GGGACCGCCU | | | -93.30 | -108.64 | -161.70 | -93.77 | -112.45 | -153.39 | -97.67 | -119.25 | -177.16 |
| AAAAAAAAAA | | | -57.62 | -58.53 | -139.73 | -56.78 | -62.68 | -137.10 | -67.44 | -71.29 | -144.33 |
| CGGCAAGCGC | | | -93.66 | -104.30 | -162.53 | -91.45 | -107.89 | -160.73 | -98.10 | -117.02 | -172.70 |
| UAGGUUAUAA | | | -61.08 | -68.57 | -144.40 | -59.00 | -71.24 | -143.75 | -65.79 | -81.25 | -161.23 |
| CGUACACAUGC | | | -86.40 | -100.18 | -156.07 | -85.63 | -104.17 | -159.22 | -91.06 | -112.38 | -180.47 |
| CCAUUGCUACC | | | -84.59 | -96.72 | -159.72 | -84.52 | -101.08 | -161.63 | -91.25 | -110.44 | -182.15 |
| CCAUCGCUACC | | | -88.72 | -101.92 | -159.52 | -86.37 | -104.27 | -162.04 | -97.62 | -117.25 | -180.68 |
| ACGUAUUAUGC | -95.4 | -279 | -76.34 | -86.53 | -157.88 | -71.14 | -87.92 | -156.09 | -79.85 | -98.72 | -177.61 |
| GCAUAAUACGU | -97.3 | -282 | -76.67 | -86.46 | -160.77 | -76.23 | -91.16 | -161.92 | -81.90 | -98.66 | -170.06 |
| CGCGAAUUCGCG | | | -100.66 | -113.23 | -178.63 | -99.69 | -118.40 | -170.47 | -104.53 | -127.58 | -197.64 |
| CUGACAAGUGUC | | | -91.08 | -105.22 | -168.30 | -90.79 | -110.60 | -177.66 | -96.86 | -120.24 | -197.43 |
| UUUUAUAAUAAA | | | -59.75 | -67.71 | -151.83 | -58.94 | -71.32 | -162.43 | -71.03 | -85.14 | -175.00 |
| AUUGGAUACAAA | -93.6 | -267 | -76.49 | -89.72 | -163.22 | -73.97 | -93.15 | -164.13 | -81.41 | -103.23 | -187.78 |
| CAACCAACCAAC | | | -93.73 | -108.93 | -176.48 | -91.44 | -110.47 | -170.80 | -100.71 | -121.65 | -190.75 |
| CUUCCUUCCUUC | | | -89.04 | -99.22 | -161.61 | -90.28 | -103.57 | -174.09 | -93.64 | -113.19 | -200.33 |
| GGAACAAGAUGC | | | -92.37 | -106.05 | -167.13 | -91.77 | -111.80 | -159.26 | -97.99 | -120.77 | -197.16 |
| GGAACUUGAUGC | | | -91.32 | -104.40 | -168.14 | -91.47 | -110.77 | -173.81 | -94.73 | -118.19 | -192.38 |
| AAUGGAUUACAA | -90.7 | -260 | -76.03 | -89.41 | -163.41 | -75.72 | -94.02 | -164.14 | -84.20 | -105.25 | -182.49 |
| GUCAGGAAUCUG | -79.8 | -220 | -89.42 | -103.90 | -168.73 | -88.51 | -107.49 | -166.82 | -98.18 | -120.66 | -199.71 |
| UUGUAAUCCAUU | -76.9 | -221 | -75.17 | -88.75 | -158.68 | -77.76 | -94.81 | -167.12 | -75.57 | -97.07 | -179.83 |

| | | | | | | | | | |
|---|---|---|---|---|---|---|---|---|---|
| UUUGUAUCCAAU | -76.52 | -88.54 | -153.41 | -74.58 | -91.69 | -160.84 | -77.96 | -98.81 | -185.76 |
| CGCAUGGGUACGC | -110.16 | -130.53 | -195.90 | -109.50 | -137.01 | -189.18 | -113.06 | -146.03 | -216.45 |
| AGCCUAACUCAGC | -91.64 | -109.25 | -177.05 | -97.83 | -119.69 | -187.67 | -101.96 | -128.61 | -209.71 |
| CAUAUGGCCAUAUG | -100.01 | -119.34 | -186.50 | -88.48 | -115.19 | -190.76 | -106.71 | -136.43 | -216.25 |
| GUAUACCGGUAUAC | -97.38 | -119.53 | -193.11 | -95.21 | -123.54 | -201.15 | -102.45 | -136.55 | -230.12 |
| ACAUUAUUAUUACA | -63.42 | -77.00 | -190.48 | -76.87 | -97.87 | -188.25 | -78.28 | -102.48 | -203.38 |
| UACUAACAUUAACUA | -81.16 | -100.34 | -190.92 | -76.25 | -100.59 | -192.52 | -92.25 | -122.15 | -215.61 |
| AUACUUACUGAUUAG | -90.00 | -107.65 | -189.97 | -64.39 | -90.32 | -208.15 | -95.87 | -125.27 | -229.98 |
| GUACACUGUCUUAUA | -87.78 | -110.56 | -190.57 | -90.80 | -119.42 | -204.76 | -95.36 | -129.32 | -225.09 |
| GUAUGAGAGACUUUA | -96.66 | -116.21 | -197.26 | -96.45 | -123.17 | -193.29 | -93.79 | -128.44 | -222.80 |
| UUCUACCUAUGUGAU | -94.95 | -115.91 | -191.56 | -94.92 | -121.37 | -198.26 | -98.46 | -130.99 | -229.98 |
| AGUAGUAAUCACACC | -93.55 | -118.32 | -203.60 | -96.41 | -126.72 | -207.28 | -101.36 | -139.22 | -229.51 |
| AUCGUCUCGGUAUAA | -94.97 | -117.82 | -199.41 | -93.68 | -123.31 | -205.52 | -103.89 | -139.35 | -230.07 |
| ACGACAGGUUUACCA | -94.23 | -119.74 | -208.03 | -99.46 | -132.78 | -215.73 | -111.01 | -150.54 | -238.89 |
| CUUUCAUGUCCGCAU | -103.40 | -125.10 | -203.69 | -81.42 | -111.42 | -200.55 | -109.86 | -142.06 | -238.16 |
| UGGAUGUGUGAACAC | -105.83 | -134.35 | -206.85 | -98.77 | -132.34 | -203.99 | -103.90 | -145.68 | -237.34 |
| ACCCCGCAAUACAUG | -111.16 | -136.13 | -202.22 | -101.06 | -133.50 | -206.20 | -109.11 | -144.00 | -224.34 |
| GCAGUGGAUGUGAGA | -111.42 | -137.17 | -209.75 | -110.78 | -144.44 | -199.73 | -106.88 | -147.59 | -237.27 |
| GGUCCUUACUUGGUG | -112.49 | -136.44 | -209.61 | -99.24 | -134.26 | -203.03 | -104.26 | -141.42 | -242.34 |
| CGCCUCAUGCUCAUC | -111.10 | -134.14 | -203.09 | -110.16 | -138.96 | -205.83 | -116.92 | -150.89 | -242.27 |
| AAAUAGCCGGGCCGC | -120.53 | -146.58 | -226.68 | -118.55 | -152.90 | -220.41 | -121.79 | -163.65 | -251.33 |
| CCAGCCAGUCUCUCC | -116.05 | -140.69 | -207.86 | -116.58 | -146.84 | -211.42 | -122.04 | -159.53 | -245.44 |
| GACGACAAGACCGCG | -119.40 | -145.86 | -215.22 | -115.00 | -147.23 | -215.51 | -117.34 | -155.67 | -241.97 |
| CAGCCUCGUCGCAGC | -121.48 | -147.83 | -224.89 | -122.24 | -155.45 | -226.00 | -134.00 | -171.23 | -250.60 |
| CUCGCGGUCGAAGCG | -119.77 | -144.58 | -217.73 | -117.81 | -149.76 | -218.93 | -107.93 | -146.77 | -241.01 |
| GCGUCGGUCCGGGCU | -128.15 | -159.37 | -226.58 | -129.71 | -167.88 | -226.64 | -129.00 | -174.76 | -253.29 |
| CAACUUGAUAUUAUUA | -86.91 | -106.92 | -204.96 | -87.92 | -114.49 | -207.17 | -94.34 | -127.34 | -238.60 |
| CAUAUUGGCCAAUAUG | -108.22 | -131.32 | -209.80 | -99.31 | -131.28 | -219.96 | -112.52 | -149.50 | -252.92 |
| GUAUAACCGGUUAUAC | -100.06 | -126.71 | -214.24 | -94.63 | -126.19 | -214.15 | -109.37 | -150.42 | -252.89 |
| CGCGUACGCGUACGCG | -125.03 | -159.71 | -247.12 | -125.30 | -163.90 | -232.51 | -137.84 | -185.04 | -263.31 |
| CAACUUGAUAUUAAUA | -92.80 | -111.96 | -208.63 | -87.78 | -115.38 | -196.90 | -92.66 | -125.44 | -237.79 |
| UAUGUAUAUUUUGUAAUCAG | -102.19 | -134.58 | -248.60 | -97.17 | -139.20 | -252.83 | -107.18 | -160.02 | -293.04 |
| UUCAAGUUAAACAUUCUAUC | -102.49 | -136.46 | -240.02 | -97.09 | -142.30 | -253.50 | -114.22 | -165.07 | -295.38 |
| UGAUUCUACCUAUGUGAUUU | -109.48 | -148.17 | -248.14 | -113.32 | -159.60 | -263.65 | -105.22 | -159.40 | -286.19 |
| GAGAUUGUUUCCCUUUCAAA | -95.35 | -133.68 | -256.29 | -103.15 | -145.98 | -254.84 | -105.31 | -166.20 | -308.62 |
| AUGCAAUGCUACAUAUUCGC | -87.02 | -125.38 | -247.56 | -86.75 | -137.63 | -254.04 | -124.69 | -181.46 | -306.70 |
| CCACUAUACCAUCUAUGUAC | -112.75 | -154.04 | -267.34 | -111.76 | -158.54 | -258.57 | -129.49 | -184.62 | -299.08 |
| CCAUCAUUGUGUCUACCUCA | -119.28 | -160.18 | -264.45 | -58.67 | -102.99 | -171.41 | -128.31 | -186.38 | -306.91 |
| CGGGACCAACUAAAGGAAAU | -109.45 | -161.47 | -280.52 | -106.91 | -155.70 | -246.89 | -136.69 | -195.12 | -313.46 |
| UAGUGGCGAUUAGAUUCUGC | -118.37 | -158.53 | -271.00 | -121.93 | -172.77 | -276.35 | -130.60 | -191.71 | -308.52 |
| AGCUGCAGUGGAUGUGAGAA | -129.32 | -172.16 | -265.22 | -114.30 | -167.31 | -242.91 | -127.53 | -190.23 | -318.03 |
| UACUUCCAGUGCUCAGCGUA | -103.67 | -140.38 | -250.39 | -116.65 | -161.72 | -258.74 | -119.67 | -180.57 | -312.87 |
| CAGUGAGACAGCAAUGGUCG | -70.86 | -105.91 | -237.51 | -119.16 | -169.66 | -269.23 | -131.87 | -193.96 | -315.89 |
| CGAGCUUAUCCCUAUCCCUC | -89.68 | -129.22 | -258.39 | -124.88 | -175.79 | -277.40 | -119.87 | -177.33 | -307.97 |
| CGUACUAGCGUUGGUCAUGG | -136.88 | -180.26 | -272.24 | -138.22 | -191.00 | -272.69 | -143.31 | -205.19 | -325.80 |
| AAGGCGAGUCAGGCUCAGUG | -138.03 | -182.99 | -273.35 | -127.20 | -184.00 | -275.95 | -147.93 | -212.65 | -314.67 |
| ACCGACGACGCUGAUCCGAU | -134.42 | -181.39 | -276.13 | -129.68 | -184.47 | -284.55 | -143.81 | -204.54 | -313.69 |
| AGCAGUCCGCCACACCCUGA | -142.21 | -191.50 | -277.73 | -136.83 | -192.63 | -277.66 | -154.68 | -219.95 | -315.63 |
| CAGCCUCGUUCGCACAGCCC | -131.08 | -176.74 | -271.07 | -116.01 | -166.65 | -278.44 | -156.89 | -220.55 | -310.13 |
| GUGGUGGGCCGUGCGCUCUG | -134.16 | -192.29 | -294.69 | -146.58 | -206.64 | -286.83 | -155.71 | -228.09 | -337.35 |
| GUCCACGCCCGGUGCGACGG | -143.41 | -196.43 | -283.35 | -158.15 | -219.72 | -305.57 | -166.12 | -235.66 | -334.82 |
| GAUAUAGCAAAAUUCUAAGUUAAUA | -120.66 | -170.75 | -303.61 | -69.87 | -139.96 | -303.13 | -116.10 | -193.79 | -362.85 |
| AUAACUUUACGUGUGUGACCUAUUA | -106.74 | -167.11 | -299.62 | -74.83 | -150.31 | -287.97 | -123.85 | -207.94 | -369.78 |
| GUUCUAUACUCUUGAAGUUGAUUAC | -114.47 | -168.56 | -311.74 | -110.22 | -176.06 | -315.52 | -128.11 | -210.14 | -384.44 |
| CCCUGCACUUUAACUGAAUUGUUUA | -129.05 | -191.16 | -306.55 | -133.19 | -199.53 | -317.34 | -127.69 | -208.52 | -369.58 |
| UAACCAUACUGAAUACCUUUUGACG | -70.16 | -145.75 | -295.99 | -133.29 | -200.93 | -331.90 | -102.67 | -182.89 | -351.00 |
| UCCACACGGUAGUAAAAUUAGGCUU | -127.05 | -188.29 | -322.91 | -90.19 | -172.55 | -312.40 | -146.26 | -233.33 | -380.52 |
| UUCCAAAAGGAGUUAUGAGUUGCGA | -120.43 | -174.61 | -309.40 | -79.32 | -158.22 | -292.10 | -145.16 | -230.99 | -393.27 |

| | | | | | | | | | |
|---|---|---|---|---|---|---|---|---|---|
| AAUAUCUCUCAUGCGCCAAGCUACA | -93.38 | -148.44 | -307.81 | -108.17 | -179.16 | -328.12 | -129.61 | -211.79 | -356.77 |
| UAGUAUAUCGCAGCAUCAUACAGGC | -139.08 | -201.52 | -330.33 | -108.14 | -179.65 | -325.67 | -136.56 | -220.75 | -369.78 |
| UGGAUUCUACUCAACCUUAGUCUGG | -138.71 | -197.53 | -322.41 | -138.57 | -205.68 | -322.64 | -132.90 | -216.27 | -383.36 |
| CGGAAUCCAUGUUACUUCGGCUAUC | -110.95 | -177.87 | -323.24 | -122.85 | -201.27 | -341.31 | -102.88 | -186.52 | -352.73 |
| CUGGUCUGGAUCUGAGAACUUCAGG | -107.31 | -175.95 | -321.37 | -156.63 | -228.74 | -315.49 | -158.53 | -248.68 | -399.15 |
| ACAGCGAAUGGACCUACGUGGCCUU | -144.01 | -212.23 | -332.45 | -158.53 | -238.04 | -339.57 | -163.52 | -256.99 | -384.71 |
| AGCAAGUCGAGCAGGGCCUACGUUU | -133.43 | -197.38 | -325.82 | -150.79 | -224.73 | -333.90 | -140.86 | -230.62 | -385.39 |
| GCGAGCGACAGGUUACUUGGCUGAU | -155.37 | -222.36 | -349.82 | -151.72 | -230.19 | -348.67 | -160.09 | -252.75 | -402.29 |
| AAAGGUGUCGCGGAGAGUCGUGCUG | -154.58 | -222.31 | -330.40 | -160.13 | -238.35 | -328.66 | -170.47 | -264.42 | -394.64 |
| AUGGGUGGGAGCCUCGGUAGCAGCC | -173.53 | -241.59 | -336.86 | -148.44 | -237.32 | -343.86 | -177.57 | -279.18 | -412.29 |
| CAGUGGGCUCCUGGGCGUGCUGGUC | -166.16 | -236.99 | -346.56 | -114.99 | -193.52 | -315.84 | -156.79 | -257.46 | -396.91 |
| GCCAACUCCGUCGCCGUUCGUGCGC | -121.26 | -186.72 | -354.86 | -162.81 | -245.39 | -359.68 | -187.45 | -279.28 | -407.25 |
| ACGGGUCCCCGCACCGCACCGCCAG | -171.52 | -248.57 | -351.05 | -174.66 | -261.96 | -355.80 | -202.62 | -301.64 | -419.05 |
| UUAUGUAUUAAGUUAUAUAGUAGUAGUAGU | -87.00 | -167.93 | -371.71 | -70.45 | -164.46 | -344.09 | -79.68 | -187.90 | -432.10 |
| AUUGAUAUCCUUUUCUAUUCAUCUUUCAUU | -115.82 | -188.42 | -351.70 | -113.77 | -212.91 | -369.44 | -119.16 | -227.10 | -431.58 |
| AAAGUACAUCAACAUAGAGAAUUGCAUUUC | -122.66 | -206.91 | -369.89 | -82.85 | -190.59 | -375.50 | -139.11 | -248.91 | -432.09 |
| CUUAAGAUAUGAGAACUUCAACUAAUGUGU | -114.71 | -188.90 | -363.01 | -94.02 | -188.68 | -358.28 | -105.32 | -214.69 | -421.84 |
| CUCAACUUGCGGUAAAUAAAUCGCUUAAUC | -146.98 | -222.80 | -372.44 | -92.85 | -191.44 | -371.38 | -143.40 | -253.98 | -459.74 |
| UAUUGAGAACAAGUGUCCGAUUAGCAGAAA | -113.20 | -193.93 | -376.78 | -137.59 | -235.42 | -366.43 | -121.77 | -235.50 | -447.84 |
| GUCAUACGACUGAGUGCAACAUUGUUCAAA | -76.50 | -178.66 | -379.81 | -126.01 | -228.01 | -377.87 | -71.85 | -207.82 | -435.68 |
| AACCUGCAACAUGGAGUUUUUGUCUCAUGC | -127.35 | -210.71 | -372.80 | -98.48 | -203.96 | -379.28 | -129.44 | -247.97 | -445.29 |
| CCGUGCGGUGUGUACGUUUUAUUCAUCAUA | -155.39 | -246.13 | -396.76 | -152.19 | -253.56 | -402.64 | -156.36 | -282.12 | -458.15 |
| GUUCACGUCCGAAAGCUCGAAAAAGGAUAC | -118.76 | -207.13 | -383.72 | -147.35 | -244.30 | -384.24 | -149.16 | -258.19 | -455.99 |
| AGUCUGGUCUGGAUCUGAGAACUUCAGGCU | -114.36 | -201.79 | -373.53 | -147.78 | -252.67 | -370.96 | -157.55 | -278.37 | -461.62 |
| UCGGAGAAAUCACUGAGCUGCCUGAGAAGA | -108.80 | -197.18 | -380.26 | -117.29 | -234.82 | -387.29 | -124.38 | -233.92 | -433.90 |
| CUUCAACGGAUCAGGUAGGACUGUGGUGGG | -94.01 | -201.80 | -382.74 | -140.37 | -243.51 | -361.36 | -161.37 | -292.11 | -463.44 |
| ACGCCCACAGGAUUAGGCUGGCCCACAUUG | -138.82 | -238.69 | -386.47 | -127.76 | -233.14 | -398.33 | -142.39 | -272.04 | -456.99 |
| GUUAUUCCGCAGUCCGAUGGCAGCAGGCUC | -143.69 | -231.10 | -380.48 | -83.42 | -209.82 | -400.32 | -167.75 | -288.82 | -452.37 |
| UCAGUAGGCGUGACGCAGAGCUGGCGAUGG | -143.97 | -238.16 | -393.71 | -145.18 | -255.12 | -393.04 | -174.80 | -297.25 | -467.67 |
| CGCGCCACGUGUGAUCUACAGCCGUUCGGC | -161.75 | -253.57 | -407.37 | -151.06 | -253.97 | -413.40 | -139.37 | -261.15 | -471.41 |
| GACCUGACGUGGACCGCUCCUGGGCGUGGU | -121.60 | -225.37 | -391.61 | -177.25 | -286.00 | -399.49 | -145.49 | -268.24 | -442.98 |
| GCCCCUCCACUGGCCGACGGCAGCAGGCUC | -165.09 | -260.27 | -393.74 | -203.92 | -316.87 | -429.80 | -203.24 | -328.48 | -492.30 |
| CGCCGCUGCCGACUGGAGGAGCGCGGGACG | -169.15 | -259.91 | -431.26 | -155.28 | -290.29 | -418.71 | -219.22 | -350.64 | -500.54 |

[a] All duplexes consist of the denoted RNA strand and its complementary DNA strand.

[b] Sugimoto, N.; Nakano, S. I.; Katoh, M.; Matsumura, A.; Nakamuta, H.; Ohmichi, T.; Yoneyam, M.; Sasaki, M. Thermodynamic parameters to predict stability of RNA/DNA hybrid duplexes. *Biochemistry* **1995**, *34*, 11211-11216.

**Table S1.** (Continued)

| RNA Sequence[a] | RNA/PSDNA-PSMOE | | | RNA/PSMOE-PSDNA | | |
|---|---|---|---|---|---|---|
| | $\Delta H°_{PB}$ (kcal/mol) | $\Delta H°_{GB}$ (kcal/mol) | $\Delta S°$ (cal/mol/K) | $\Delta H°_{PB}$ (kcal/mol) | $\Delta H°_{GB}$ (kcal/mol) | $\Delta S°$ (cal/mol/K) |
| CCGG | -39.11 | -41.79 | -82.95 | -35.86 | -38.59 | -84.30 |
| CGCG | -46.22 | -46.72 | -80.94 | -39.81 | -41.54 | -83.11 |
| GCGC | -41.18 | -43.76 | -82.00 | -42.70 | -44.84 | -79.36 |
| AGCCG | -47.75 | -50.89 | -86.80 | -47.92 | -51.51 | -97.92 |
| CACAG | -44.63 | -46.35 | -89.47 | -43.71 | -45.62 | -91.21 |
| CGUGC | -47.58 | -51.63 | -100.80 | -47.21 | -50.89 | -89.48 |
| GCACG | -48.89 | -51.12 | -95.09 | -47.21 | -50.83 | -95.85 |
| CGGCU | -49.74 | -53.16 | -98.20 | -51.93 | -55.99 | -95.42 |
| GGUGG | -48.66 | -54.98 | -100.04 | -54.90 | -59.93 | -94.14 |
| CCGCGG | -65.10 | -71.28 | -113.12 | -69.42 | -74.05 | -113.71 |
| CGAUCG | -58.91 | -60.68 | -104.55 | -50.01 | -54.16 | -113.28 |
| CGCGCG | -69.49 | -72.96 | -110.64 | -62.63 | -68.84 | -110.64 |
| CGGCCG | -58.24 | -63.93 | -111.10 | -65.22 | -71.99 | -119.54 |
| CGUACG | -57.56 | -61.10 | -104.91 | -51.71 | -57.00 | -115.70 |
| GACGUC | -54.65 | -59.34 | -105.17 | -52.86 | -58.19 | -107.59 |
| GCAUGC | -55.07 | -59.64 | -103.75 | -56.44 | -60.28 | -109.74 |
| GCCGGC | -66.87 | -73.67 | -108.99 | -62.45 | -70.93 | -113.26 |
| GCGAGC | -58.32 | -63.62 | -114.89 | -62.59 | -67.06 | -113.86 |
| GCGCGC | -60.15 | -66.84 | -110.91 | -67.13 | -72.36 | -105.04 |
| GCUAGC | -56.25 | -59.74 | -110.17 | -53.24 | -57.58 | -109.51 |
| GGAUCC | -56.07 | -60.62 | -103.90 | -54.23 | -60.00 | -109.53 |
| GGCGCC | -66.78 | -73.33 | -113.10 | -57.04 | -64.44 | -106.53 |
| GGGACC | -60.23 | -67.49 | -109.14 | -60.81 | -68.21 | -116.91 |
| GUGAAC | -49.08 | -53.59 | -101.81 | -51.81 | -56.04 | -103.83 |
| UCAUGA | -41.56 | -45.34 | -91.08 | -44.87 | -48.33 | -106.45 |
| UGAUCA | -44.03 | -46.71 | -95.68 | -41.22 | -45.39 | -93.19 |
| CCCGGG | -72.78 | -78.36 | -106.61 | -62.97 | -68.51 | -107.90 |
| ACCGCA | -56.95 | -61.04 | -104.20 | -53.91 | -58.77 | -98.17 |
| AGUUGC | -48.32 | -53.40 | -106.74 | -48.91 | -53.85 | -105.89 |
| AUGCGC | -51.70 | -56.45 | -104.83 | -51.92 | -55.94 | -98.03 |
| CCAACG | -57.91 | -59.62 | -99.74 | -55.59 | -59.16 | -110.96 |
| CGGACG | -62.96 | -67.27 | -110.66 | -58.72 | -64.88 | -119.88 |
| CGGUGC | -60.81 | -67.66 | -109.54 | -60.43 | -66.59 | -110.82 |
| CGUGCC | -62.15 | -67.66 | -106.80 | -54.70 | -61.05 | -110.04 |
| CGUUGA | -53.54 | -56.92 | -102.00 | -46.85 | -50.36 | -100.71 |
| CGUUGC | -54.73 | -59.09 | -108.26 | -54.47 | -59.56 | -106.74 |
| CGUUGU | -48.99 | -52.63 | -103.32 | -46.01 | -50.43 | -99.52 |
| GCCUGC | -61.01 | -65.79 | -109.63 | -61.75 | -66.32 | -112.00 |
| UGCGCA | -53.85 | -58.27 | -96.08 | -52.54 | -58.55 | -99.86 |
| UGUUGC | -45.27 | -49.87 | -101.17 | -40.58 | -45.04 | -105.17 |
| CAAUCG | -55.44 | -55.91 | -103.95 | -50.71 | -52.60 | -113.04 |
| CACGGC | -63.78 | -69.23 | -112.43 | -64.23 | -70.29 | -113.17 |
| CGAUUG | -41.53 | -44.26 | -95.11 | -46.28 | -49.61 | -105.80 |
| GCACCG | -61.08 | -65.72 | -105.54 | -56.60 | -62.43 | -114.51 |
| GGCACG | -63.44 | -68.31 | -110.40 | -55.99 | -62.21 | -103.35 |
| CCCAGGG | -71.46 | -78.49 | -116.65 | -79.74 | -87.54 | -122.44 |
| CAAAAAG | -57.15 | -57.68 | -114.87 | -59.01 | -59.13 | -112.21 |
| AAUACCG | -58.94 | -63.28 | -121.03 | -52.72 | -58.41 | -121.02 |
| AGCCGUG | -67.14 | -75.23 | -121.40 | -56.76 | -64.36 | -112.47 |
| AGCUUCA | -50.95 | -55.98 | -115.83 | -55.52 | -60.46 | -117.18 |
| GGACUUA | -52.77 | -58.84 | -118.24 | -57.97 | -63.73 | -117.38 |
| ACGUAUG | -57.55 | -62.51 | -116.81 | -48.21 | -55.31 | -112.52 |
| CACGGCU | -67.69 | -75.67 | -124.18 | -67.30 | -76.63 | -126.03 |

| | | | | | | |
|---|---|---|---|---|---|---|
| CAUACGU | -49.77 | -55.67 | -112.13 | -58.90 | -64.40 | -122.26 |
| UAAGUCC | -53.86 | -58.73 | -116.58 | -58.11 | -63.60 | -118.65 |
| UGAAGCU | -55.24 | -59.44 | -118.39 | -58.66 | -63.98 | -117.00 |
| AAAAAAAA | -50.25 | -51.53 | -120.00 | -52.83 | -54.15 | -121.06 |
| UAGAUCUA | -53.58 | -61.85 | -128.07 | -57.59 | -63.79 | -124.05 |
| UCUAUAGA | -54.59 | -60.87 | -126.34 | -59.26 | -65.42 | -123.46 |
| CAAAAAAG | -64.81 | -65.58 | -127.32 | -64.20 | -65.51 | -124.90 |
| CAAGCUUG | -68.26 | -73.73 | -126.27 | -67.58 | -72.99 | -140.23 |
| CAUCGAUG | -66.19 | -72.95 | -132.61 | -70.95 | -77.30 | -127.64 |
| CGAUAUCG | -68.01 | -74.66 | -129.09 | -62.44 | -71.18 | -141.49 |
| CGUCGACG | -70.61 | -81.53 | -137.91 | -73.23 | -83.65 | -142.72 |
| GAAGCUUC | -56.45 | -63.62 | -129.74 | -57.83 | -65.78 | -134.56 |
| GAUCGAUC | -56.79 | -65.87 | -126.65 | -69.51 | -78.13 | -138.53 |
| GAUGCAUC | -63.52 | -72.06 | -130.88 | -63.25 | -71.85 | -134.63 |
| GGAAUUCC | -66.49 | -74.65 | -128.80 | -64.96 | -73.82 | -135.34 |
| GGACGUCC | -75.15 | -87.44 | -134.95 | -75.50 | -87.33 | -136.22 |
| GGAGCUCC | -76.48 | -86.27 | -136.37 | -72.13 | -83.09 | -139.10 |
| GGUAUACC | -65.57 | -76.88 | -130.12 | -63.20 | -75.50 | -127.86 |
| GUACGUAC | -61.05 | -72.39 | -130.95 | -65.97 | -76.23 | -132.31 |
| GUAGCUAC | -64.87 | -73.68 | -126.09 | -62.69 | -71.74 | -130.65 |
| GUUGCAAC | -65.44 | -74.09 | -128.33 | -62.34 | -71.97 | -129.52 |
| AAGCGUAG | -68.77 | -77.62 | -132.60 | -68.74 | -75.88 | -134.36 |
| AAUCCAGU | -62.67 | -69.48 | -128.66 | -63.78 | -70.91 | -126.15 |
| ACAUAUGU | -53.32 | -59.84 | -124.30 | -57.01 | -64.17 | -130.66 |
| ACCUAGUC | -67.46 | -75.55 | -128.17 | -65.55 | -73.33 | -130.61 |
| ACGACCUC | -64.05 | -74.43 | -139.49 | -71.76 | -81.03 | -141.23 |
| AGAGAGAG | -71.73 | -79.14 | -129.96 | -66.41 | -73.77 | -134.27 |
| AGAGCUCU | -66.98 | -74.18 | -137.63 | -64.30 | -72.36 | -138.06 |
| AGCGUAAG | -67.84 | -76.42 | -124.54 | -63.05 | -72.87 | -133.10 |
| AGUCCUGA | -62.51 | -71.83 | -127.49 | -59.08 | -68.75 | -129.57 |
| AUGCGCAU | -55.03 | -66.86 | -122.00 | -65.18 | -73.63 | -133.12 |
| CACGGCUC | -76.74 | -87.39 | -137.85 | -78.10 | -88.42 | -141.26 |
| CCAUAUGG | -69.56 | -77.41 | -123.15 | -70.41 | -78.56 | -131.45 |
| CGCGUAUA | -58.08 | -66.34 | -129.07 | -59.32 | -67.70 | -132.24 |
| CGCUGUAA | -67.68 | -75.42 | -131.83 | -66.91 | -76.23 | -131.63 |
| CGUCGUCC | -72.03 | -83.22 | -132.46 | -75.13 | -84.71 | -137.39 |
| CUAGUGGA | -70.54 | -79.32 | -123.77 | -69.97 | -80.01 | -138.28 |
| CUCACGGC | -77.93 | -87.73 | -140.85 | -73.37 | -84.70 | -133.90 |
| CUGAGUCC | -69.81 | -78.83 | -134.91 | -74.32 | -83.49 | -134.59 |
| GAAUAUUC | -50.87 | -56.42 | -122.44 | -49.19 | -54.44 | -121.91 |
| GACUAGUC | -65.47 | -74.19 | -130.57 | -60.85 | -70.06 | -132.70 |
| GAGUACUC | -61.34 | -71.84 | -134.46 | -65.32 | -74.21 | -137.44 |
| GAUUAAUC | -52.21 | -58.74 | -124.74 | -56.44 | -61.98 | -124.94 |
| GCAUAUGC | -63.51 | -72.22 | -125.19 | -66.14 | -74.59 | -134.26 |
| GCCAGUUA | -64.79 | -74.13 | -130.87 | -67.75 | -76.17 | -130.59 |
| GGACCUCG | -77.44 | -86.86 | -141.72 | -73.80 | -84.19 | -137.22 |
| GGAGCACG | -79.97 | -89.70 | -135.21 | -71.46 | -82.72 | -140.64 |
| GGUGCCAA | -78.85 | -90.37 | -135.82 | -66.51 | -79.81 | -135.36 |
| GUCGAACA | -65.42 | -73.49 | -131.09 | -60.54 | -72.36 | -131.97 |
| GUCUAGAC | -65.84 | -74.78 | -129.61 | -63.81 | -72.94 | -129.31 |
| UAGGCCUA | -61.35 | -71.03 | -129.55 | -70.24 | -80.24 | -139.02 |
| UAUGCAUA | -49.94 | -57.31 | -123.86 | -54.10 | -61.91 | -119.75 |
| UGAGCUCA | -65.75 | -72.99 | -133.09 | -63.51 | -72.05 | -133.28 |
| AAUGUCGC | -60.25 | -69.58 | -132.40 | -65.59 | -74.29 | -130.15 |
| ACUGGAUU | -57.82 | -66.41 | -130.81 | -62.68 | -71.73 | -132.82 |
| CAACAGCA | -69.37 | -76.04 | -127.70 | -68.90 | -76.13 | -135.08 |
| CUACGCUU | -62.32 | -70.75 | -121.97 | -68.66 | -76.15 | -134.21 |
| CUUACGCU | -68.91 | -75.44 | -128.62 | -67.69 | -76.27 | -138.91 |

| | | | | | | |
|---|---|---|---|---|---|---|
| GACUAGGU | -64.12 | -73.28 | -130.67 | -69.30 | -79.00 | -136.27 |
| GAGCCGUG | -77.65 | -88.39 | -137.34 | -74.61 | -84.99 | -135.22 |
| GAGGUCGU | -66.02 | -76.76 | -135.00 | -72.59 | -83.60 | -143.53 |
| GCCGUGAG | -79.99 | -89.83 | -131.52 | -72.75 | -84.84 | -136.40 |
| GCGACAUU | -56.51 | -68.79 | -133.30 | -56.09 | -70.66 | -130.72 |
| UAACUGGC | -68.20 | -77.29 | -133.56 | -66.36 | -75.12 | -134.41 |
| UCCACUAG | -67.26 | -74.43 | -125.84 | -68.85 | -76.44 | -130.93 |
| UGCUGUUG | -63.81 | -72.55 | -122.51 | -63.31 | -71.64 | -128.24 |
| UGUUCGAC | -62.78 | -72.63 | -130.97 | -59.28 | -70.21 | -132.85 |
| UUACAGCG | -68.53 | -75.86 | -129.35 | -61.96 | -71.47 | -134.67 |
| UUGGCACC | -68.76 | -80.27 | -125.77 | -57.60 | -70.01 | -122.76 |
| CAAAAAAAG | -67.61 | -70.62 | -140.21 | -69.20 | -72.51 | -140.25 |
| CAAACAAAG | -72.23 | -78.19 | -139.57 | -74.07 | -79.77 | -146.72 |
| CAAAUAAAG | -67.55 | -72.14 | -137.06 | -68.49 | -73.48 | -142.58 |
| CAAAGAAAG | -73.69 | -78.75 | -139.42 | -72.62 | -78.65 | -145.49 |
| AAAAAAAAA | -56.09 | -58.35 | -128.54 | -58.13 | -60.30 | -131.33 |
| AUAACUGGC | -70.26 | -81.24 | -140.42 | -72.58 | -83.78 | -147.21 |
| AUCUAUCCG | -70.61 | -80.10 | -148.85 | -66.02 | -75.40 | -141.35 |
| CGCUGUUAC | -74.43 | -86.25 | -145.60 | -68.28 | -79.02 | -149.78 |
| GCCAGUUAA | -72.70 | -83.59 | -134.86 | -75.35 | -87.80 | -142.93 |
| CAACAGCAA | -69.90 | -79.07 | -142.35 | -72.67 | -82.51 | -140.15 |
| CAACAGCAU | -68.94 | -79.07 | -139.92 | -72.70 | -82.97 | -146.18 |
| CGCUGUUAG | -71.68 | -82.78 | -142.11 | -71.06 | -82.74 | -142.41 |
| CUAACAGCG | -80.56 | -88.99 | -149.92 | -76.97 | -86.63 | -146.18 |
| GUAACAGCG | -79.16 | -89.65 | -147.92 | -73.80 | -85.73 | -148.06 |
| UUAACUGGC | -64.19 | -75.62 | -138.96 | -72.69 | -83.96 | -136.27 |
| AUGAGCUCAU | -67.24 | -81.51 | -151.78 | -73.56 | -86.54 | -164.59 |
| GCGAAUUCGC | -77.46 | -91.76 | -159.32 | -85.76 | -99.18 | -163.64 |
| GCGAAAAGCG | -87.05 | -99.35 | -165.60 | -84.19 | -96.89 | -164.18 |
| AUCAAUCAUA | -66.31 | -75.36 | -147.94 | -49.01 | -62.34 | -139.72 |
| UUGUAGUCAU | -63.08 | -78.36 | -143.53 | -67.29 | -81.79 | -152.69 |
| GAAAUGAAAG | -75.34 | -84.60 | -154.92 | -73.93 | -83.30 | -159.52 |
| CCAACUUCUU | -68.56 | -79.80 | -144.28 | -80.38 | -89.77 | -160.43 |
| AUCGUCUGGA | -80.66 | -94.75 | -159.51 | -68.77 | -85.05 | -155.65 |
| AGCGUAAGUC | -80.29 | -97.02 | -157.59 | -75.17 | -92.17 | -163.51 |
| CGAUCUGCGA | -80.20 | -94.88 | -162.23 | -84.06 | -99.21 | -174.22 |
| UGGCGAGCAC | -77.52 | -97.24 | -153.47 | -89.98 | -107.46 | -159.20 |
| GAUGCGCUCG | -90.45 | -104.93 | -164.88 | -81.59 | -99.17 | -163.94 |
| GGGACCGCCU | -94.18 | -113.91 | -167.70 | -96.21 | -116.32 | -168.66 |
| AAAAAAAAAA | -61.41 | -65.14 | -143.21 | -62.83 | -67.62 | -144.90 |
| CGGCAAGCGC | -90.42 | -108.26 | -161.25 | -96.97 | -113.56 | -171.20 |
| UAGGUUAUAA | -67.82 | -79.94 | -149.82 | -66.27 | -79.51 | -150.29 |
| CGUACACAUGC | -83.38 | -103.69 | -165.09 | -88.04 | -106.46 | -172.28 |
| CCAUUGCUACC | -89.15 | -106.02 | -168.50 | -89.88 | -107.24 | -166.19 |
| CCAUCGCUACC | -93.54 | -111.37 | -165.52 | -94.53 | -112.01 | -164.73 |
| ACGUAUUAUGC | -75.63 | -92.29 | -164.56 | -62.83 | -81.22 | -165.63 |
| GCAUAAUACGU | -75.41 | -90.12 | -173.48 | -80.28 | -96.95 | -166.02 |
| CGCGAAUUCGCG | -107.17 | -125.92 | -194.70 | -94.74 | -118.59 | -190.12 |
| CUGACAAGUGUC | -81.85 | -104.77 | -173.18 | -94.81 | -115.69 | -194.53 |
| UUUUAUAAUAAA | -69.82 | -81.56 | -159.75 | -68.08 | -81.59 | -169.11 |
| AUUGGAUACAAA | -77.22 | -96.65 | -172.56 | -78.56 | -99.64 | -175.39 |
| CAACCAACCAAC | -98.43 | -117.35 | -188.22 | -96.76 | -116.04 | -189.12 |
| CUUCCUUCCUUC | -95.01 | -110.57 | -190.34 | -94.09 | -110.39 | -201.31 |
| GGAACAAGAUGC | -95.40 | -116.20 | -184.77 | -92.15 | -113.32 | -184.53 |
| GGAACUUGAUGC | -94.21 | -114.94 | -182.34 | -89.68 | -110.90 | -187.22 |
| AAUGGAUUACAA | -78.26 | -97.89 | -163.82 | -79.64 | -99.22 | -168.40 |
| GUCAGGAAUCUG | -88.63 | -109.17 | -180.96 | -92.30 | -113.57 | -177.61 |
| UUGUAAUCCAUU | -73.07 | -92.53 | -172.66 | -70.30 | -89.90 | -178.55 |

| | | | | | | |
|---|---|---|---|---|---|---|
| UUUGUAUCCAAU | -76.20 | -94.16 | -176.68 | -68.11 | -89.00 | -167.24 |
| CGCAUGGGUACGC | -104.69 | -135.73 | -205.32 | -114.30 | -143.37 | -203.54 |
| AGCCUAACUCAGC | -99.10 | -122.02 | -196.13 | -93.83 | -115.04 | -198.31 |
| CAUAUGGCCAUAUG | -99.28 | -125.56 | -210.31 | -99.95 | -126.97 | -200.17 |
| GUAUACCGGUAUAC | -96.48 | -126.32 | -207.71 | -93.88 | -123.79 | -214.87 |
| ACAUUAUUAUUACA | -73.93 | -95.24 | -181.90 | -81.61 | -102.69 | -200.78 |
| UACUAACAUUAACUA | -74.08 | -101.70 | -206.94 | -81.09 | -108.46 | -213.88 |
| AUACUUACUGAUUAG | -80.25 | -106.69 | -212.69 | -85.78 | -113.83 | -216.75 |
| GUACACUGUCUUAUA | -87.65 | -116.41 | -210.09 | -95.70 | -128.28 | -221.71 |
| GUAUGAGAGACUUUA | -94.28 | -125.41 | -217.25 | -90.53 | -122.51 | -215.78 |
| UUCUACCUAUGUGAU | -97.17 | -125.97 | -207.25 | -90.43 | -119.08 | -213.80 |
| AGUAGUAAUCACACC | -94.78 | -128.54 | -219.54 | -90.42 | -127.00 | -208.89 |
| AUCGUCUCGGUAUAA | -97.63 | -128.16 | -224.87 | -98.93 | -130.72 | -219.13 |
| ACGACAGGUUUACCA | -105.52 | -139.00 | -211.77 | -101.46 | -137.71 | -224.86 |
| CUUUCAUGUCCGCAU | -99.12 | -129.90 | -215.10 | -108.56 | -137.92 | -220.44 |
| UGGAUGUGUGAACAC | -100.09 | -137.99 | -219.47 | -91.38 | -126.98 | -218.11 |
| ACCCCGCAAUACAUG | -101.56 | -134.24 | -214.24 | -106.11 | -137.59 | -217.32 |
| GCAGUGGAUGUGAGA | -68.09 | -109.85 | -204.55 | -116.81 | -151.21 | -229.77 |
| GGUCCUUACUUGGUG | -104.85 | -137.96 | -227.16 | -110.83 | -144.02 | -239.46 |
| CGCCUCAUGCUCAUC | -102.75 | -131.81 | -221.15 | -111.13 | -143.28 | -215.98 |
| AAAUAGCCGGGCCGC | -124.22 | -160.16 | -243.51 | -116.11 | -151.79 | -241.06 |
| CCAGCCAGUCUCUCC | -120.15 | -152.36 | -231.63 | -115.93 | -148.52 | -232.06 |
| GACGACAAGACCGCG | -122.75 | -156.39 | -241.17 | -121.52 | -156.42 | -232.06 |
| CAGCCUCGUCGCAGC | -121.10 | -155.10 | -239.37 | -121.63 | -157.29 | -230.06 |
| CUCGCGGUCGAAGCG | -115.58 | -154.29 | -242.78 | -128.76 | -161.15 | -238.05 |
| GCGUCGGUCCGGGCU | -104.63 | -145.53 | -225.94 | -132.27 | -172.90 | -246.97 |
| CAACUUGAUAUUAUUA | -82.91 | -111.17 | -228.30 | -89.87 | -119.27 | -231.03 |
| CAUAUUGGCCAAUAUG | -108.29 | -141.29 | -237.68 | -104.56 | -136.13 | -231.03 |
| GUAUAACCGGUUAUAC | -99.38 | -136.78 | -232.53 | -102.01 | -136.43 | -237.50 |
| CGCGUACGCGUACGCG | -130.93 | -169.78 | -245.48 | -114.70 | -161.07 | -251.19 |
| CAACUUGAUAUUAAUA | -83.49 | -115.07 | -219.52 | -78.19 | -108.00 | -218.75 |
| UAUGUAUAUUUUGUAAUCAG | -94.66 | -141.69 | -279.25 | -92.10 | -140.94 | -268.22 |
| UUCAAGUUAAACAUUCUAUC | -108.50 | -153.42 | -276.46 | -82.02 | -129.51 | -277.33 |
| UGAUUCUACCUAUGUGAUUU | -87.06 | -135.07 | -272.63 | -107.81 | -157.75 | -284.85 |
| GAGAUUGUUUCCCUUUCAAA | -103.21 | -153.08 | -282.28 | -113.83 | -164.71 | -268.63 |
| AUGCAAUGCUACAUAUUCGC | -112.18 | -162.64 | -281.98 | -97.55 | -149.73 | -288.75 |
| CCACUAUACCAUCUAUGUAC | -93.97 | -142.75 | -254.05 | -114.74 | -166.22 | -273.03 |
| CCAUCAUUGUGUCUACCUCA | -105.81 | -157.93 | -274.78 | -98.99 | -167.99 | -291.28 |
| CGGGACCAACUAAAGGAAAU | -136.72 | -190.02 | -295.65 | -128.49 | -184.14 | -297.30 |
| UAGUGGCGAUUAGAUUCUGC | -76.66 | -133.29 | -271.38 | -125.03 | -181.82 | -296.89 |
| AGCUGCAGUGGAUGUGAGAA | -133.47 | -189.75 | -286.75 | -122.52 | -183.03 | -285.59 |
| UACUUCCAGUGCUCAGCGUA | -117.19 | -172.12 | -284.84 | -130.87 | -183.15 | -297.08 |
| CAGUGAGACAGCAAUGGUCG | -125.68 | -182.72 | -299.38 | -131.97 | -186.26 | -299.29 |
| CGAGCUUAUCCCUAUCCCUC | -133.45 | -184.17 | -304.03 | -121.03 | -175.95 | -296.27 |
| CGUACUAGCGUUGGUCAUGG | -127.26 | -184.54 | -288.25 | -134.46 | -192.89 | -298.15 |
| AAGGCGAGUCAGGCUCAGUG | -139.45 | -195.47 | -297.64 | -138.07 | -195.80 | -307.75 |
| ACCGACGACGCUGAUCCGAU | -137.58 | -194.03 | -292.49 | -136.27 | -190.40 | -300.46 |
| AGCAGUCCGCCACACCCUGA | -133.58 | -190.98 | -288.39 | -131.15 | -191.47 | -292.95 |
| CAGCCUCGUUCGCACAGCCC | -155.02 | -210.94 | -311.72 | -125.79 | -191.20 | -291.00 |
| GUGGUGGGCCGUGCGCUCUG | -141.34 | -208.12 | -299.43 | -158.08 | -221.89 | -298.99 |
| GUCCACGCCCGGUGCGACGG | -144.81 | -205.67 | -309.10 | -136.61 | -193.98 | -306.57 |
| GAUAUAGCAAAAUUCUAAGUUAAUA | -116.37 | -185.18 | -347.77 | -108.96 | -179.12 | -346.87 |
| AUAACUUUACGUGUGUGACCUAUUA | -129.39 | -207.45 | -355.23 | -59.99 | -133.96 | -295.46 |
| GUUCUAUACUCUUGAAGUUGAUUAC | -112.67 | -182.08 | -345.55 | -122.28 | -195.62 | -342.24 |
| CCCUGCACUUUAACUGAAUUGUUUA | -126.83 | -194.73 | -346.57 | -89.28 | -166.20 | -327.62 |
| UAACCAUACUGAAUACCUUUUGACG | -89.68 | -158.12 | -299.31 | -110.33 | -182.97 | -348.61 |
| UCCACACGGUAGUAAAAUUAGGCUU | -123.55 | -197.65 | -348.78 | -132.53 | -208.51 | -368.63 |
| UUCCAAAAGGAGUUAUGAGUUGCGA | -127.74 | -197.95 | -334.37 | -108.23 | -184.24 | -321.96 |

| | | | | | | |
|---|---|---|---|---|---|---|
| AAUAUCUCUCAUGCGCCAAGCUACA | -129.93 | -203.35 | -345.23 | -125.61 | -203.77 | -351.91 |
| UAGUAUAUCGCAGCAUCAUACAGGC | -116.65 | -193.92 | -357.01 | -98.74 | -180.91 | -340.36 |
| UGGAUUCUACUCAACCUUAGUCUGG | -143.26 | -219.99 | -365.04 | -132.20 | -208.16 | -353.03 |
| CGGAAUCCAUGUUACUUCGGCUAUC | -124.33 | -202.77 | -346.54 | -104.43 | -185.86 | -331.51 |
| CUGGUCUGGAUCUGAGAACUUCAGG | -140.92 | -218.59 | -360.23 | -124.36 | -210.30 | -363.12 |
| ACAGCGAAUGGACCUACGUGGCCUU | -125.89 | -213.38 | -340.92 | -170.12 | -252.55 | -367.99 |
| AGCAAGUCGAGCAGGGCCUACGUUU | -157.32 | -241.27 | -361.81 | -166.05 | -247.60 | -381.69 |
| GCGAGCGACAGGUUACUUGGCUGAU | -147.98 | -231.65 | -351.34 | -144.39 | -231.50 | -370.64 |
| AAAGGUGUCGCGGAGAGUCGUGCUG | -142.19 | -224.86 | -359.28 | -104.41 | -187.47 | -352.01 |
| AUGGGUGGGAGCCUCGGUAGCAGCC | -152.35 | -238.05 | -359.92 | -169.29 | -259.87 | -380.06 |
| CAGUGGGCUCCUGGGCGUGCUGGUC | -173.63 | -262.37 | -388.63 | -167.71 | -263.50 | -383.71 |
| GCCAACUCCGUCGCCGUUCGUGCGC | -138.17 | -221.73 | -355.09 | -127.46 | -214.89 | -356.20 |
| ACGGGUCCCCGCACCGCACCGCCAG | -151.56 | -237.63 | -329.53 | -185.91 | -278.92 | -378.46 |
| UUAUGUAUUAAGUUAUAUAGUAGUAGUAGU | -111.19 | -216.09 | -398.43 | -58.60 | -160.91 | -379.77 |
| AUUGAUAUCCUUUUCUAUUCAUCUUUCAUU | -75.28 | -176.68 | -390.20 | -96.05 | -198.64 | -401.90 |
| AAAGUACAUCAACAUAGAGAAUUGCAUUUC | -119.50 | -221.83 | -408.80 | -104.02 | -201.01 | -391.93 |
| CUUAAGAUAUGAGAACUUCAACUAAUGUGU | -102.37 | -209.70 | -406.59 | -140.29 | -237.50 | -401.85 |
| CUCAACUUGCGGUAAAUAAAUCGCUUAAUC | -111.82 | -215.42 | -394.29 | -61.71 | -163.18 | -392.06 |
| UAUUGAGAACAAGUGUCCGAUUAGCAGAAA | -134.90 | -243.63 | -423.77 | -150.57 | -255.72 | -427.25 |
| GUCAUACGACUGAGUGCAACAUUGUUCAAA | -131.25 | -238.11 | -399.79 | -133.53 | -246.97 | -408.11 |
| AACCUGCAACAUGGAGUUUUUGUCUCAUGC | -131.23 | -232.85 | -404.41 | -105.53 | -210.90 | -410.10 |
| CCGUGCGGUGUGUACGUUUUAUUCAUCAUA | -140.26 | -255.20 | -424.95 | -145.70 | -255.31 | -421.22 |
| GUUCACGUCCGAAAGCUCGAAAAAGGAUAC | -82.69 | -191.73 | -386.97 | -125.89 | -235.22 | -420.21 |
| AGUCUGGUCUGGAUCUGAGAACUUCAGGCU | -160.61 | -271.51 | -433.84 | -95.65 | -209.99 | -416.25 |
| UCGGAGAAAUCACUGAGCUGCCUGAGAAGA | -109.41 | -218.64 | -405.89 | -127.23 | -242.81 | -414.33 |
| CUUCAACGGAUCAGGUAGGACUGUGGUGGG | -86.16 | -225.85 | -430.99 | -178.00 | -290.83 | -452.39 |
| ACGCCCACAGGAUUAGGCUGGCCCACAUUG | -174.68 | -293.20 | -425.93 | -169.65 | -280.19 | -446.37 |
| GUUAUUCCGCAGUCCGAUGGCAGCAGGCUC | -166.84 | -279.83 | -437.61 | -105.96 | -222.75 | -425.83 |
| UCAGUAGGCGUGACGCAGAGCUGGCGAUGG | -178.46 | -298.50 | -448.35 | -179.31 | -298.44 | -431.73 |
| CGCGCCACGUGUGAUCUACAGCCGUUCGGC | -129.59 | -251.71 | -422.34 | -162.95 | -281.29 | -443.36 |
| GACCUGACGUGGACCGCUCCUGGGCGUGGU | -202.89 | -319.57 | -444.48 | -144.14 | -262.04 | -419.36 |
| GCCCCUCCACUGGCCGACGGCAGCAGGCUC | -164.13 | -291.04 | -435.88 | -181.95 | -304.38 | -460.28 |
| CGCCGCUGCCGACUGGAGGAGCGCGGGACG | -162.89 | -296.15 | -456.55 | -121.84 | -240.37 | -427.20 |

[a] All duplexes consist of the denoted RNA strand and its complementary DNA strand.

**Table S2.** Corrected Thermodynamic Parameters for Natural RNA/DNA Duplexes

| RNA Sequence[a] | MMPBSA | | | | MMGBSA | | | |
|---|---|---|---|---|---|---|---|---|
| | $\Delta H°$ (kcal/mol) | $\Delta S°$ (cal/mol/K) | $\Delta G°_{37}$ (kcal/mol) | $T_m$ (°C) | $\Delta H°$ (kcal/mol) | $\Delta S°$ (cal/mol/K) | $\Delta G°_{37}$ (kcal/mol) | $T_m$ (°C) |
| CCGG | -34.57 | -100.79 | -3.32 | 10.54 | -34.64 | -99.45 | -3.81 | 14.31 |
| CGCG | -33.61 | -96.68 | -3.64 | 12.34 | -33.64 | -96.23 | -3.81 | 13.69 |
| GCGC | -37.05 | -109.42 | -3.14 | 10.85 | -36.96 | -108.01 | -3.48 | 13.21 |
| AGCCG | -39.83 | -116.20 | -3.81 | 17.06 | -39.79 | -115.06 | -4.12 | 19.19 |
| CACAG | -36.87 | -107.17 | -3.64 | 14.36 | -36.84 | -106.30 | -3.89 | 16.14 |
| CGUGC | -42.32 | -121.19 | -4.75 | 24.36 | -42.34 | -120.77 | -4.91 | 25.42 |
| GCACG | -43.51 | -129.41 | -3.40 | 16.05 | -43.32 | -127.55 | -3.78 | 18.38 |
| CGGCU | -41.18 | -118.29 | -4.51 | 22.39 | -41.27 | -117.42 | -4.87 | 24.91 |
| GGUGG | -38.50 | -109.20 | -4.65 | 22.41 | -38.78 | -108.24 | -5.22 | 26.77 |
| CCGCGG | -53.59 | -148.71 | -7.49 | 42.54 | -53.72 | -149.90 | -7.25 | 41.07 |
| CGAUCG | -42.29 | -120.99 | -4.78 | 24.55 | -42.22 | -121.03 | -4.70 | 23.98 |
| CGCGCG | -47.59 | -131.79 | -6.73 | 38.20 | -47.83 | -132.47 | -6.76 | 38.39 |
| CGGCCG | -46.40 | -127.47 | -6.88 | 39.23 | -46.80 | -127.92 | -7.15 | 41.01 |
| CGUACG | -49.98 | -142.54 | -5.79 | 32.35 | -49.88 | -142.77 | -5.62 | 31.28 |
| GACGUC | -49.02 | -141.99 | -5.01 | 27.54 | -48.93 | -141.14 | -5.18 | 28.52 |
| GCAUGC | -47.18 | -134.63 | -5.44 | 29.87 | -47.15 | -134.53 | -5.45 | 29.91 |
| GCCGGC | -56.14 | -157.13 | -7.43 | 41.91 | -56.31 | -157.51 | -7.48 | 42.20 |
| GCGAGC | -53.36 | -151.46 | -6.40 | 36.12 | -53.27 | -151.92 | -6.17 | 34.80 |
| GCGCGC | -55.11 | -155.27 | -6.97 | 39.37 | -55.10 | -155.93 | -6.76 | 38.17 |
| GCUAGC | -50.76 | -146.02 | -5.49 | 30.65 | -50.51 | -146.25 | -5.17 | 28.76 |
| GGAUCC | -43.22 | -123.36 | -4.98 | 26.14 | -43.36 | -122.62 | -5.34 | 28.61 |
| GGCGCC | -51.14 | -142.41 | -6.99 | 39.68 | -51.44 | -142.51 | -7.26 | 41.35 |
| GGGACC | -49.00 | -137.71 | -6.31 | 35.45 | -49.32 | -137.07 | -6.83 | 38.75 |
| GUGAAC | -45.87 | -133.10 | -4.60 | 24.38 | -45.78 | -132.19 | -4.80 | 25.59 |
| UCAUGA | -40.44 | -119.43 | -3.42 | 14.74 | -40.30 | -118.02 | -3.72 | 16.63 |
| UGAUCA | -36.60 | -108.32 | -3.02 | 9.73 | -36.56 | -106.60 | -3.51 | 13.20 |
| CCCGGG | -51.28 | -138.49 | -8.35 | 48.28 | -51.75 | -140.01 | -8.34 | 48.12 |
| ACCGCA | -46.51 | -131.86 | -5.64 | 31.04 | -46.65 | -131.50 | -5.88 | 32.63 |
| AGUUGC | -47.84 | -140.78 | -4.19 | 22.44 | -47.56 | -139.67 | -4.27 | 22.79 |
| AUGCGC | -45.59 | -129.73 | -5.37 | 29.19 | -45.59 | -129.72 | -5.38 | 29.23 |
| CCAACG | -46.59 | -133.55 | -5.19 | 28.20 | -46.54 | -133.26 | -5.23 | 28.45 |
| CGGACG | -49.48 | -138.51 | -6.54 | 36.93 | -49.62 | -138.95 | -6.55 | 36.98 |
| CGGUGC | -49.10 | -136.89 | -6.67 | 37.74 | -49.36 | -137.12 | -6.85 | 38.90 |
| CGUGCC | -48.69 | -136.20 | -6.47 | 36.51 | -48.92 | -136.32 | -6.66 | 37.68 |
| CGUUGA | -44.87 | -128.24 | -5.12 | 27.41 | -44.81 | -128.23 | -5.06 | 27.04 |
| CGUUGC | -49.62 | -140.95 | -5.93 | 33.15 | -49.55 | -141.35 | -5.73 | 31.94 |
| CGUUGU | -46.28 | -133.91 | -4.77 | 25.49 | -46.08 | -133.70 | -4.63 | 24.60 |
| GCCUGC | -46.54 | -129.05 | -6.54 | 36.93 | -46.80 | -129.57 | -6.63 | 37.56 |
| UGCGCA | -47.84 | -137.55 | -5.20 | 28.50 | -47.84 | -136.85 | -5.41 | 29.79 |
| UGUUGC | -45.91 | -133.83 | -4.42 | 23.26 | -45.77 | -132.87 | -4.58 | 24.20 |
| CAAUCG | -46.22 | -133.27 | -4.91 | 26.34 | -45.90 | -133.81 | -4.42 | 23.26 |
| CACGGC | -50.86 | -141.91 | -6.87 | 38.95 | -51.08 | -142.27 | -6.98 | 39.60 |
| CGAUUG | -43.92 | -127.06 | -4.53 | 23.37 | -43.69 | -127.09 | -4.29 | 21.77 |
| GCACCG | -49.47 | -139.52 | -6.21 | 34.89 | -49.59 | -139.52 | -6.34 | 35.67 |
| GGCACG | -50.03 | -140.72 | -6.41 | 36.13 | -50.18 | -140.83 | -6.52 | 36.83 |
| CCCAGGG | -55.64 | -148.48 | -9.61 | 55.06 | -56.25 | -150.50 | -9.59 | 54.71 |
| CAAAAAG | -51.13 | -148.62 | -5.06 | 28.20 | -50.68 | -149.00 | -4.49 | 24.89 |
| AAUACCG | -55.32 | -160.62 | -5.53 | 31.34 | -55.02 | -160.22 | -5.36 | 30.39 |
| AGCCGUG | -58.96 | -166.12 | -7.46 | 41.84 | -58.97 | -166.68 | -7.30 | 40.96 |
| AGCUUCA | -49.35 | -142.76 | -5.10 | 28.13 | -49.20 | -142.26 | -5.10 | 28.08 |
| GGACUUA | -53.03 | -153.36 | -5.49 | 30.92 | -52.86 | -152.81 | -5.49 | 30.88 |
| ACGUAUG | -54.72 | -159.68 | -5.21 | 29.59 | -54.39 | -159.02 | -5.10 | 28.92 |
| CACGGCU | -54.64 | -153.65 | -7.01 | 39.60 | -54.91 | -153.20 | -7.42 | 41.96 |

| | | | | | | | | |
|---|---|---|---|---|---|---|---|---|
| CAUACGU | -52.23 | -149.00 | -6.04 | 33.97 | -52.13 | -149.15 | -5.89 | 33.13 |
| UAAGUCC | -48.81 | -142.31 | -4.70 | 25.64 | -48.68 | -141.14 | -4.93 | 27.01 |
| UGAAGCU | -50.41 | -146.24 | -5.08 | 28.18 | -50.17 | -145.85 | -4.96 | 27.47 |
| AAAAAAAA | -53.61 | -158.28 | -4.55 | 25.81 | -52.95 | -158.29 | -3.88 | 22.11 |
| UAGAUCUA | -53.27 | -158.15 | -4.24 | 24.11 | -52.91 | -156.40 | -4.43 | 25.02 |
| UCUAUAGA | -53.79 | -155.52 | -5.58 | 31.48 | -53.55 | -155.25 | -5.42 | 30.59 |
| CAAAAAAG | -56.81 | -161.85 | -6.64 | 37.47 | -56.40 | -163.45 | -5.73 | 32.54 |
| CAAGCUUG | -61.79 | -173.35 | -8.05 | 44.71 | -61.59 | -175.24 | -7.27 | 40.63 |
| CAUCGAUG | -56.79 | -157.83 | -7.86 | 44.31 | -56.90 | -159.04 | -7.59 | 42.77 |
| CGAUAUCG | -57.72 | -162.28 | -7.42 | 41.70 | -57.69 | -163.23 | -7.09 | 39.88 |
| CGUCGACG | -69.68 | -194.58 | -9.36 | 49.99 | -69.71 | -195.91 | -8.98 | 48.14 |
| GAAGCUUC | -59.76 | -171.68 | -6.54 | 36.90 | -59.45 | -172.05 | -6.11 | 34.68 |
| GAUCGAUC | -60.15 | -172.85 | -6.56 | 37.03 | -59.97 | -172.58 | -6.47 | 36.56 |
| GAUGCAUC | -57.95 | -164.97 | -6.81 | 38.35 | -57.96 | -164.67 | -6.91 | 38.93 |
| GGAAUUCC | -55.96 | -157.59 | -7.10 | 40.07 | -56.05 | -157.83 | -7.12 | 40.16 |
| GGACGUCC | -62.51 | -173.85 | -8.61 | 47.55 | -62.91 | -173.78 | -9.04 | 49.74 |
| GGAGCUCC | -58.70 | -160.70 | -8.88 | 49.82 | -59.11 | -161.80 | -8.95 | 50.12 |
| GGUAUACC | -59.07 | -168.02 | -6.98 | 39.27 | -59.25 | -167.02 | -7.48 | 41.90 |
| GUACGUAC | -62.93 | -179.97 | -7.14 | 39.88 | -62.96 | -179.11 | -7.43 | 41.38 |
| GUAGCUAC | -64.21 | -184.68 | -6.96 | 38.95 | -63.96 | -184.56 | -6.74 | 37.90 |
| GUUGCAAC | -61.38 | -174.87 | -7.17 | 40.11 | -61.38 | -174.52 | -7.28 | 40.68 |
| AAGCGUAG | -64.90 | -186.32 | -7.14 | 39.83 | -64.59 | -186.63 | -6.73 | 37.83 |
| AAUCCAGU | -56.93 | -163.41 | -6.27 | 35.47 | -56.76 | -163.32 | -6.13 | 34.67 |
| ACAUAUGU | -55.55 | -161.04 | -5.62 | 31.88 | -55.27 | -160.63 | -5.48 | 31.08 |
| ACCUAGUC | -59.09 | -168.63 | -6.81 | 38.34 | -59.04 | -168.41 | -6.83 | 38.45 |
| ACGACCUC | -66.43 | -189.80 | -7.59 | 41.91 | -66.35 | -189.45 | -7.62 | 42.05 |
| AGAGAGAG | -57.63 | -161.26 | -7.64 | 42.95 | -57.61 | -162.50 | -7.24 | 40.72 |
| AGAGCUCU | -61.51 | -176.61 | -6.76 | 38.01 | -61.26 | -176.70 | -6.48 | 36.60 |
| AGCGUAAG | -63.30 | -181.14 | -7.14 | 39.90 | -63.07 | -181.37 | -6.85 | 38.42 |
| AGUCCUGA | -55.99 | -157.51 | -7.16 | 40.38 | -56.15 | -157.50 | -7.33 | 41.35 |
| AUGCGCAU | -60.69 | -173.07 | -7.04 | 39.51 | -60.64 | -172.90 | -7.04 | 39.50 |
| CACGGCUC | -66.17 | -183.90 | -9.16 | 49.70 | -66.40 | -184.72 | -9.14 | 49.56 |
| CCAUAUGG | -55.74 | -153.49 | -8.16 | 46.19 | -56.04 | -154.54 | -8.13 | 45.97 |
| CGCGUAUA | -61.47 | -173.71 | -7.62 | 42.48 | -61.46 | -174.14 | -7.47 | 41.69 |
| CGCUGUAA | -59.85 | -170.51 | -7.00 | 39.30 | -59.78 | -170.53 | -6.92 | 38.88 |
| CGUCGUCC | -64.41 | -181.34 | -8.19 | 45.08 | -64.55 | -181.36 | -8.33 | 45.77 |
| CUAGUGGA | -60.05 | -167.85 | -8.01 | 44.71 | -60.17 | -168.56 | -7.92 | 44.19 |
| CUCACGGC | -66.04 | -182.82 | -9.37 | 50.78 | -66.32 | -183.78 | -9.35 | 50.62 |
| CUGAGUCC | -62.17 | -174.34 | -8.12 | 45.02 | -62.26 | -174.96 | -8.02 | 44.47 |
| GAAUAUUC | -53.39 | -155.59 | -5.16 | 29.10 | -53.07 | -155.11 | -4.98 | 28.08 |
| GACUAGUC | -58.35 | -166.31 | -6.79 | 38.27 | -58.37 | -165.92 | -6.93 | 39.02 |
| GAGUACUC | -59.52 | -171.35 | -6.40 | 36.19 | -59.45 | -170.44 | -6.62 | 37.33 |
| GAUUAAUC | -51.69 | -150.28 | -5.10 | 28.51 | -51.44 | -149.68 | -5.04 | 28.15 |
| GCAUAUGC | -62.92 | -179.35 | -7.32 | 40.82 | -62.77 | -179.61 | -7.10 | 39.68 |
| GCCAGUUA | -60.02 | -169.86 | -7.37 | 41.25 | -60.08 | -169.86 | -7.42 | 41.54 |
| GGACCUCG | -63.30 | -174.03 | -9.35 | 51.31 | -63.61 | -175.37 | -9.25 | 50.69 |
| GGAGCACG | -63.73 | -175.43 | -9.35 | 51.20 | -64.02 | -176.79 | -9.22 | 50.44 |
| GGUGCCAA | -61.05 | -169.80 | -8.41 | 46.74 | -61.49 | -169.56 | -8.93 | 49.44 |
| GUCGAACA | -60.34 | -171.34 | -7.22 | 40.45 | -60.42 | -170.91 | -7.44 | 41.61 |
| GUCUAGAC | -59.09 | -168.10 | -6.97 | 39.21 | -59.09 | -167.88 | -7.05 | 39.62 |
| UAGGCCUA | -66.33 | -192.69 | -6.60 | 37.18 | -66.23 | -190.89 | -7.06 | 39.34 |
| UAUGCAUA | -58.31 | -170.32 | -5.52 | 31.56 | -57.97 | -169.52 | -5.42 | 31.02 |
| UGAGCUCA | -55.73 | -158.37 | -6.64 | 37.46 | -55.73 | -158.33 | -6.65 | 37.54 |
| AAUGUCGC | -63.83 | -183.01 | -7.10 | 39.65 | -63.60 | -183.11 | -6.83 | 38.35 |
| ACUGGAUU | -55.17 | -155.35 | -7.01 | 39.61 | -55.30 | -155.45 | -7.11 | 40.17 |
| CAACAGCA | -60.93 | -170.43 | -8.10 | 45.06 | -61.00 | -171.34 | -7.89 | 43.92 |
| CUACGCUU | -63.13 | -180.32 | -7.23 | 40.35 | -62.93 | -180.64 | -6.93 | 38.83 |
| CUUACGCU | -61.90 | -176.25 | -7.26 | 40.56 | -61.74 | -176.62 | -6.99 | 39.20 |

| | | | | | | | | |
|---|---|---|---|---|---|---|---|---|
| GACUAGGU | -59.68 | -168.59 | -7.42 | 41.55 | -59.78 | -168.54 | -7.53 | 42.14 |
| GAGCCGUG | -65.30 | -179.88 | -9.54 | 51.83 | -65.57 | -181.33 | -9.36 | 50.82 |
| GAGGUCGU | -62.99 | -175.63 | -8.54 | 47.09 | -63.15 | -176.43 | -8.45 | 46.61 |
| GCCGUGAG | -68.40 | -190.60 | -9.32 | 50.03 | -68.57 | -191.50 | -9.21 | 49.45 |
| GCGACAUU | -65.17 | -187.77 | -6.96 | 38.91 | -64.99 | -187.09 | -7.00 | 39.11 |
| UAACUGGC | -61.75 | -175.66 | -7.29 | 40.75 | -61.70 | -175.67 | -7.24 | 40.49 |
| UCCACUAG | -55.16 | -153.86 | -7.46 | 42.20 | -55.38 | -154.28 | -7.55 | 42.71 |
| UGCUGUUG | -55.29 | -154.43 | -7.41 | 41.91 | -55.54 | -154.61 | -7.61 | 43.00 |
| UGUUCGAC | -60.76 | -173.72 | -6.90 | 38.79 | -60.74 | -173.14 | -7.07 | 39.66 |
| UUACAGCG | -62.30 | -176.55 | -7.57 | 42.14 | -62.23 | -176.98 | -7.37 | 41.09 |
| UUGGCACC | -58.47 | -162.40 | -8.13 | 45.59 | -58.96 | -162.02 | -8.73 | 48.90 |
| CAAAAAAAG | -65.04 | -185.94 | -7.40 | 41.04 | -64.52 | -187.55 | -6.38 | 36.12 |
| CAAACAAAG | -68.51 | -195.70 | -7.85 | 42.94 | -68.19 | -196.40 | -7.31 | 40.45 |
| CAAAUAAAG | -62.79 | -180.07 | -6.97 | 39.04 | -62.40 | -180.86 | -6.34 | 35.90 |
| CAAAGAAAG | -67.16 | -189.33 | -8.47 | 46.07 | -66.83 | -191.34 | -7.52 | 41.52 |
| AAAAAAAAA | -59.95 | -175.05 | -5.69 | 32.56 | -59.29 | -175.61 | -4.85 | 28.30 |
| AUAACUGGC | -67.20 | -190.18 | -8.25 | 44.99 | -67.22 | -190.28 | -8.24 | 44.94 |
| AUCUAUCCG | -64.97 | -184.72 | -7.71 | 42.59 | -64.85 | -185.05 | -7.49 | 41.50 |
| CGCUGUUAC | -74.54 | -210.62 | -9.25 | 48.61 | -74.46 | -211.27 | -8.97 | 47.36 |
| GCCAGUUAA | -66.73 | -189.46 | -7.99 | 43.82 | -66.76 | -189.18 | -8.11 | 44.40 |
| CAACAGCAA | -67.71 | -189.65 | -8.92 | 48.21 | -67.82 | -190.32 | -8.82 | 47.71 |
| CAACAGCAU | -68.86 | -195.85 | -8.14 | 44.30 | -68.74 | -196.07 | -7.95 | 43.42 |
| CGCUGUUAG | -71.06 | -199.95 | -9.07 | 48.38 | -70.98 | -201.03 | -8.66 | 46.46 |
| CUAACAGCG | -72.20 | -201.93 | -9.60 | 50.64 | -72.12 | -203.60 | -9.01 | 47.90 |
| GUAACAGCG | -73.94 | -208.63 | -9.26 | 48.75 | -73.87 | -209.36 | -8.97 | 47.46 |
| UUAACUGGC | -63.70 | -179.47 | -8.06 | 44.50 | -63.83 | -179.47 | -8.20 | 45.18 |
| AUGAGCUCAU | -71.53 | -202.86 | -8.65 | 46.31 | -71.51 | -202.90 | -8.61 | 46.17 |
| GCGAAUUCGC | -80.54 | -225.96 | -10.49 | 52.90 | -80.55 | -226.96 | -10.19 | 51.63 |
| GCGAAAAGCG | -84.76 | -237.79 | -11.04 | 54.30 | -84.56 | -239.75 | -10.24 | 51.08 |
| AUCAAUCAUA | -68.74 | -199.84 | -6.79 | 38.06 | -68.45 | -198.71 | -6.85 | 38.32 |
| UUGUAGUCAU | -70.91 | -203.68 | -7.77 | 42.36 | -70.88 | -202.50 | -8.10 | 43.89 |
| GAAAUGAAAG | -72.45 | -206.68 | -8.37 | 44.96 | -72.17 | -207.25 | -7.92 | 42.95 |
| CCAACUUCUU | -74.82 | -212.08 | -9.07 | 47.78 | -74.68 | -212.69 | -8.75 | 46.34 |
| AUCGUCUGGA | -71.01 | -197.78 | -9.70 | 51.36 | -71.41 | -197.75 | -10.11 | 53.20 |
| AGCGUAAGUC | -76.03 | -214.99 | -9.39 | 48.96 | -76.13 | -214.79 | -9.54 | 49.64 |
| CGAUCUGCGA | -71.08 | -196.96 | -10.03 | 52.90 | -71.53 | -197.19 | -10.41 | 54.62 |
| UGGCGAGCAC | -77.57 | -215.74 | -10.69 | 54.42 | -78.04 | -215.62 | -11.20 | 56.58 |
| GAUGCGCUCG | -81.17 | -226.62 | -10.91 | 54.55 | -81.33 | -227.50 | -10.80 | 54.05 |
| GGGACCGCCU | -85.61 | -235.40 | -12.64 | 60.67 | -86.36 | -235.51 | -13.35 | 63.45 |
| AAAAAAAAAA | -70.76 | -209.07 | -5.95 | 34.33 | -69.82 | -209.07 | -5.01 | 30.25 |
| CGGCAAGCGC | -86.13 | -236.87 | -12.70 | 60.79 | -86.52 | -238.58 | -12.56 | 60.09 |
| UAGGUUAUAA | -73.74 | -216.62 | -6.59 | 37.12 | -73.29 | -214.88 | -6.67 | 37.47 |
| CGUACACAUGC | -81.90 | -227.58 | -11.35 | 56.24 | -82.45 | -227.28 | -11.99 | 58.85 |
| CCAUUGCUACC | -84.05 | -235.65 | -11.00 | 54.26 | -84.32 | -235.59 | -11.29 | 55.40 |
| CCAUCGCUACC | -84.09 | -233.28 | -11.78 | 57.49 | -84.59 | -233.46 | -12.21 | 59.19 |
| ACGUAUUAUGC | -82.58 | -235.94 | -9.44 | 48.19 | -82.49 | -235.35 | -9.53 | 48.55 |
| GCAUAAUACGU | -84.36 | -241.49 | -9.50 | 48.17 | -84.17 | -241.05 | -9.44 | 47.98 |
| CGCGAAUUCGCG | -96.22 | -265.28 | -13.99 | 62.90 | -96.57 | -267.17 | -13.75 | 61.91 |
| CUGACAAGUGUC | -89.54 | -249.47 | -12.20 | 57.83 | -89.95 | -249.58 | -12.58 | 59.23 |
| UUUUAUAAUAAA | -78.22 | -231.92 | -6.32 | 36.04 | -77.56 | -229.73 | -6.34 | 36.11 |
| AUUGGAUACAAA | -85.84 | -246.40 | -9.46 | 47.80 | -85.84 | -244.76 | -9.96 | 49.77 |
| CAACCAACCAAC | -94.63 | -264.35 | -12.68 | 58.42 | -95.00 | -264.38 | -13.04 | 59.67 |
| CUUCCUUCCUUC | -85.38 | -237.25 | -11.83 | 57.39 | -85.61 | -238.46 | -11.68 | 56.72 |
| GGAACAAGAUGC | -88.88 | -246.56 | -12.45 | 58.98 | -89.33 | -247.02 | -12.76 | 60.10 |
| GGAACUUGAUGC | -89.45 | -249.03 | -12.25 | 58.03 | -89.80 | -249.53 | -12.44 | 58.71 |
| AAUGGAUUACAA | -85.94 | -247.00 | -9.37 | 47.46 | -85.93 | -245.24 | -9.90 | 49.53 |
| GUCAGGAAUCUG | -89.73 | -251.11 | -11.89 | 56.54 | -90.11 | -250.86 | -12.34 | 58.23 |
| UUGUAAUCCAUU | -83.02 | -238.08 | -9.22 | 47.24 | -83.12 | -236.18 | -9.90 | 49.96 |

| | | | | | | | | |
|---|---|---|---|---|---|---|---|---|
| UUUGUAUCCAAU | -79.87 | -227.04 | -9.49 | 48.78 | -80.03 | -225.92 | -9.99 | 50.88 |
| CGCAUGGGUACGC | -107.13 | -294.80 | -15.74 | 66.02 | -107.93 | -295.26 | -16.40 | 68.05 |
| AGCCUAACUCAGC | -94.89 | -266.46 | -12.29 | 56.88 | -95.35 | -265.39 | -13.08 | 59.74 |
| CAUAUGGCCAUAUG | -100.99 | -281.11 | -13.85 | 61.07 | -101.62 | -280.56 | -14.64 | 63.75 |
| GUAUACCGGUAUAC | -104.90 | -295.39 | -13.33 | 58.36 | -105.48 | -293.45 | -14.51 | 62.25 |
| ACAUUAUUAUUACA | -101.91 | -306.40 | -6.92 | 38.06 | -100.79 | -302.41 | -7.04 | 38.44 |
| UACUAACAUUAACUA | -102.90 | -298.80 | -10.27 | 48.56 | -102.78 | -295.52 | -11.17 | 51.51 |
| AUACUUACUGAUUAG | -102.69 | -292.72 | -11.95 | 54.12 | -102.77 | -291.23 | -12.49 | 55.93 |
| GUACACUGUCUUAUA | -102.96 | -294.96 | -11.52 | 52.66 | -103.34 | -291.45 | -12.99 | 57.52 |
| GUAUGAGAGACUUUA | -107.41 | -303.94 | -13.19 | 57.34 | -107.66 | -302.70 | -13.82 | 59.39 |
| UUCUACCUAUGUGAU | -103.86 | -293.49 | -12.88 | 57.04 | -104.31 | -291.61 | -13.91 | 60.47 |
| AGUAGUAAUCACACC | -111.14 | -317.91 | -12.59 | 54.72 | -111.51 | -314.41 | -14.04 | 59.26 |
| AUCGUCUCGGUAUAA | -108.64 | -308.96 | -12.86 | 56.05 | -109.03 | -306.36 | -14.06 | 59.85 |
| ACGACAGGUUUACCA | -113.86 | -326.32 | -12.70 | 54.63 | -114.20 | -322.63 | -14.19 | 59.14 |
| CUUUCAUGUCCGCAU | -111.59 | -313.39 | -14.45 | 60.52 | -112.07 | -312.33 | -15.24 | 62.99 |
| UGGAUGUGUGAACAC | -113.62 | -318.48 | -14.90 | 61.50 | -114.60 | -315.48 | -16.80 | 67.39 |
| ACCCCGCAAUACAUG | -111.02 | -306.80 | -15.91 | 65.48 | -112.03 | -305.81 | -17.23 | 69.60 |
| GCAGUGGAUGUGAGA | -115.62 | -321.52 | -15.95 | 64.34 | -116.50 | -320.21 | -17.23 | 68.22 |
| GGUCCUUACUUGGUG | -115.58 | -320.75 | -16.15 | 65.00 | -116.37 | -320.19 | -17.11 | 67.86 |
| CGCCUCAUGCUCAUC | -111.55 | -308.54 | -15.90 | 65.29 | -112.39 | -308.16 | -16.86 | 68.23 |
| AAAUAGCCGGGCCGC | -126.31 | -350.58 | -17.63 | 66.72 | -127.08 | -350.29 | -18.49 | 69.06 |
| CCAGCCAGUCUCUCC | -114.66 | -315.59 | -16.83 | 67.44 | -115.66 | -315.35 | -17.90 | 70.67 |
| GACGACAAGACCGCG | -119.28 | -328.51 | -17.44 | 68.07 | -120.34 | -328.06 | -18.64 | 71.55 |
| CAGCCUCGUCGCAGC | -125.26 | -346.59 | -17.81 | 67.54 | -126.12 | -346.36 | -18.75 | 70.12 |
| CUCGCGGUCGAAGCG | -120.82 | -333.29 | -17.50 | 67.83 | -121.71 | -333.40 | -18.35 | 70.21 |
| GCGUCGGUCCGGGCU | -126.56 | -346.75 | -19.07 | 70.94 | -127.97 | -345.84 | -20.76 | 75.63 |
| CAACUUGAUAUUAUUA | -111.69 | -323.75 | -11.33 | 50.77 | -111.46 | -320.86 | -11.99 | 52.83 |
| CAUAUUGGCCAAUAUG | -115.52 | -323.15 | -15.34 | 62.46 | -116.10 | -322.26 | -16.20 | 65.01 |
| GUAUAACCGGUUAUAC | -117.88 | -335.79 | -13.79 | 57.20 | -118.34 | -332.48 | -15.27 | 61.58 |
| CGCGUACGCGUACGCG | -138.94 | -388.74 | -18.43 | 65.89 | -139.97 | -385.99 | -20.31 | 70.72 |
| CAACUUGAUAUUAAUA | -114.17 | -328.20 | -12.43 | 53.75 | -113.97 | -326.39 | -12.79 | 54.88 |
| UAUGUAUAUUUUGUAAUCAG | -138.90 | -402.53 | -14.11 | 54.75 | -138.96 | -397.21 | -15.83 | 59.08 |
| UUCAAGUUAAACAUUCUAUC | -133.68 | -385.47 | -14.19 | 55.69 | -134.10 | -379.78 | -16.37 | 61.40 |
| UGAUUCUACCUAUGUGAUUU | -138.92 | -398.15 | -15.49 | 58.23 | -139.70 | -391.80 | -18.24 | 65.23 |
| GAGAUUGUUUCCCUUUCAAA | -143.30 | -420.97 | -12.80 | 51.04 | -143.38 | -412.59 | -15.48 | 57.49 |
| AUGCAAUGCUACAUAUUCGC | -137.64 | -407.72 | -11.25 | 47.86 | -137.67 | -398.23 | -14.22 | 55.20 |
| CCACUAUACCAUCUAUGUAC | -150.75 | -434.47 | -16.06 | 57.78 | -151.34 | -427.49 | -18.82 | 64.25 |
| CCAUCAUUGUGUCUACCUCA | -149.25 | -425.65 | -17.30 | 60.97 | -150.11 | -419.78 | -19.98 | 67.36 |
| CGGGACCAACUAAAGGAAAU | -158.64 | -462.04 | -15.41 | 55.23 | -159.58 | -450.86 | -19.81 | 65.00 |
| UAGUGGCGAUUAGAUUCUGC | -153.21 | -439.01 | -17.11 | 59.86 | -153.81 | -433.18 | -19.52 | 65.46 |
| AGCUGCAGUGGAUGUGAGAA | -150.14 | -422.39 | -19.20 | 65.42 | -151.45 | -417.32 | -22.08 | 72.33 |
| UACUUCCAGUGCUCAGCGUA | -140.05 | -405.37 | -14.39 | 55.28 | -140.44 | -398.81 | -16.81 | 61.34 |
| CAGUGAGACAGCAAUGGUCG | -130.85 | -395.60 | -8.22 | 40.91 | -130.36 | -384.99 | -11.02 | 47.91 |
| CGAGCUUAUCCCUAUCCCUC | -144.34 | -427.80 | -11.72 | 48.43 | -144.27 | -418.17 | -14.64 | 55.32 |
| CGUACUAGCGUUGGUCAUGG | -154.72 | -432.63 | -20.61 | 67.89 | -156.15 | -428.40 | -23.34 | 74.26 |
| AAGGCGAGUCAGGCUCAGUG | -155.45 | -434.27 | -20.82 | 68.24 | -157.00 | -429.66 | -23.80 | 75.17 |
| ACCGACGACGCUGAUCCGAU | -156.99 | -441.47 | -20.13 | 66.27 | -158.50 | -435.64 | -23.45 | 73.90 |
| AGCAGUCCGCCACACCCUGA | -158.29 | -440.92 | -21.60 | 69.48 | -160.18 | -435.43 | -25.20 | 77.76 |
| CAGCCUCGUUCGCACAGCCC | -153.77 | -433.09 | -19.51 | 65.45 | -155.20 | -427.27 | -22.75 | 73.03 |
| GUGGUGGGCCGUGCGCUCUG | -168.29 | -478.21 | -20.04 | 63.92 | -170.14 | -468.41 | -24.93 | 74.44 |
| GUCCACGCCCGGUGCGACGG | -161.76 | -451.43 | -21.81 | 69.20 | -163.83 | -444.80 | -25.94 | 78.52 |
| GAUAUAGCAAAAUUCUAAGUUAAUA | -173.16 | -502.23 | -17.47 | 57.76 | -173.73 | -493.03 | -20.89 | 64.79 |
| AUAACUUUACGUGUGUGACCUAUUA | -170.15 | -500.98 | -14.85 | 52.79 | -171.13 | -486.41 | -20.35 | 64.08 |
| GUUCUAUACUCUUGAAGUUGAUUAC | -177.86 | -521.22 | -16.28 | 54.84 | -178.31 | -509.70 | -20.31 | 62.81 |
| CCCUGCACUUUAACUGAAUUGUUUA | -175.30 | -504.03 | -19.05 | 60.70 | -176.97 | -492.03 | -24.44 | 71.75 |
| UAACCAUACUGAAUACCUUUUGACG | -166.44 | -511.27 | -7.95 | 39.52 | -167.43 | -486.39 | -16.65 | 56.80 |
| UCCACACGGUAGUAAAAUUAGGCUU | -185.18 | -537.25 | -18.63 | 58.53 | -186.29 | -525.04 | -23.53 | 67.98 |
| UUCCAAAAGGAGUUAUGAGUUGCGA | -176.68 | -513.76 | -17.41 | 57.20 | -177.40 | -503.10 | -21.44 | 65.29 |

| | | | | | | | | |
|---|---|---|---|---|---|---|---|---|
| AAUAUCUCUCAUGCGCCAAGCUACA | -174.59 | -523.51 | -12.31 | 47.46 | -174.52 | -508.66 | -16.84 | 56.31 |
| UAGUAUAUCGCAGCAUCAUACAGGC | -190.19 | -546.15 | -20.89 | 62.16 | -191.60 | -535.20 | -25.69 | 71.30 |
| UGGAUUCUACUCAACCUUAGUCUGG | -185.35 | -530.70 | -20.84 | 62.78 | -186.69 | -521.00 | -25.18 | 71.26 |
| CGGAAUCCAUGUUACUUCGGCUAUC | -184.71 | -545.56 | -15.59 | 52.84 | -185.71 | -529.13 | -21.68 | 64.38 |
| CUGGUCUGGAUCUGAGAACUUCAGG | -183.43 | -543.62 | -14.91 | 51.69 | -184.48 | -526.12 | -21.38 | 64.00 |
| ACAGCGAAUGGACCUACGUGGCCUU | -191.69 | -547.98 | -21.81 | 63.71 | -193.63 | -535.79 | -27.54 | 74.58 |
| AGCAAGUCGAGCAGGGCCUACGUUU | -187.21 | -539.94 | -19.83 | 60.56 | -188.66 | -527.73 | -25.07 | 70.64 |
| GCGAGCGACAGGUUACUUGGCUGAU | -202.73 | -576.83 | -23.92 | 65.94 | -204.52 | -566.49 | -28.91 | 74.94 |
| AAAGGUGUCGCGGAGAGUCGUGCUG | -190.87 | -538.90 | -23.81 | 67.72 | -193.19 | -528.44 | -29.37 | 78.43 |
| AUGGGUGGGAGCCUCGGUAGCAGCC | -195.59 | -542.62 | -27.38 | 73.84 | -198.39 | -534.71 | -32.63 | 83.82 |
| CAGUGGGCUCCUGGGCGUGCUGGUC | -201.19 | -565.25 | -25.96 | 70.00 | -203.70 | -555.23 | -31.58 | 80.32 |
| GCCAACUCCGUCGCCGUUCGUGCGC | -204.39 | -603.01 | -17.46 | 54.37 | -204.80 | -588.17 | -22.47 | 63.02 |
| ACGGGUCCCCGCACCGCACCGCCAG | -204.15 | -571.56 | -26.97 | 71.34 | -207.19 | -560.20 | -33.52 | 83.29 |
| UUAUGUAUUAAGUUAUAUAGUAGUAGUAGU | -213.24 | -652.57 | -10.95 | 43.41 | -213.23 | -627.43 | -18.73 | 55.67 |
| AUUGAUAUCCUUUUCUAUUCAUCUUUCAUU | -202.24 | -599.37 | -16.44 | 52.83 | -203.09 | -581.41 | -22.85 | 63.94 |
| AAAGUACAUCAACAUAGAGAAUUGCAUUUC | -213.60 | -631.98 | -17.69 | 53.94 | -215.07 | -610.93 | -25.68 | 67.16 |
| CUUAAGAUAUGAGAACUUCAACUAAUGUGU | -209.09 | -622.20 | -16.20 | 51.89 | -209.72 | -603.42 | -22.66 | 62.68 |
| CUCAACUUGCGGUAAAUAAAUCGCUUAAUC | -216.16 | -625.43 | -22.28 | 61.21 | -217.74 | -610.67 | -28.43 | 71.53 |
| UAUUGAGAACAAGUGUCCGAUUAGCAGAAA | -217.41 | -650.08 | -15.89 | 50.80 | -218.12 | -628.74 | -23.21 | 62.53 |
| GUCAUACGACUGAGUGCAACAUUGUUCAAA | -217.74 | -673.54 | -8.95 | 40.33 | -218.75 | -639.74 | -20.44 | 57.90 |
| AACCUGCAACAUGGAGUUUUUGUCUCAUGC | -215.57 | -635.49 | -18.57 | 55.19 | -217.05 | -615.38 | -26.28 | 67.89 |
| CCGUGCGGUGUGUACGUUUUAUUCAUCAUA | -231.32 | -669.39 | -23.81 | 61.88 | -233.66 | -650.60 | -31.97 | 74.74 |
| GUUCACGUCCGAAAGCUCGAAAAAGGAUAC | -221.87 | -661.12 | -16.92 | 52.08 | -223.15 | -637.97 | -25.38 | 65.46 |
| AGUCUGGUCUGGAUCUGAGAACUUCAGGCU | -215.48 | -643.11 | -16.11 | 51.28 | -216.81 | -619.76 | -24.69 | 65.19 |
| UCGGAGAAAUCACUGAGCUGCCUGAGAAGA | -219.35 | -659.04 | -15.05 | 49.38 | -220.40 | -634.48 | -23.71 | 63.05 |
| CUUCAACGGAUCAGGUAGGACUGUGGUGGG | -220.25 | -670.97 | -12.25 | 45.12 | -222.18 | -637.80 | -24.46 | 64.07 |
| ACGCCCACAGGAUUAGGCUGGCCCACAUUG | -224.37 | -656.98 | -20.70 | 57.76 | -227.10 | -632.89 | -30.91 | 74.13 |
| GUUAUUCCGCAGUCCGAUGGCAGCAGGCUC | -220.92 | -642.85 | -21.64 | 59.61 | -223.05 | -623.66 | -29.71 | 72.81 |
| UCAGUAGGCGUGACGCAGAGCUGGCGAUGG | -228.99 | -668.80 | -21.66 | 58.78 | -231.28 | -647.25 | -30.64 | 72.93 |
| CGCGCCACGUGUGAUCUACAGCCGUUCGGC | -238.04 | -687.27 | -24.99 | 62.91 | -240.40 | -668.91 | -33.03 | 75.27 |
| GACCUGACGUGGACCGCUCCUGGGCGUGGU | -226.79 | -675.32 | -17.44 | 52.52 | -229.11 | -647.37 | -28.42 | 69.61 |
| GCCCCUCCACUGGCCGACGGCAGCAGGCUC | -229.88 | -658.80 | -25.65 | 64.98 | -232.95 | -639.98 | -34.55 | 79.25 |
| CGCCGCUGCCGACUGGAGGAGCGCGGGACG | -252.89 | -730.86 | -26.33 | 63.18 | -254.80 | -713.62 | -33.58 | 73.67 |

[a] All duplexes consist of the denoted RNA strand and its complementary DNA strand.

**Table S3.** Corrected Thermodynamic Parameters for RNA/PSDNA Duplexes

| RNA Sequence[a] | MMPBSA | | | | MMGBSA | | | |
|---|---|---|---|---|---|---|---|---|
| | $\Delta H°$ (kcal/mol) | $\Delta S°$ (cal/mol/K) | $\Delta G°_{37}$ (kcal/mol) | $T_m$ (°C) | $\Delta H°$ (kcal/mol) | $\Delta S°$ (cal/mol/K) | $\Delta G°_{37}$ (kcal/mol) | $T_m$ (°C) |
| CCGG | -32.26 | -93.45 | -3.29 | 8.60 | -32.51 | -91.76 | -4.06 | 14.99 |
| CGCG | -33.25 | -96.27 | -3.40 | 10.24 | -33.35 | -95.20 | -3.83 | 13.67 |
| GCGC | -35.02 | -103.63 | -2.90 | 7.72 | -35.06 | -101.67 | -3.54 | 12.51 |
| AGCCG | -40.79 | -118.45 | -4.06 | 19.20 | -40.87 | -116.96 | -4.61 | 22.98 |
| CACAG | -40.54 | -119.83 | -3.39 | 14.57 | -40.40 | -118.32 | -3.72 | 16.71 |
| CGUGC | -35.37 | -106.74 | -2.28 | 3.64 | -35.48 | -103.43 | -3.41 | 11.84 |
| GCACG | -42.74 | -124.32 | -4.20 | 20.81 | -42.76 | -122.94 | -4.65 | 23.79 |
| CGGCU | -43.12 | -125.61 | -4.18 | 20.84 | -43.29 | -123.46 | -5.02 | 26.42 |
| GGUGG | -40.20 | -115.39 | -4.43 | 21.48 | -40.53 | -113.53 | -5.34 | 28.00 |
| CCGCGG | -50.88 | -139.86 | -7.52 | 43.04 | -51.36 | -140.09 | -7.93 | 45.55 |
| CGAUCG | -52.23 | -151.39 | -5.30 | 29.73 | -52.05 | -150.73 | -5.32 | 29.84 |
| CGCGCG | -51.57 | -143.15 | -7.20 | 40.92 | -51.88 | -143.49 | -7.40 | 42.14 |
| CGGCCG | -52.88 | -147.39 | -7.19 | 40.81 | -53.33 | -146.83 | -7.81 | 44.51 |
| CGUACG | -47.07 | -134.70 | -5.32 | 29.09 | -47.23 | -133.63 | -5.80 | 32.17 |
| GACGUC | -45.32 | -133.74 | -3.86 | 19.60 | -45.32 | -131.38 | -4.59 | 24.15 |
| GCAUGC | -49.25 | -141.54 | -5.38 | 29.78 | -49.34 | -140.43 | -5.80 | 32.37 |
| GCCGGC | -51.56 | -142.67 | -7.33 | 41.73 | -52.10 | -142.13 | -8.04 | 46.14 |
| GCGAGC | -50.90 | -144.32 | -6.16 | 34.60 | -51.11 | -143.56 | -6.61 | 37.33 |
| GCGCGC | -53.86 | -151.18 | -6.99 | 39.55 | -54.19 | -150.59 | -7.51 | 42.55 |
| GCUAGC | -48.08 | -137.45 | -5.47 | 30.21 | -48.19 | -136.65 | -5.82 | 32.40 |
| GGAUCC | -49.36 | -143.98 | -4.73 | 25.94 | -49.40 | -142.02 | -5.37 | 29.78 |
| GGCGCC | -50.73 | -144.63 | -5.89 | 33.03 | -51.07 | -142.89 | -6.78 | 38.37 |
| GGGACC | -50.97 | -147.03 | -5.39 | 30.07 | -51.24 | -144.74 | -6.37 | 35.92 |
| GUGAAC | -46.58 | -136.05 | -4.41 | 23.34 | -46.62 | -134.15 | -5.03 | 27.19 |
| UCAUGA | -43.72 | -130.15 | -3.37 | 15.99 | -43.61 | -127.82 | -3.99 | 19.80 |
| UGAUCA | -40.27 | -124.69 | -1.61 | 3.14 | -40.01 | -120.95 | -2.51 | 8.57 |
| CCCGGG | -50.93 | -137.33 | -8.36 | 48.44 | -51.61 | -138.01 | -8.83 | 51.30 |
| ACCGCA | -47.07 | -134.62 | -5.34 | 29.20 | -47.22 | -133.59 | -5.81 | 32.22 |
| AGUUGC | -45.01 | -130.60 | -4.52 | 23.63 | -45.11 | -128.90 | -5.15 | 27.67 |
| AUGCGC | -45.79 | -131.49 | -5.02 | 27.00 | -45.95 | -130.17 | -5.60 | 30.70 |
| CCAACG | -51.69 | -149.72 | -5.28 | 29.56 | -51.58 | -148.83 | -5.45 | 30.48 |
| CGGACG | -46.93 | -130.48 | -6.48 | 36.56 | -47.39 | -129.96 | -7.10 | 40.65 |
| CGGUGC | -51.55 | -145.50 | -6.44 | 36.35 | -51.88 | -144.53 | -7.07 | 40.16 |
| CGUGCC | -50.90 | -144.67 | -6.05 | 33.99 | -51.15 | -143.56 | -6.65 | 37.58 |
| CGUUGA | -44.03 | -126.08 | -4.94 | 26.08 | -44.20 | -124.96 | -5.46 | 29.57 |
| CGUUGC | -47.99 | -136.07 | -5.81 | 32.26 | -48.18 | -135.42 | -6.20 | 34.79 |
| CGUUGU | -43.28 | -127.96 | -3.61 | 17.27 | -43.25 | -125.72 | -4.28 | 21.54 |
| GCCUGC | -48.55 | -136.05 | -6.38 | 35.89 | -48.88 | -135.56 | -6.86 | 38.98 |
| UGCGCA | -43.08 | -122.92 | -4.97 | 26.05 | -43.40 | -121.36 | -5.78 | 31.60 |
| UGUUGC | -45.40 | -131.89 | -4.52 | 23.71 | -45.48 | -130.22 | -5.11 | 27.50 |
| CAAUCG | -48.28 | -146.53 | -2.86 | 14.96 | -47.91 | -143.63 | -3.39 | 17.79 |
| CACGGC | -49.43 | -137.44 | -6.82 | 38.70 | -49.89 | -136.92 | -7.44 | 42.64 |
| CGAUUG | -47.23 | -136.70 | -4.85 | 26.23 | -47.13 | -136.01 | -4.96 | 26.89 |
| GCACCG | -52.91 | -151.61 | -5.91 | 33.28 | -52.99 | -150.65 | -6.28 | 35.43 |
| GGCACG | -51.90 | -148.00 | -6.02 | 33.85 | -52.10 | -146.83 | -6.59 | 37.20 |
| CCCAGGG | -58.27 | -157.65 | -9.39 | 52.89 | -59.04 | -158.04 | -10.05 | 56.49 |
| CAAAAAG | -51.27 | -147.97 | -5.40 | 30.18 | -51.04 | -147.86 | -5.21 | 29.05 |
| AAUACCG | -50.37 | -148.71 | -4.27 | 23.57 | -50.36 | -146.08 | -5.07 | 28.14 |
| AGCCGUG | -55.63 | -160.40 | -5.91 | 33.43 | -55.84 | -158.28 | -6.78 | 38.23 |
| AGCUUCA | -51.65 | -152.46 | -4.39 | 24.53 | -51.59 | -149.92 | -5.12 | 28.60 |
| GGACUUA | -46.04 | -132.95 | -4.83 | 25.81 | -46.30 | -130.83 | -5.74 | 31.69 |
| ACGUAUG | -56.37 | -166.27 | -4.82 | 27.75 | -56.18 | -164.05 | -5.32 | 30.33 |
| CACGGCU | -57.78 | -161.93 | -7.58 | 42.63 | -58.23 | -160.99 | -8.32 | 46.70 |

| | | | | | | | | |
|---|---|---|---|---|---|---|---|---|
| CAUACGU | -54.24 | -156.44 | -5.74 | 32.43 | -54.24 | -155.26 | -6.11 | 34.51 |
| UAAGUCC | -51.62 | -150.52 | -4.95 | 27.68 | -51.62 | -148.61 | -5.55 | 31.11 |
| UGAAGCU | -50.25 | -146.30 | -4.90 | 27.13 | -50.30 | -144.37 | -5.55 | 30.95 |
| AAAAAAAA | -51.06 | -154.22 | -3.25 | 18.16 | -50.83 | -150.75 | -4.10 | 22.73 |
| UAGAUCUA | -56.28 | -166.53 | -4.65 | 26.85 | -56.15 | -163.76 | -5.39 | 30.70 |
| UCUAUAGA | -57.39 | -166.70 | -5.72 | 32.54 | -57.52 | -164.28 | -6.59 | 37.21 |
| CAAAAAAG | -59.26 | -170.96 | -6.26 | 35.46 | -58.98 | -170.83 | -6.03 | 34.24 |
| CAAGCUUG | -62.55 | -175.70 | -8.08 | 44.76 | -62.67 | -176.04 | -8.10 | 44.81 |
| CAUCGAUG | -58.02 | -163.23 | -7.41 | 41.66 | -58.35 | -162.47 | -7.99 | 44.81 |
| CGAUAUCG | -63.39 | -181.87 | -7.01 | 39.23 | -63.42 | -180.74 | -7.39 | 41.10 |
| CGUCGACG | -66.12 | -188.87 | -7.57 | 41.82 | -66.36 | -187.12 | -8.35 | 45.62 |
| GAAGCUUC | -60.45 | -172.19 | -7.07 | 39.68 | -60.55 | -171.45 | -7.40 | 41.40 |
| GAUCGAUC | -59.74 | -173.71 | -5.89 | 33.57 | -59.80 | -171.37 | -6.68 | 37.64 |
| GAUGCAUC | -62.89 | -184.01 | -5.85 | 33.55 | -62.87 | -181.39 | -6.64 | 37.39 |
| GGAAUUCC | -61.92 | -180.41 | -6.00 | 34.21 | -61.88 | -178.29 | -6.61 | 37.24 |
| GGACGUCC | -67.29 | -191.05 | -8.06 | 44.09 | -67.71 | -189.04 | -9.11 | 49.15 |
| GGAGCUCC | -62.95 | -174.99 | -8.71 | 47.96 | -63.50 | -174.33 | -9.46 | 51.85 |
| GGUAUACC | -67.36 | -196.42 | -6.47 | 36.61 | -67.41 | -193.55 | -7.41 | 40.95 |
| GUACGUAC | -64.15 | -185.79 | -6.56 | 36.99 | -64.36 | -182.97 | -7.64 | 42.32 |
| GUAGCUAC | -65.10 | -189.17 | -6.46 | 36.54 | -65.04 | -187.22 | -7.00 | 39.14 |
| GUUGCAAC | -58.67 | -167.22 | -6.83 | 38.46 | -59.05 | -165.17 | -7.85 | 43.95 |
| AAGCGUAG | -60.32 | -172.07 | -6.97 | 39.16 | -60.54 | -170.64 | -7.64 | 42.67 |
| AAUCCAGU | -55.44 | -160.04 | -5.83 | 32.98 | -55.59 | -158.08 | -6.59 | 37.20 |
| ACAUAUGU | -57.79 | -168.94 | -5.42 | 31.03 | -57.74 | -166.78 | -6.04 | 34.23 |
| ACCUAGUC | -68.88 | -202.64 | -6.06 | 34.78 | -68.59 | -200.32 | -6.49 | 36.68 |
| ACGACCUC | -64.59 | -183.93 | -7.57 | 41.95 | -64.88 | -182.28 | -8.37 | 45.94 |
| AGAGAGAG | -55.95 | -154.73 | -7.98 | 45.13 | -56.46 | -154.51 | -8.56 | 48.43 |
| AGAGCUCU | -58.92 | -167.75 | -6.92 | 38.93 | -59.12 | -166.60 | -7.48 | 41.92 |
| AGCGUAAG | -61.47 | -176.48 | -6.77 | 38.06 | -61.63 | -174.78 | -7.45 | 41.56 |
| AGUCCUGA | -60.32 | -173.21 | -6.62 | 37.35 | -60.56 | -171.12 | -7.52 | 42.01 |
| AUGCGCAU | -60.07 | -171.87 | -6.79 | 38.20 | -60.34 | -169.96 | -7.65 | 42.75 |
| CACGGCUC | -68.33 | -191.62 | -8.93 | 48.13 | -68.76 | -190.75 | -9.63 | 51.48 |
| CCAUAUGG | -59.69 | -166.83 | -7.97 | 44.55 | -60.10 | -166.28 | -8.55 | 47.66 |
| CGCGUAUA | -63.57 | -183.59 | -6.65 | 37.46 | -63.62 | -181.73 | -7.29 | 40.59 |
| CGCUGUAA | -62.67 | -177.98 | -7.50 | 41.72 | -62.89 | -176.95 | -8.03 | 44.44 |
| CGUCGUCC | -63.81 | -181.53 | -7.54 | 41.84 | -64.15 | -179.77 | -8.43 | 46.30 |
| CUAGUGGA | -59.64 | -168.32 | -7.46 | 41.75 | -59.98 | -167.26 | -8.13 | 45.37 |
| CUCACGGC | -66.56 | -187.04 | -8.58 | 46.70 | -66.93 | -186.26 | -9.19 | 49.68 |
| CUGAGUCC | -63.44 | -179.56 | -7.77 | 43.06 | -63.73 | -178.48 | -8.40 | 46.23 |
| GAAUAUUC | -54.58 | -162.75 | -4.12 | 23.78 | -54.34 | -159.97 | -4.76 | 27.06 |
| GACUAGUC | -61.17 | -175.80 | -6.67 | 37.58 | -61.33 | -173.97 | -7.40 | 41.35 |
| GAGUACUC | -62.12 | -179.26 | -6.55 | 36.96 | -62.27 | -177.08 | -7.38 | 41.15 |
| GAUUAAUC | -58.34 | -173.31 | -4.61 | 26.98 | -58.08 | -170.66 | -5.18 | 29.80 |
| GCAUAUGC | -60.64 | -173.35 | -6.90 | 38.78 | -60.88 | -171.65 | -7.67 | 42.79 |
| GCCAGUUA | -58.86 | -169.70 | -6.25 | 35.41 | -59.08 | -167.43 | -7.18 | 40.30 |
| GGACCUCG | -61.47 | -170.28 | -8.68 | 48.10 | -62.08 | -169.59 | -9.51 | 52.49 |
| GGAGCACG | -64.10 | -177.75 | -8.99 | 49.25 | -64.69 | -177.12 | -9.78 | 53.29 |
| GGUGCCAA | -63.02 | -178.18 | -7.79 | 43.17 | -63.60 | -175.90 | -9.07 | 49.78 |
| GUCGAACA | -67.18 | -195.67 | -6.52 | 36.83 | -67.17 | -193.16 | -7.29 | 40.42 |
| GUCUAGAC | -59.86 | -171.20 | -6.78 | 38.18 | -60.16 | -169.18 | -7.72 | 43.11 |
| UAGGCCUA | -64.15 | -184.44 | -6.97 | 39.01 | -64.37 | -182.25 | -7.87 | 43.46 |
| UAUGCAUA | -60.91 | -180.06 | -5.10 | 29.73 | -60.70 | -177.45 | -5.69 | 32.63 |
| UGAGCUCA | -55.41 | -157.71 | -6.52 | 36.80 | -55.73 | -156.11 | -7.34 | 41.43 |
| AAUGUCGC | -62.52 | -180.21 | -6.66 | 37.50 | -62.64 | -178.31 | -7.36 | 41.04 |
| ACUGGAUU | -62.16 | -178.93 | -6.69 | 37.65 | -62.35 | -176.80 | -7.55 | 42.00 |
| CAACAGCA | -62.33 | -176.48 | -7.62 | 42.38 | -62.57 | -175.60 | -8.13 | 45.01 |
| CUACGCUU | -62.97 | -181.33 | -6.76 | 37.98 | -63.01 | -179.82 | -7.26 | 40.52 |
| CUUACGCU | -62.10 | -177.63 | -7.03 | 39.39 | -62.20 | -176.45 | -7.50 | 41.80 |

| | | | | | | | | |
|---|---|---|---|---|---|---|---|---|
| GACUAGGU | -62.11 | -176.27 | -7.47 | 41.61 | -62.47 | -174.67 | -8.32 | 45.99 |
| GAGCCGUG | -63.38 | -177.03 | -8.50 | 46.82 | -63.93 | -175.95 | -9.39 | 51.38 |
| GAGGUCGU | -60.54 | -167.31 | -8.68 | 48.26 | -61.18 | -166.71 | -9.50 | 52.68 |
| GCCGUGAG | -67.29 | -187.77 | -9.08 | 49.09 | -67.84 | -186.83 | -9.92 | 53.17 |
| GCGACAUU | -65.43 | -187.41 | -7.34 | 40.73 | -65.64 | -185.60 | -8.10 | 44.46 |
| UAACUGGC | -59.74 | -168.64 | -7.46 | 41.78 | -60.12 | -167.38 | -8.24 | 45.92 |
| UCCACUAG | -56.91 | -162.60 | -6.50 | 36.73 | -57.19 | -160.87 | -7.32 | 41.19 |
| UGCUGUUG | -57.02 | -160.93 | -7.13 | 40.16 | -57.52 | -159.18 | -8.17 | 45.99 |
| UGUUCGAC | -58.11 | -167.00 | -6.33 | 35.82 | -58.42 | -164.58 | -7.41 | 41.58 |
| UUACAGCG | -63.17 | -181.98 | -6.75 | 37.97 | -63.23 | -180.36 | -7.32 | 40.78 |
| UUGGCACC | -64.07 | -182.96 | -7.35 | 40.88 | -64.45 | -180.67 | -8.44 | 46.35 |
| CAAAAAAAG | -62.48 | -177.90 | -7.33 | 40.89 | -62.35 | -178.19 | -7.11 | 39.77 |
| CAAACAAAG | -70.25 | -202.78 | -7.39 | 40.71 | -70.09 | -201.73 | -7.55 | 41.45 |
| CAAAUAAAG | -68.65 | -200.21 | -6.59 | 37.12 | -68.28 | -199.12 | -6.55 | 36.98 |
| CAAAGAAAG | -65.75 | -184.94 | -8.42 | 46.02 | -65.81 | -185.43 | -8.33 | 45.56 |
| AAAAAAAAA | -59.94 | -176.74 | -5.15 | 29.89 | -59.49 | -175.48 | -5.09 | 29.55 |
| AUAACUGGC | -67.34 | -190.78 | -8.20 | 44.73 | -67.69 | -189.32 | -9.00 | 48.60 |
| AUCUAUCCG | -66.02 | -189.92 | -7.15 | 39.78 | -66.12 | -188.15 | -7.79 | 42.90 |
| CGCUGUUAC | -70.64 | -198.60 | -9.08 | 48.47 | -71.04 | -197.66 | -9.76 | 51.65 |
| GCCAGUUAA | -64.21 | -182.69 | -7.58 | 42.02 | -64.53 | -181.02 | -8.41 | 46.17 |
| CAACAGCAA | -72.58 | -209.08 | -7.77 | 42.23 | -72.73 | -206.74 | -8.64 | 46.13 |
| CAACAGCAU | -70.95 | -202.95 | -8.03 | 43.57 | -71.21 | -200.86 | -8.94 | 47.73 |
| CGCUGUUAG | -69.25 | -192.62 | -9.54 | 50.97 | -69.68 | -192.54 | -10.00 | 53.10 |
| CUAACAGCG | -72.32 | -203.48 | -9.25 | 48.95 | -72.55 | -203.21 | -9.56 | 50.37 |
| GUAACAGCG | -71.92 | -203.15 | -8.95 | 47.64 | -72.26 | -202.01 | -9.63 | 50.78 |
| UUAACUGGC | -67.15 | -190.50 | -8.09 | 44.26 | -67.52 | -188.82 | -8.99 | 48.56 |
| AUGAGCUCAU | -71.12 | -204.71 | -7.66 | 41.87 | -71.38 | -202.00 | -8.76 | 46.87 |
| GCGAAUUCGC | -80.44 | -226.70 | -10.17 | 51.54 | -80.86 | -225.38 | -11.00 | 54.98 |
| GCGAAAAGCG | -79.43 | -221.07 | -10.90 | 54.92 | -79.87 | -221.06 | -11.34 | 56.73 |
| AUCAAUCAUA | -76.08 | -224.44 | -6.50 | 36.75 | -75.71 | -221.67 | -7.00 | 38.78 |
| UUGUAGUCAU | -69.83 | -204.16 | -6.54 | 36.91 | -70.01 | -200.31 | -7.91 | 43.11 |
| GAAAUGAAAG | -73.57 | -210.46 | -8.33 | 44.63 | -73.74 | -208.74 | -9.03 | 47.74 |
| CCAACUUCUU | -74.58 | -213.79 | -8.30 | 44.41 | -74.67 | -212.17 | -8.89 | 47.00 |
| AUCGUCUGGA | -73.29 | -206.55 | -9.26 | 48.86 | -74.03 | -203.84 | -10.84 | 56.02 |
| AGCGUAAGUC | -75.07 | -212.07 | -9.33 | 48.86 | -75.61 | -209.98 | -10.52 | 54.11 |
| CGAUCUGCGA | -73.91 | -206.10 | -10.01 | 52.20 | -74.51 | -205.05 | -10.94 | 56.39 |
| UGGCGAGCAC | -76.22 | -209.79 | -11.18 | 57.01 | -77.24 | -208.27 | -12.67 | 63.64 |
| GAUGCGCUCG | -79.44 | -221.05 | -10.92 | 54.98 | -80.12 | -219.99 | -11.92 | 59.24 |
| GGGACCGCCU | -80.57 | -218.78 | -12.74 | 62.77 | -81.79 | -217.95 | -14.23 | 69.07 |
| AAAAAAAAAA | -69.12 | -204.27 | -5.80 | 33.61 | -68.59 | -202.53 | -5.81 | 33.64 |
| CGGCAAGCGC | -84.94 | -234.36 | -12.29 | 59.41 | -85.74 | -233.85 | -13.24 | 63.19 |
| UAGGUUAUAA | -73.27 | -216.35 | -6.20 | 35.47 | -73.11 | -212.74 | -7.16 | 39.57 |
| CGUACACAUGC | -83.78 | -234.16 | -11.19 | 55.13 | -84.58 | -232.13 | -12.62 | 60.91 |
| CCAUUGCUACC | -85.21 | -239.45 | -10.98 | 53.93 | -85.76 | -237.89 | -12.01 | 58.03 |
| CCAUCGCUACC | -85.53 | -239.37 | -11.33 | 55.28 | -86.23 | -237.63 | -12.57 | 60.20 |
| ACGUAUUAUGC | -81.28 | -234.89 | -8.46 | 44.42 | -81.54 | -231.38 | -9.82 | 49.88 |
| GCAUAAUACGU | -85.04 | -243.97 | -9.41 | 47.73 | -85.19 | -241.74 | -10.25 | 51.01 |
| CGCGAAUUCGCG | -91.21 | -249.64 | -13.82 | 63.81 | -92.20 | -249.45 | -14.87 | 67.68 |
| CUGACAAGUGUC | -95.23 | -268.08 | -12.12 | 56.21 | -95.81 | -266.14 | -13.31 | 60.46 |
| UUUUAUAAUAAA | -84.64 | -253.20 | -6.14 | 35.46 | -84.01 | -249.31 | -6.72 | 37.56 |
| AUUGGAUACAAA | -86.29 | -249.40 | -8.98 | 45.92 | -86.62 | -245.41 | -10.55 | 51.93 |
| CAACCAACCAAC | -91.07 | -254.23 | -12.26 | 57.69 | -91.80 | -252.73 | -13.45 | 62.15 |
| CUUCCUUCCUUC | -93.03 | -261.27 | -12.04 | 56.37 | -93.21 | -261.47 | -12.15 | 56.75 |
| GGAACAAGAUGC | -84.06 | -231.31 | -12.35 | 59.94 | -85.17 | -229.68 | -13.97 | 66.54 |
| GGAACUUGAUGC | -92.91 | -260.16 | -12.26 | 57.24 | -93.58 | -258.54 | -13.43 | 61.55 |
| AAUGGAUUACAA | -86.37 | -248.59 | -9.31 | 47.17 | -86.69 | -245.14 | -10.70 | 52.53 |
| GUCAGGAAUCUG | -88.53 | -247.77 | -11.72 | 56.17 | -89.26 | -245.92 | -13.02 | 61.18 |
| UUGUAAUCCAUU | -88.27 | -253.49 | -9.69 | 48.36 | -88.49 | -250.72 | -10.77 | 52.45 |

| | | | | | | | | |
|---|---|---|---|---|---|---|---|---|
| UUUGUAUCCAAU | -84.31 | -242.62 | -9.10 | 46.62 | -84.60 | -239.44 | -10.37 | 51.60 |
| CGCAUGGGUACGC | -103.01 | -281.87 | -15.63 | 66.91 | -104.49 | -279.95 | -17.71 | 73.99 |
| AGCCUAACUCAGC | -101.61 | -284.45 | -13.43 | 59.45 | -102.32 | -282.74 | -14.67 | 63.67 |
| CAUAUGGCCAUAUG | -103.11 | -295.01 | -11.66 | 53.08 | -103.79 | -290.30 | -13.80 | 60.21 |
| GUAUACCGGUAUAC | -109.71 | -312.28 | -12.90 | 55.98 | -110.47 | -307.87 | -15.03 | 62.70 |
| ACAUUAUUAUUACA | -101.10 | -295.59 | -9.47 | 46.15 | -101.04 | -291.12 | -10.79 | 50.52 |
| UACUAACAUUAACUA | -103.68 | -304.30 | -9.34 | 45.51 | -103.73 | -298.58 | -11.17 | 51.38 |
| AUACUUACUGAUUAG | -112.70 | -340.77 | -7.06 | 38.34 | -112.08 | -332.61 | -8.97 | 43.76 |
| GUACACUGUCUUAUA | -111.73 | -321.50 | -12.06 | 53.01 | -112.27 | -316.32 | -14.21 | 59.62 |
| GUAUGAGAGACUUUA | -104.98 | -296.20 | -13.16 | 57.75 | -105.86 | -292.61 | -15.15 | 64.35 |
| UUCUACCUAUGUGAU | -107.94 | -306.73 | -12.86 | 56.16 | -108.63 | -302.94 | -14.71 | 62.12 |
| AGUAGUAAUCACACC | -113.49 | -323.80 | -13.12 | 55.96 | -114.28 | -318.84 | -15.44 | 63.08 |
| AUCGUCUCGGUAUAA | -112.31 | -321.63 | -12.60 | 54.58 | -113.00 | -316.51 | -14.88 | 61.60 |
| ACGACAGGUUUACCA | -118.77 | -339.02 | -13.67 | 56.70 | -119.66 | -333.39 | -16.31 | 64.45 |
| CUUUCAUGUCCGCAU | -108.78 | -317.67 | -10.30 | 47.99 | -109.22 | -310.72 | -12.90 | 56.05 |
| UGGAUGUGUGAACAC | -111.59 | -316.19 | -13.57 | 57.73 | -112.78 | -310.53 | -16.52 | 66.98 |
| ACCCCGCAAUACAUG | -113.03 | -319.45 | -14.00 | 58.79 | -114.15 | -314.47 | -16.67 | 67.08 |
| GCAGUGGAUGUGAGA | -109.49 | -302.06 | -15.85 | 65.70 | -111.20 | -298.16 | -18.77 | 75.19 |
| GGUCCUUACUUGGUG | -111.02 | -314.07 | -13.66 | 58.14 | -112.36 | -308.01 | -16.88 | 68.31 |
| CGCCUCAUGCUCAUC | -113.18 | -314.39 | -15.72 | 64.25 | -114.35 | -311.94 | -17.65 | 70.24 |
| AAAUAGCCGGGCCGC | -122.40 | -339.14 | -17.27 | 66.67 | -123.88 | -335.89 | -19.76 | 73.92 |
| CCAGCCAGUCUCUCC | -116.85 | -322.36 | -16.92 | 67.10 | -118.19 | -320.29 | -18.91 | 73.11 |
| GACGACAAGACCGCG | -119.28 | -331.18 | -16.61 | 65.47 | -120.61 | -328.17 | -18.88 | 72.21 |
| CAGCCUCGUCGCAGC | -125.96 | -348.41 | -17.95 | 67.77 | -127.33 | -345.99 | -20.08 | 73.76 |
| CUCGCGGUCGAAGCG | -121.47 | -336.59 | -17.13 | 66.50 | -122.79 | -334.04 | -19.24 | 72.65 |
| GCGUCGGUCCGGGCU | -126.66 | -346.13 | -19.36 | 71.80 | -128.64 | -343.14 | -22.26 | 80.05 |
| CAACUUGAUAUUAUUA | -113.08 | -327.63 | -11.51 | 51.15 | -113.31 | -322.68 | -13.28 | 56.49 |
| CAUAUUGGCCAAUAUG | -121.34 | -347.43 | -13.63 | 56.14 | -122.02 | -342.18 | -15.94 | 62.77 |
| GUAUAACCGGUUAUAC | -117.61 | -338.21 | -12.76 | 54.21 | -118.25 | -332.49 | -15.18 | 61.33 |
| CGCGUACGCGUACGCG | -130.05 | -359.79 | -18.51 | 68.32 | -131.76 | -355.95 | -21.41 | 76.34 |
| CAACUUGAUAUUAAUA | -106.82 | -307.44 | -11.51 | 52.02 | -107.38 | -302.25 | -13.69 | 58.99 |
| UAUGUAUAUUUUGUAAUCAG | -141.27 | -413.28 | -13.15 | 52.10 | -141.77 | -403.98 | -16.54 | 60.41 |
| UUCAAGUUAAACAUUCUAUC | -141.67 | -414.63 | -13.13 | 52.02 | -142.39 | -404.25 | -17.07 | 61.65 |
| UGAUUCUACCUAUGUGAUUU | -148.52 | -426.91 | -16.18 | 58.39 | -149.60 | -418.41 | -19.89 | 67.26 |
| GAGAUUGUUUCCCUUUCAAA | -142.74 | -414.39 | -14.28 | 54.65 | -143.45 | -405.67 | -17.69 | 63.02 |
| AUGCAAUGCUACAUAUUCGC | -141.58 | -420.63 | -11.18 | 47.39 | -142.36 | -406.87 | -16.23 | 59.53 |
| CCACUAUACCAUCUAUGUAC | -145.37 | -417.65 | -15.89 | 58.20 | -146.56 | -408.82 | -19.83 | 67.79 |
| CCAUCAUUGUGUCUACCUCA | -90.10 | -271.05 | -6.07 | 35.29 | -91.60 | -256.42 | -12.11 | 56.98 |
| CGGGACCAACUAAAGGAAAU | -138.05 | -396.93 | -15.01 | 57.13 | -139.54 | -386.87 | -19.61 | 68.92 |
| UAGUGGCGAUUAGAUUCUGC | -156.61 | -447.86 | -17.77 | 60.83 | -157.99 | -438.93 | -21.92 | 70.30 |
| AGCUGCAGUGGAUGUGAGAA | -135.93 | -385.54 | -16.41 | 61.16 | -138.08 | -375.20 | -21.76 | 75.30 |
| UACUUCCAGUGCUCAGCGUA | -145.67 | -415.64 | -16.82 | 60.42 | -146.89 | -408.08 | -20.39 | 69.15 |
| CAGUGAGACAGCAAUGGUCG | -152.16 | -435.13 | -17.27 | 60.40 | -153.60 | -426.01 | -21.54 | 70.43 |
| CGAGCUUAUCCCUAUCCCUC | -157.37 | -448.53 | -18.33 | 61.98 | -158.82 | -439.99 | -22.43 | 71.33 |
| CGUACUAGCGUUGGUCAUGG | -155.05 | -432.89 | -20.86 | 68.42 | -157.21 | -425.72 | -25.23 | 78.72 |
| AAGGCGAGUCAGGCUCAGUG | -156.58 | -444.55 | -18.77 | 63.14 | -158.59 | -434.42 | -23.92 | 75.03 |
| ACCGACGACGCUGAUCCGAU | -161.93 | -460.35 | -19.22 | 63.21 | -163.64 | -451.13 | -23.79 | 73.41 |
| AGCAGUCCGCCACACCCUGA | -158.02 | -443.35 | -20.59 | 67.12 | -160.23 | -434.92 | -25.40 | 78.25 |
| CAGCCUCGUUCGCACAGCCC | -157.64 | -454.81 | -16.65 | 58.12 | -158.75 | -445.06 | -20.78 | 67.43 |
| GUGGUGGGCCGUGCGCUCUG | -164.01 | -456.79 | -22.41 | 70.08 | -166.62 | -448.25 | -27.66 | 81.88 |
| GUCCACGCCCGGUGCGACGG | -175.90 | -488.24 | -24.55 | 72.23 | -178.52 | -480.64 | -29.52 | 82.68 |
| GAUAUAGCAAAAUUCUAAGUUAAUA | -170.77 | -525.47 | -7.88 | 39.32 | -171.16 | -502.30 | -15.45 | 53.90 |
| AUAACUUUACGUGUGUGACCUAUUA | -161.75 | -493.21 | -8.85 | 41.37 | -163.09 | -469.15 | -17.65 | 59.55 |
| GUUCUAUACUCUUGAAGUUGAUUAC | -179.98 | -530.68 | -15.47 | 53.06 | -181.07 | -514.61 | -21.54 | 64.88 |
| CCCUGCACUUUAACUGAAUUGUUUA | -182.04 | -523.34 | -19.80 | 61.24 | -183.88 | -510.40 | -25.66 | 72.85 |
| UAACCAUACUGAAUACCUUUUGACG | -190.91 | -552.01 | -19.79 | 59.99 | -192.48 | -538.47 | -25.55 | 70.85 |
| UCCACACGGUAGUAAAAUUAGGCUU | -177.26 | -534.07 | -11.69 | 46.16 | -178.99 | -509.66 | -21.00 | 64.12 |
| UUCCAAAAGGAGUUAUGAGUUGCGA | -164.44 | -499.21 | -9.69 | 42.92 | -166.09 | -474.62 | -18.95 | 61.92 |

| | | | | | | | | |
|---|---|---|---|---|---|---|---|---|
| AAUAUCUCUCAUGCGCCAAGCUACA | -187.57 | -556.51 | -15.05 | 51.61 | -188.65 | -538.28 | -21.78 | 64.13 |
| UAGUAUAUCGCAGCAUCAUACAGGC | -186.08 | -551.70 | -15.05 | 51.73 | -187.26 | -533.32 | -21.93 | 64.63 |
| UGGAUUCUACUCAACCUUAGUCUGG | -185.49 | -531.22 | -20.81 | 62.71 | -187.43 | -518.74 | -26.62 | 74.07 |
| CGGAAUCCAUGUUACUUCGGCUAUC | -196.21 | -575.53 | -17.79 | 55.74 | -197.99 | -556.79 | -25.38 | 69.48 |
| CUGGUCUGGAUCUGAGAACUUCAGG | -181.88 | -508.53 | -24.24 | 70.29 | -184.97 | -497.09 | -30.88 | 83.84 |
| ACAGCGAAUGGACCUACGUGGCCUU | -196.62 | -555.11 | -24.54 | 68.11 | -199.71 | -541.20 | -31.94 | 82.04 |
| AGCAAGUCGAGCAGGGCCUACGUUU | -192.85 | -547.60 | -23.09 | 65.97 | -195.41 | -534.48 | -29.72 | 78.59 |
| GCGAGCGACAGGUUACUUGGCUGAU | -201.88 | -576.28 | -23.23 | 64.81 | -204.43 | -561.62 | -30.32 | 77.69 |
| AAAGGUGUCGCGGAGAGUCGUGCUG | -190.04 | -532.82 | -24.87 | 69.96 | -193.37 | -519.70 | -32.26 | 84.43 |
| AUGGGUGGGAGCCUCGGUAGCAGCC | -198.82 | -568.37 | -22.62 | 64.16 | -202.16 | -549.84 | -31.70 | 80.95 |
| CAGUGGGCUCCUGGGCGUGCUGGUC | -180.37 | -529.04 | -16.37 | 54.74 | -182.55 | -509.44 | -24.63 | 70.97 |
| GCCAACUCCGUCGCCGUUCGUGCGC | -209.04 | -592.72 | -25.30 | 67.43 | -211.98 | -578.16 | -32.75 | 80.61 |
| ACGGGUCCCCGCACCGCACCGCCAG | -207.17 | -579.43 | -27.55 | 71.85 | -210.95 | -565.07 | -35.78 | 86.76 |
| UUAUGUAUUAAGUUAUAUAGUAGUAGUAGU | -195.74 | -605.98 | -7.89 | 39.02 | -196.87 | -574.45 | -18.79 | 57.44 |
| AUUGAUAUCCUUUUCUAUUCAUCUUUCAUU | -212.97 | -635.34 | -16.01 | 51.30 | -215.26 | -608.07 | -26.76 | 69.00 |
| AAAGUACAUCAACAUAGAGAAUUGCAUUUC | -215.38 | -662.02 | -10.16 | 42.16 | -217.13 | -627.34 | -22.65 | 61.72 |
| CUUAAGAUAUGAGAACUUCAACUAAUGUGU | -205.36 | -622.74 | -12.31 | 45.83 | -206.95 | -594.23 | -22.73 | 63.19 |
| CUCAACUUGCGGUAAAUAAAUCGCUUAAUC | -213.28 | -649.13 | -12.05 | 45.10 | -214.79 | -618.98 | -22.91 | 62.44 |
| UAUUGAGAACAAGUGUCCGAUUAGCAGAAA | -212.12 | -618.06 | -20.52 | 58.74 | -215.18 | -594.71 | -30.82 | 76.30 |
| GUCAUACGACUGAGUGCAACAUUGUUCAAA | -218.61 | -646.13 | -18.30 | 54.50 | -221.30 | -619.58 | -29.23 | 72.28 |
| AACCUGCAACAUGGAGUUUUUGUCUCAUGC | -218.33 | -662.02 | -13.10 | 46.47 | -220.33 | -630.32 | -24.93 | 65.10 |
| CCGUGCGGUGUGUACGUUUUAUUCAUCAUA | -234.77 | -682.50 | -23.19 | 60.53 | -237.64 | -659.65 | -33.15 | 75.96 |
| GUUCACGUCCGAAAGCUCGAAAAAGGAUAC | -223.36 | -648.52 | -22.32 | 60.44 | -226.22 | -626.66 | -31.96 | 76.11 |
| AGUCUGGUCUGGAUCUGAGAACUUCAGGCU | -215.29 | -622.13 | -22.43 | 61.58 | -219.10 | -597.87 | -33.76 | 80.85 |
| UCGGAGAAAUCACUGAGCUGCCUGAGAAGA | -223.98 | -668.85 | -16.64 | 51.50 | -227.29 | -635.77 | -30.20 | 72.89 |
| CUUCAACGGAUCAGGUAGGACUGUGGUGGG | -209.14 | -606.73 | -21.06 | 59.99 | -212.82 | -582.09 | -32.38 | 79.71 |
| ACGCCCACAGGAUUAGGCUGGCCCACAUUG | -231.14 | -685.65 | -18.59 | 53.91 | -233.61 | -657.98 | -29.64 | 70.88 |
| GUUAUUCCGCAGUCCGAUGGCAGCAGGCUC | -230.52 | -710.69 | -10.20 | 41.88 | -233.03 | -669.60 | -25.45 | 64.26 |
| UCAGUAGGCGUGACGCAGAGCUGGCGAUGG | -228.63 | -666.92 | -21.89 | 59.18 | -232.16 | -640.33 | -33.66 | 77.87 |
| CGCGCCACGUGUGAUCUACAGCCGUUCGGC | -241.27 | -704.25 | -22.95 | 59.50 | -243.94 | -680.59 | -32.96 | 74.52 |
| GACCUGACGUGGACCGCUCCUGGGCGUGGU | -233.88 | -664.36 | -27.93 | 68.08 | -238.22 | -642.72 | -38.97 | 85.73 |
| GCCCCUCCACUGGCCGACGGCAGCAGGCUC | -253.44 | -711.42 | -32.90 | 72.85 | -258.19 | -691.88 | -43.70 | 88.99 |
| CGCCGCUGCCGACUGGAGGAGCGCGGGACG | -244.68 | -712.72 | -23.74 | 60.30 | -249.74 | -678.96 | -39.26 | 83.62 |

[a] All duplexes consist of the denoted RNA strand and its complementary DNA strand.

**Table S4.** Corrected Thermodynamic Parameters for RNA/PSMOE Duplexes

| RNA Sequence[a] | MMPBSA | | | | MMGBSA | | | |
|---|---|---|---|---|---|---|---|---|
| | $\Delta H°$ (kcal/mol) | $\Delta S°$ (cal/mol/K) | $\Delta G°_{37}$ (kcal/mol) | $T_m$ (°C) | $\Delta H°$ (kcal/mol) | $\Delta S°$ (cal/mol/K) | $\Delta G°_{37}$ (kcal/mol) | $T_m$ (°C) |
| CCGG | -40.67 | -116.21 | -4.65 | 23.16 | -40.86 | -115.24 | -5.13 | 26.63 |
| CGCG | -36.06 | -103.65 | -3.93 | 16.04 | -36.24 | -102.50 | -4.47 | 20.19 |
| GCGC | -36.84 | -107.51 | -3.51 | 13.40 | -37.02 | -105.55 | -4.29 | 19.22 |
| AGCCG | -46.44 | -132.95 | -5.22 | 28.39 | -46.56 | -132.01 | -5.64 | 31.05 |
| CACAG | -41.16 | -117.05 | -4.88 | 24.92 | -41.28 | -116.63 | -5.13 | 26.69 |
| CGUGC | -47.16 | -136.49 | -4.85 | 26.21 | -47.27 | -134.88 | -5.46 | 29.98 |
| GCACG | -43.21 | -122.87 | -5.12 | 27.10 | -43.43 | -121.97 | -5.62 | 30.53 |
| CGGCU | -43.09 | -125.28 | -4.26 | 21.33 | -43.22 | -123.43 | -4.96 | 25.99 |
| GGUGG | -46.45 | -133.83 | -4.96 | 26.72 | -46.69 | -131.93 | -5.79 | 32.03 |
| CCGCGG | -57.33 | -156.06 | -8.96 | 50.56 | -57.90 | -156.88 | -9.27 | 52.25 |
| CGAUCG | -54.06 | -153.16 | -6.58 | 37.15 | -54.09 | -153.26 | -6.58 | 37.13 |
| CGCGCG | -56.88 | -156.41 | -8.40 | 47.39 | -57.30 | -157.10 | -8.60 | 48.47 |
| CGGCCG | -59.16 | -164.55 | -8.15 | 45.59 | -59.53 | -164.55 | -8.52 | 47.59 |
| CGUACG | -52.01 | -146.80 | -6.50 | 36.67 | -52.27 | -146.13 | -6.97 | 39.50 |
| GACGUC | -57.12 | -164.18 | -6.22 | 35.20 | -57.19 | -162.88 | -6.69 | 37.76 |
| GCAUGC | -52.71 | -150.95 | -5.92 | 33.31 | -52.80 | -149.99 | -6.30 | 35.52 |
| GCCGGC | -60.42 | -168.89 | -8.06 | 44.93 | -60.83 | -168.33 | -8.65 | 48.04 |
| GCGAGC | -60.31 | -171.86 | -7.03 | 39.48 | -60.36 | -171.27 | -7.27 | 40.71 |
| GCGCGC | -56.65 | -157.80 | -7.73 | 43.59 | -57.09 | -157.35 | -8.31 | 46.86 |
| GCUAGC | -56.74 | -163.12 | -6.17 | 34.90 | -56.63 | -162.60 | -6.22 | 35.19 |
| GGAUCC | -52.98 | -151.58 | -5.99 | 33.75 | -53.16 | -150.26 | -6.58 | 37.16 |
| GGCGCC | -57.37 | -159.30 | -7.99 | 44.95 | -57.91 | -158.67 | -8.72 | 49.08 |
| GGGACC | -54.80 | -153.49 | -7.22 | 40.83 | -55.30 | -152.38 | -8.06 | 45.68 |
| GUGAAC | -50.12 | -144.97 | -5.18 | 28.71 | -50.20 | -143.37 | -5.76 | 32.16 |
| UCAUGA | -47.22 | -136.93 | -4.77 | 25.74 | -47.17 | -135.87 | -5.05 | 27.45 |
| UGAUCA | -43.93 | -130.30 | -3.54 | 17.11 | -43.82 | -128.25 | -4.06 | 20.33 |
| CCCGGG | -61.26 | -167.33 | -9.38 | 52.02 | -61.81 | -168.09 | -9.70 | 53.63 |
| ACCGCA | -47.44 | -133.96 | -5.91 | 32.87 | -47.76 | -133.02 | -6.52 | 36.81 |
| AGUUGC | -52.85 | -154.52 | -4.95 | 27.86 | -52.80 | -152.59 | -5.50 | 30.93 |
| AUGCGC | -52.02 | -149.24 | -5.76 | 32.32 | -52.17 | -147.90 | -6.32 | 35.59 |
| CCAACG | -53.26 | -152.62 | -5.95 | 33.52 | -53.15 | -152.47 | -5.89 | 33.17 |
| CGGACG | -55.02 | -152.64 | -7.70 | 43.61 | -55.41 | -152.70 | -8.07 | 45.75 |
| CGGUGC | -56.23 | -157.79 | -7.32 | 41.27 | -56.64 | -156.93 | -7.99 | 45.06 |
| CGUGCC | -55.02 | -155.66 | -6.77 | 38.22 | -55.39 | -154.33 | -7.55 | 42.69 |
| CGUUGA | -44.73 | -131.98 | -3.81 | 19.10 | -44.72 | -129.72 | -4.50 | 23.43 |
| CGUUGC | -52.85 | -150.63 | -6.16 | 34.71 | -53.01 | -149.71 | -6.60 | 37.28 |
| CGUUGU | -47.97 | -138.49 | -5.04 | 27.52 | -48.06 | -137.07 | -5.57 | 30.80 |
| GCCUGC | -55.41 | -156.38 | -6.93 | 39.13 | -55.65 | -155.78 | -7.36 | 41.56 |
| UGCGCA | -47.47 | -134.29 | -5.84 | 32.43 | -47.85 | -132.96 | -6.63 | 37.54 |
| UGUUGC | -44.24 | -130.95 | -3.64 | 17.88 | -44.27 | -128.32 | -4.50 | 23.25 |
| CAAUCG | -50.66 | -144.13 | -5.98 | 33.56 | -50.62 | -144.29 | -5.89 | 32.98 |
| CACGGC | -56.08 | -156.57 | -7.54 | 42.55 | -56.47 | -156.14 | -8.07 | 45.55 |
| CGAUUG | -54.22 | -155.75 | -5.93 | 33.49 | -54.05 | -155.64 | -5.80 | 32.73 |
| GCACCG | -54.16 | -150.63 | -7.46 | 42.31 | -54.51 | -150.65 | -7.81 | 44.33 |
| GGCACG | -54.49 | -152.07 | -7.35 | 41.59 | -54.90 | -151.59 | -7.90 | 44.81 |
| CCCAGGG | -70.04 | -191.81 | -10.58 | 55.88 | -70.49 | -193.13 | -10.62 | 55.95 |
| CAAAAAG | -57.15 | -162.44 | -6.79 | 38.30 | -56.86 | -163.64 | -6.13 | 34.72 |
| AAUACCG | -59.00 | -167.83 | -6.97 | 39.18 | -59.04 | -167.44 | -7.14 | 40.10 |
| AGCCGUG | -62.72 | -180.23 | -6.84 | 38.42 | -62.92 | -178.17 | -7.69 | 42.68 |
| AGCUUCA | -60.24 | -175.79 | -5.74 | 32.86 | -60.06 | -174.19 | -6.06 | 34.47 |
| GGACUUA | -59.68 | -172.60 | -6.18 | 35.05 | -59.64 | -171.23 | -6.56 | 37.01 |
| ACGUAUG | -58.58 | -172.38 | -5.15 | 29.71 | -58.39 | -170.24 | -5.61 | 32.07 |
| CACGGCU | -64.76 | -182.62 | -8.15 | 44.83 | -65.05 | -181.87 | -8.67 | 47.42 |

| | | | | | | | | |
|---|---|---|---|---|---|---|---|---|
| CAUACGU | -56.44 | -159.79 | -6.90 | 38.93 | -56.63 | -159.16 | -7.29 | 41.10 |
| UAAGUCC | -57.21 | -163.73 | -6.45 | 36.45 | -57.29 | -162.74 | -6.84 | 38.53 |
| UGAAGCU | -58.47 | -169.16 | -6.03 | 34.24 | -58.37 | -168.05 | -6.28 | 35.52 |
| AAAAAAAA | -55.80 | -161.18 | -5.84 | 33.06 | -55.41 | -161.59 | -5.32 | 30.22 |
| UAGAUCUA | -66.45 | -192.32 | -6.83 | 38.28 | -66.26 | -191.24 | -6.98 | 38.98 |
| UCUAUAGA | -66.66 | -192.92 | -6.86 | 38.39 | -66.47 | -191.87 | -6.99 | 39.02 |
| CAAAAAAG | -65.46 | -185.51 | -7.95 | 43.75 | -65.14 | -187.02 | -7.16 | 39.89 |
| CAAGCUUG | -71.96 | -202.09 | -9.31 | 49.34 | -71.94 | -203.12 | -8.97 | 47.74 |
| CAUCGAUG | -72.38 | -204.19 | -9.08 | 48.19 | -72.44 | -204.37 | -9.09 | 48.20 |
| CGAUAUCG | -70.27 | -199.28 | -8.49 | 45.75 | -70.32 | -198.99 | -8.63 | 46.42 |
| CGUCGACG | -73.12 | -204.01 | -9.87 | 51.72 | -73.54 | -203.72 | -10.38 | 54.00 |
| GAAGCUUC | -72.66 | -208.56 | -8.01 | 43.30 | -72.50 | -208.01 | -8.01 | 43.34 |
| GAUCGAUC | -67.76 | -193.17 | -7.88 | 43.16 | -67.92 | -191.95 | -8.41 | 45.71 |
| GAUGCAUC | -67.43 | -192.73 | -7.69 | 42.28 | -67.59 | -191.29 | -8.29 | 45.15 |
| GGAAUUCC | -68.19 | -194.38 | -7.94 | 43.39 | -68.38 | -193.07 | -8.52 | 46.18 |
| GGACGUCC | -73.18 | -203.44 | -10.11 | 52.82 | -73.85 | -202.38 | -11.11 | 57.37 |
| GGAGCUCC | -73.02 | -204.21 | -9.72 | 51.00 | -73.45 | -203.65 | -10.32 | 53.73 |
| GGUAUACC | -71.52 | -205.58 | -7.78 | 42.40 | -71.77 | -203.03 | -8.83 | 47.13 |
| GUACGUAC | -69.01 | -196.71 | -8.03 | 43.75 | -69.35 | -194.69 | -8.99 | 48.28 |
| GUAGCUAC | -70.58 | -201.91 | -7.99 | 43.39 | -70.68 | -200.55 | -8.50 | 45.77 |
| GUUGCAAC | -66.10 | -187.78 | -7.89 | 43.38 | -66.37 | -186.42 | -8.59 | 46.77 |
| AAGCGUAG | -70.43 | -197.96 | -9.07 | 48.44 | -70.65 | -197.84 | -9.32 | 49.62 |
| AAUCCAGU | -65.54 | -187.60 | -7.39 | 40.96 | -65.56 | -186.68 | -7.69 | 42.43 |
| ACAUAUGU | -62.77 | -182.02 | -6.34 | 35.95 | -62.67 | -180.52 | -6.71 | 37.78 |
| ACCUAGUC | -68.79 | -195.51 | -8.18 | 44.48 | -68.96 | -194.50 | -8.66 | 46.76 |
| ACGACCUC | -74.03 | -209.42 | -9.11 | 48.04 | -74.27 | -208.51 | -9.63 | 50.38 |
| AGAGAGAG | -74.18 | -210.84 | -8.82 | 46.73 | -74.16 | -210.58 | -8.88 | 47.02 |
| AGAGCUCU | -72.15 | -207.83 | -7.72 | 42.07 | -72.05 | -206.64 | -7.99 | 43.26 |
| AGCGUAAG | -68.62 | -193.76 | -8.56 | 46.31 | -68.93 | -192.80 | -9.16 | 49.18 |
| AGUCCUGA | -68.12 | -194.62 | -7.79 | 42.71 | -68.40 | -192.67 | -8.67 | 46.86 |
| AUGCGCAU | -65.09 | -185.13 | -7.71 | 42.56 | -65.36 | -183.68 | -8.42 | 46.10 |
| CACGGCUC | -78.28 | -219.20 | -10.32 | 52.66 | -78.63 | -218.88 | -10.77 | 54.55 |
| CCAUAUGG | -71.94 | -202.09 | -9.29 | 49.25 | -72.09 | -202.33 | -9.37 | 49.57 |
| CGCGUAUA | -66.76 | -193.47 | -6.79 | 38.07 | -66.85 | -191.04 | -7.63 | 42.03 |
| CGCUGUAA | -70.93 | -201.00 | -8.62 | 46.27 | -71.09 | -200.34 | -8.98 | 47.92 |
| CGUCGUCC | -69.77 | -193.57 | -9.76 | 51.91 | -70.42 | -192.73 | -10.68 | 56.27 |
| CUAGUGGA | -69.05 | -193.10 | -9.19 | 49.28 | -69.43 | -192.70 | -9.70 | 51.68 |
| CUCACGGC | -74.52 | -206.78 | -10.41 | 53.91 | -75.01 | -206.71 | -10.93 | 56.20 |
| CUGAGUCC | -72.33 | -203.50 | -9.24 | 48.93 | -72.61 | -202.97 | -9.69 | 50.97 |
| GAAUAUUC | -63.01 | -183.39 | -6.16 | 35.04 | -62.81 | -181.98 | -6.40 | 36.20 |
| GACUAGUC | -72.19 | -207.19 | -7.96 | 43.12 | -72.21 | -205.84 | -8.40 | 45.09 |
| GAGUACUC | -69.51 | -199.41 | -7.70 | 42.15 | -69.64 | -197.68 | -8.36 | 45.23 |
| GAUUAAUC | -64.55 | -189.49 | -5.81 | 33.44 | -64.26 | -187.65 | -6.09 | 34.73 |
| GCAUAUGC | -68.84 | -196.59 | -7.90 | 43.15 | -68.95 | -195.42 | -8.37 | 45.35 |
| GCCAGUUA | -68.60 | -194.47 | -8.31 | 45.13 | -68.81 | -193.55 | -8.81 | 47.48 |
| GGACCUCG | -74.92 | -209.14 | -10.08 | 52.29 | -75.32 | -208.89 | -10.56 | 54.41 |
| GGAGCACG | -70.95 | -195.66 | -10.30 | 54.25 | -71.59 | -195.54 | -10.97 | 57.35 |
| GGUGCCAA | -69.21 | -194.08 | -9.04 | 48.54 | -69.75 | -192.72 | -10.01 | 53.12 |
| GUCGAACA | -65.19 | -184.67 | -7.95 | 43.75 | -65.49 | -183.49 | -8.60 | 47.00 |
| GUCUAGAC | -69.45 | -198.14 | -8.03 | 43.71 | -69.62 | -196.79 | -8.62 | 46.44 |
| UAGGCCUA | -75.36 | -214.12 | -8.99 | 47.30 | -75.46 | -213.42 | -9.30 | 48.67 |
| UAUGCAUA | -65.75 | -191.46 | -6.39 | 36.23 | -65.61 | -189.62 | -6.82 | 38.26 |
| UGAGCUCA | -67.16 | -193.40 | -7.20 | 39.99 | -67.15 | -191.98 | -7.63 | 42.04 |
| AAUGUCGC | -69.01 | -196.33 | -8.15 | 44.30 | -69.20 | -195.14 | -8.70 | 46.91 |
| ACUGGAUU | -64.80 | -188.82 | -6.26 | 35.60 | -64.73 | -186.64 | -6.87 | 38.52 |
| CAACAGCA | -65.13 | -184.84 | -7.83 | 43.17 | -65.22 | -184.39 | -8.05 | 44.28 |
| CUACGCUU | -68.52 | -194.89 | -8.11 | 44.16 | -68.61 | -194.18 | -8.42 | 45.63 |
| CUUACGCU | -72.88 | -208.36 | -8.29 | 44.51 | -72.83 | -207.68 | -8.44 | 45.23 |

| | | | | | | | | |
|---|---|---|---|---|---|---|---|---|
| GACUAGGU | -72.77 | -208.51 | -8.13 | 43.83 | -72.81 | -207.19 | -8.58 | 45.84 |
| GAGCCGUG | -77.15 | -214.49 | -10.66 | 54.40 | -77.59 | -214.56 | -11.08 | 56.15 |
| GAGGUCGU | -64.99 | -180.80 | -8.94 | 48.80 | -65.64 | -179.64 | -9.95 | 53.90 |
| GCCGUGAG | -76.27 | -212.72 | -10.33 | 53.11 | -76.72 | -212.38 | -10.88 | 55.50 |
| GCGACAUU | -66.44 | -190.71 | -7.32 | 40.60 | -66.58 | -188.99 | -7.99 | 43.80 |
| UAACUGGC | -67.09 | -188.48 | -8.66 | 47.03 | -67.43 | -187.83 | -9.21 | 49.67 |
| UCCACUAG | -65.90 | -183.95 | -8.87 | 48.28 | -66.21 | -184.03 | -9.16 | 49.67 |
| UGCUGUUG | -69.15 | -196.59 | -8.21 | 44.56 | -69.32 | -195.55 | -8.70 | 46.87 |
| UGUUCGAC | -68.70 | -196.21 | -7.88 | 43.08 | -68.96 | -194.37 | -8.71 | 46.97 |
| UUACAGCG | -70.28 | -197.92 | -8.93 | 47.82 | -70.45 | -197.84 | -9.12 | 48.69 |
| UUGGCACC | -70.25 | -196.91 | -9.21 | 49.17 | -70.79 | -195.63 | -10.15 | 53.55 |
| CAAAAAAAG | -72.65 | -205.64 | -8.90 | 47.32 | -72.35 | -207.11 | -8.15 | 43.95 |
| CAAACAAAG | -74.53 | -209.12 | -9.70 | 50.64 | -74.50 | -210.28 | -9.31 | 48.90 |
| CAAAUAAAG | -74.55 | -212.44 | -8.70 | 46.15 | -74.26 | -213.17 | -8.18 | 43.90 |
| CAAAGAAAG | -74.04 | -207.45 | -9.73 | 50.86 | -74.01 | -208.71 | -9.31 | 48.97 |
| AAAAAAAAA | -64.74 | -186.81 | -6.83 | 38.32 | -64.29 | -187.26 | -6.24 | 35.49 |
| AUAACUGGC | -72.21 | -203.33 | -9.18 | 48.67 | -72.64 | -202.09 | -9.99 | 52.36 |
| AUCUAUCCG | -75.82 | -216.67 | -8.66 | 45.81 | -75.91 | -215.37 | -9.15 | 47.93 |
| CGCUGUUAC | -80.84 | -229.64 | -9.65 | 49.32 | -81.09 | -228.20 | -10.35 | 52.18 |
| GCCAGUUAA | -74.58 | -209.45 | -9.65 | 50.40 | -74.95 | -208.71 | -10.25 | 53.06 |
| CAACAGCAA | -73.44 | -206.72 | -9.36 | 49.30 | -73.68 | -206.35 | -9.71 | 50.87 |
| CAACAGCAU | -73.07 | -205.39 | -9.40 | 49.51 | -73.37 | -204.89 | -9.85 | 51.55 |
| CGCUGUUAG | -82.78 | -233.44 | -10.41 | 52.10 | -82.94 | -233.16 | -10.66 | 53.12 |
| CUAACAGCG | -78.02 | -219.29 | -10.04 | 51.47 | -78.23 | -219.20 | -10.28 | 52.46 |
| GUAACAGCG | -81.35 | -228.78 | -10.43 | 52.47 | -81.62 | -228.40 | -10.81 | 54.02 |
| UUAACUGGC | -75.09 | -211.70 | -9.47 | 49.49 | -75.45 | -210.64 | -10.15 | 52.50 |
| AUGAGCUCAU | -89.07 | -256.61 | -9.52 | 47.63 | -89.02 | -254.60 | -10.10 | 49.81 |
| GCGAAUUCGC | -92.49 | -261.62 | -11.38 | 54.03 | -92.75 | -260.49 | -12.00 | 56.29 |
| GCGAAAAGCG | -88.95 | -247.04 | -12.37 | 58.64 | -89.37 | -247.51 | -12.64 | 59.62 |
| AUCAAUCAUA | -77.50 | -225.98 | -7.45 | 40.57 | -77.26 | -223.89 | -7.85 | 42.24 |
| UUGUAGUCAU | -83.66 | -241.49 | -8.80 | 45.52 | -83.78 | -238.69 | -9.79 | 49.41 |
| GAAAUGAAAG | -83.92 | -239.04 | -9.82 | 49.52 | -83.83 | -238.75 | -9.82 | 49.52 |
| CCAACUUCUU | -88.15 | -251.96 | -10.05 | 49.74 | -87.99 | -251.47 | -10.04 | 49.73 |
| AUCGUCUGGA | -86.06 | -243.53 | -10.56 | 52.10 | -86.50 | -241.59 | -11.61 | 56.20 |
| AGCGUAAGUC | -86.61 | -245.04 | -10.65 | 52.36 | -87.07 | -243.09 | -11.71 | 56.49 |
| CGAUCUGCGA | -89.27 | -253.01 | -10.84 | 52.60 | -89.60 | -251.41 | -11.66 | 55.69 |
| UGGCGAGCAC | -94.53 | -264.89 | -12.41 | 57.43 | -95.23 | -263.01 | -13.70 | 62.10 |
| GAUGCGCUCG | -90.54 | -252.19 | -12.36 | 58.21 | -91.18 | -251.37 | -13.25 | 61.54 |
| GGGACCGCCU | -95.21 | -263.81 | -13.43 | 61.07 | -96.16 | -262.29 | -14.85 | 66.23 |
| AAAAAAAAAA | -73.96 | -213.45 | -7.79 | 42.25 | -73.45 | -213.84 | -7.16 | 39.53 |
| CGGCAAGCGC | -92.50 | -254.80 | -13.52 | 62.19 | -93.39 | -254.28 | -14.57 | 66.05 |
| UAGGUUAUAA | -84.19 | -247.57 | -7.44 | 40.26 | -84.05 | -243.67 | -8.51 | 44.34 |
| CGUACACAUGC | -96.95 | -273.49 | -12.17 | 56.01 | -97.58 | -271.06 | -13.56 | 60.91 |
| CCAUUGCUACC | -97.98 | -276.71 | -12.20 | 55.90 | -98.42 | -275.00 | -13.17 | 59.29 |
| CCAUCGCUACC | -97.35 | -270.78 | -13.41 | 60.43 | -98.07 | -269.86 | -14.41 | 63.96 |
| ACGUAUUAUGC | -94.74 | -273.17 | -10.06 | 48.85 | -94.90 | -269.97 | -11.21 | 52.93 |
| GCAUAAUACGU | -90.23 | -257.32 | -10.46 | 50.99 | -90.49 | -255.21 | -11.38 | 54.41 |
| CGCGAAUUCGCG | -107.96 | -300.92 | -14.67 | 62.15 | -108.72 | -299.66 | -15.83 | 65.86 |
| CUGACAAGUGUC | -107.52 | -304.17 | -13.23 | 57.45 | -108.06 | -301.69 | -14.53 | 61.66 |
| UUUUAUAAUAAA | -92.79 | -272.23 | -8.40 | 43.23 | -92.36 | -269.36 | -8.86 | 44.89 |
| AUUGGAUACAAA | -101.00 | -292.50 | -10.33 | 48.97 | -101.16 | -288.42 | -11.76 | 53.74 |
| CAACCAACCAAC | -103.61 | -289.16 | -13.97 | 60.84 | -104.27 | -288.13 | -14.95 | 64.08 |
| CUUCCUUCCUUC | -109.15 | -311.41 | -12.61 | 55.15 | -109.22 | -309.70 | -13.21 | 57.07 |
| GGAACAAGAUGC | -107.40 | -303.09 | -13.44 | 58.17 | -107.94 | -300.97 | -14.64 | 62.03 |
| GGAACUUGAUGC | -104.35 | -295.22 | -12.83 | 56.79 | -104.96 | -292.47 | -14.29 | 61.63 |
| AAUGGAUUACAA | -97.89 | -280.72 | -10.87 | 51.23 | -98.23 | -277.37 | -12.24 | 56.00 |
| GUCAGGAAUCUG | -108.96 | -308.04 | -13.47 | 57.94 | -109.42 | -306.02 | -14.55 | 61.39 |
| UUGUAAUCCAUU | -95.92 | -279.59 | -9.24 | 45.89 | -96.07 | -274.87 | -10.86 | 51.48 |

| | | | | | | | | |
|---|---|---|---|---|---|---|---|---|
| UUUGUAUCCAAU | -99.63 | -290.14 | -9.68 | 46.99 | -99.65 | -285.91 | -11.02 | 51.49 |
| CGCAUGGGUACGC | -119.77 | -333.96 | -16.24 | 64.21 | -121.07 | -330.42 | -18.64 | 71.31 |
| AGCCUAACUCAGC | -115.20 | -325.94 | -14.16 | 58.84 | -115.84 | -322.97 | -15.72 | 63.56 |
| CAUAUGGCCAUAUG | -119.38 | -336.58 | -15.04 | 60.65 | -120.23 | -333.20 | -16.94 | 66.25 |
| GUAUACCGGUAUAC | -127.65 | -365.96 | -14.20 | 56.68 | -128.33 | -360.33 | -16.63 | 63.33 |
| ACAUUAUUAUUACA | -110.38 | -324.76 | -9.70 | 46.02 | -110.21 | -319.24 | -11.24 | 50.71 |
| UACUAACAUUAACUA | -118.40 | -342.22 | -12.31 | 52.77 | -118.80 | -336.69 | -14.43 | 58.94 |
| AUACUUACUGAUUAG | -127.30 | -368.84 | -12.96 | 53.35 | -127.42 | -363.82 | -14.63 | 57.91 |
| GUACACUGUCUUAUA | -124.30 | -359.42 | -12.88 | 53.53 | -124.86 | -352.88 | -15.47 | 60.76 |
| GUAUGAGAGACUUUA | -122.84 | -355.67 | -12.58 | 52.93 | -123.46 | -348.69 | -15.37 | 60.76 |
| UUCUACCUAUGUGAU | -127.40 | -367.59 | -13.45 | 54.66 | -127.84 | -361.91 | -15.64 | 60.66 |
| AGUAGUAAUCACACC | -127.23 | -365.28 | -14.00 | 56.19 | -128.17 | -358.25 | -17.11 | 64.76 |
| AUCGUCUCGGUAUAA | -127.69 | -365.19 | -14.48 | 57.43 | -128.51 | -359.32 | -17.12 | 64.71 |
| ACGACAGGUUUACCA | -133.35 | -379.20 | -15.80 | 60.02 | -134.49 | -372.90 | -18.89 | 68.23 |
| CUUUCAUGUCCGCAU | -132.86 | -378.30 | -15.58 | 59.53 | -133.43 | -374.28 | -17.41 | 64.37 |
| UGGAUGUGUGAACAC | -132.11 | -379.53 | -14.46 | 56.66 | -133.22 | -371.48 | -18.07 | 66.25 |
| ACCCCGCAAUACAUG | -124.41 | -351.40 | -15.47 | 60.87 | -125.51 | -346.54 | -18.09 | 68.30 |
| GCAGUGGAUGUGAGA | -132.19 | -377.95 | -15.02 | 58.14 | -133.32 | -370.69 | -18.41 | 67.18 |
| GGUCCUUACUUGGUG | -135.17 | -389.22 | -14.52 | 56.32 | -135.82 | -382.69 | -17.19 | 63.26 |
| CGCCUCAUGCUCAUC | -135.65 | -383.04 | -16.91 | 62.54 | -136.49 | -379.41 | -18.87 | 67.67 |
| AAAUAGCCGGGCCGC | -141.37 | -398.58 | -17.81 | 63.73 | -142.72 | -392.92 | -20.91 | 71.60 |
| CCAGCCAGUCUCUCC | -137.79 | -386.84 | -17.87 | 64.65 | -138.97 | -382.74 | -20.32 | 71.02 |
| GACGACAAGACCGCG | -135.48 | -382.24 | -16.99 | 62.79 | -136.66 | -377.22 | -19.72 | 69.99 |
| CAGCCUCGUCGCAGC | -141.43 | -391.33 | -20.12 | 69.80 | -142.86 | -388.99 | -22.27 | 75.24 |
| CUCGCGGUCGAAGCG | -134.51 | -384.83 | -15.21 | 58.25 | -135.44 | -378.29 | -18.17 | 66.01 |
| GCGUCGGUCCGGGCU | -142.86 | -399.02 | -19.16 | 66.93 | -144.69 | -393.09 | -22.83 | 76.22 |
| CAACUUGAUAUUAUUA | -132.49 | -386.56 | -12.65 | 51.88 | -132.59 | -380.02 | -14.79 | 57.44 |
| CAUAUUGGCCAAUAUG | -141.95 | -406.13 | -16.05 | 59.14 | -142.59 | -400.73 | -18.36 | 64.91 |
| GUAUAACCGGUUAUAC | -141.81 | -407.57 | -15.46 | 57.68 | -142.64 | -400.37 | -18.53 | 65.33 |
| CGCGUACGCGUACGCG | -149.32 | -414.56 | -20.81 | 69.64 | -151.29 | -409.30 | -24.41 | 78.40 |
| CAACUUGAUAUUAAUA | -131.92 | -385.76 | -12.34 | 51.13 | -131.98 | -379.06 | -14.47 | 56.70 |
| UAUGUAUAUUUUGUAAUCAG | -166.16 | -487.79 | -14.95 | 53.40 | -166.77 | -475.87 | -19.25 | 62.45 |
| UUCAAGUUAAACAUUCUAUC | -167.88 | -489.06 | -16.27 | 55.95 | -168.51 | -478.78 | -20.09 | 63.98 |
| UGAUUCUACCUAUGUGAUUU | -161.92 | -475.23 | -14.59 | 53.11 | -162.73 | -462.66 | -19.31 | 63.27 |
| GAGAUUGUUUCCCUUUCAAA | -175.58 | -519.42 | -14.56 | 51.71 | -176.32 | -504.36 | -19.96 | 62.43 |
| AUGCAAUGCUACAUAUUCGC | -175.21 | -506.40 | -18.22 | 59.02 | -176.33 | -495.53 | -22.72 | 68.19 |
| CCACUAUACCAUCUAUGUAC | -170.76 | -489.08 | -19.15 | 61.59 | -172.12 | -479.54 | -23.46 | 70.68 |
| CCAUCAUUGUGUCUACCUCA | -175.48 | -505.09 | -18.91 | 60.38 | -176.82 | -494.31 | -23.58 | 69.94 |
| CGGGACCAACUAAAGGAAAU | -179.82 | -514.02 | -20.47 | 62.91 | -181.29 | -504.25 | -24.97 | 71.96 |
| UAGUGGCGAUUAGAUUCUGC | -176.56 | -507.18 | -19.34 | 61.10 | -178.16 | -495.71 | -24.49 | 71.60 |
| AGCUGCAGUGGAUGUGAGAA | -182.22 | -527.38 | -18.73 | 59.11 | -183.59 | -514.82 | -23.99 | 69.44 |
| UACUUCCAGUGCUCAGCGUA | -178.76 | -520.97 | -17.26 | 56.65 | -179.86 | -507.93 | -22.41 | 66.87 |
| CAGUGAGACAGCAAUGGUCG | -181.10 | -521.10 | -19.56 | 60.88 | -182.62 | -509.40 | -24.70 | 71.12 |
| CGAGCUUAUCCCUAUCCCUC | -175.78 | -511.20 | -17.31 | 57.11 | -176.76 | -499.39 | -21.95 | 66.49 |
| CGUACUAGCGUUGGUCAUGG | -187.60 | -535.19 | -21.69 | 64.12 | -189.23 | -525.09 | -26.45 | 73.33 |
| AAGGCGAGUCAGGCUCAGUG | -181.02 | -511.05 | -22.59 | 67.05 | -183.30 | -500.82 | -28.05 | 78.09 |
| ACCGACGACGCUGAUCCGAU | -180.25 | -511.08 | -21.82 | 65.58 | -182.12 | -501.58 | -26.63 | 75.32 |
| AGCAGUCCGCCACACCCUGA | -181.88 | -509.72 | -23.87 | 69.52 | -184.40 | -500.27 | -29.32 | 80.56 |
| CAGCCUCGUUCGCACAGCCC | -178.62 | -497.83 | -24.30 | 71.10 | -181.24 | -489.30 | -29.56 | 81.98 |
| GUGGUGGGCCGUGCGCUCUG | -195.15 | -552.07 | -24.01 | 67.35 | -197.67 | -540.14 | -30.23 | 79.08 |
| GUCCACGCCCGGUGCGACGG | -194.04 | -542.12 | -25.98 | 71.40 | -196.76 | -532.67 | -31.63 | 82.19 |
| GAUAUAGCAAAAUUCUAAGUUAAUA | -209.05 | -621.23 | -16.47 | 52.33 | -209.99 | -601.49 | -23.53 | 64.16 |
| AUAACUUUACGUGUGUGACCUAUUA | -213.59 | -631.20 | -17.92 | 54.31 | -215.08 | -610.38 | -25.87 | 67.48 |
| GUUCUAUACUCUUGAAGUUGAUUAC | -222.69 | -658.08 | -18.68 | 54.75 | -223.80 | -638.37 | -25.90 | 66.23 |
| CCCUGCACUUUAACUGAAUUGUUUA | -213.62 | -628.97 | -18.64 | 55.48 | -215.01 | -609.79 | -25.97 | 67.68 |
| UAACCAUACUGAAUACCUUUUGACG | -201.28 | -604.27 | -13.96 | 48.73 | -202.27 | -581.89 | -21.89 | 62.32 |
| UCCACACGGUAGUAAAAUUAGGCUU | -221.05 | -641.72 | -22.12 | 60.38 | -223.24 | -623.01 | -30.10 | 73.46 |
| UUCCAAAAGGAGUUAUGAGUUGCGA | -228.77 | -667.38 | -21.88 | 59.16 | -230.50 | -648.77 | -29.38 | 70.96 |

| | | | | | | | | |
|---|---|---|---|---|---|---|---|---|
| AAUAUCUCUCAUGCGCCAAGCUACA | -205.90 | -602.79 | -19.03 | 56.90 | -207.78 | -583.60 | -26.87 | 70.49 |
| UAGUAUAUCGCAGCAUCAUACAGGC | -214.11 | -625.14 | -20.32 | 58.19 | -216.03 | -606.12 | -28.14 | 71.31 |
| UGGAUUCUACUCAACCUUAGUCUGG | -222.23 | -653.67 | -19.59 | 56.21 | -223.62 | -634.23 | -27.01 | 68.11 |
| CGGAAUCCAUGUUACUUCGGCUAUC | -202.34 | -607.57 | -13.99 | 48.73 | -203.55 | -584.07 | -22.49 | 63.23 |
| CUGGUCUGGAUCUGAGAACUUCAGG | -232.90 | -672.59 | -24.40 | 62.61 | -235.24 | -654.42 | -32.37 | 75.11 |
| ACAGCGAAUGGACCUACGUGGCCUU | -224.32 | -641.75 | -25.37 | 65.28 | -227.44 | -623.38 | -34.19 | 79.78 |
| AGCAAGUCGAGCAGGGCCUACGUUU | -223.80 | -653.88 | -21.09 | 58.43 | -225.87 | -633.45 | -29.50 | 71.96 |
| GCGAGCGACAGGUUACUUGGCUGAU | -234.88 | -678.06 | -24.68 | 62.82 | -237.38 | -659.24 | -33.01 | 75.78 |
| AAAGGUGUCGCGGAGAGUCGUGCUG | -230.65 | -658.01 | -26.66 | 66.50 | -233.78 | -640.36 | -35.27 | 80.30 |
| AUGGGUGGGAGCCUCGGUAGCAGCC | -241.69 | -689.45 | -27.96 | 67.02 | -245.17 | -670.07 | -37.45 | 81.59 |
| CAGUGGGCUCCUGGGCGUGCUGGUC | -231.47 | -669.02 | -24.07 | 62.28 | -234.59 | -647.13 | -33.98 | 77.93 |
| GCCAACUCCGUCGCCGUUCGUGCGC | -239.03 | -674.81 | -29.84 | 70.35 | -242.24 | -660.16 | -37.59 | 82.45 |
| ACGGGUCCCCGCACCGCACCGCCAG | -246.84 | -690.85 | -32.68 | 73.58 | -250.79 | -675.88 | -41.27 | 86.70 |
| UUAUGUAUUAAGUUAUAUAGUAGUAGUAGU | -249.72 | -775.15 | -9.42 | 40.49 | -249.93 | -739.16 | -20.80 | 55.62 |
| AUUGAUAUCCUUUUCUAUUCAUCUUUCAUU | -251.03 | -755.31 | -16.88 | 50.19 | -252.55 | -725.14 | -27.75 | 65.30 |
| AAAGUACAUCAACAUAGAGAAUUGCAUUUC | -252.16 | -746.82 | -20.65 | 55.24 | -254.47 | -718.91 | -31.61 | 70.74 |
| CUUAAGAUAUGAGAACUUCAACUAAUGUGU | -244.53 | -742.70 | -14.29 | 47.01 | -245.94 | -710.16 | -25.79 | 63.20 |
| CUCAACUUGCGGUAAAUAAAUCGCUUAAUC | -269.18 | -799.30 | -21.39 | 54.97 | -270.97 | -771.41 | -31.83 | 68.78 |
| UAUUGAGAACAAGUGUCCGAUUAGCAGAAA | -261.04 | -786.13 | -17.34 | 50.24 | -262.65 | -754.22 | -28.85 | 65.64 |
| GUCAUACGACUGAGUGCAACAUUGUUCAAA | -251.58 | -785.94 | -7.94 | 38.59 | -253.51 | -739.57 | -24.24 | 60.13 |
| AACCUGCAACAUGGAGUUUUUGUCUCAUGC | -259.80 | -777.44 | -18.79 | 52.21 | -262.09 | -745.08 | -31.12 | 68.95 |
| CCGUGCGGUGUGUACGUUUUAUUCAUCAUA | -268.74 | -789.99 | -23.85 | 58.21 | -272.13 | -758.97 | -36.85 | 75.73 |
| GUUCACGUCCGAAAGCUCGAAAAAGGAUAC | -267.13 | -789.15 | -22.49 | 56.55 | -269.09 | -762.66 | -32.67 | 70.20 |
| AGUCUGGUCUGGAUCUGAGAACUUCAGGCU | -270.91 | -796.27 | -24.06 | 58.31 | -273.88 | -767.02 | -36.10 | 74.38 |
| UCGGAGAAAUCACUGAGCUGCCUGAGAAGA | -252.65 | -757.39 | -17.86 | 51.41 | -254.40 | -727.42 | -28.90 | 66.75 |
| CUUCAACGGAUCAGGUAGGACUGUGGUGGG | -272.17 | -798.04 | -24.78 | 59.14 | -275.96 | -766.02 | -38.50 | 77.47 |
| ACGCCCACAGGAUUAGGCUGGCCCACAUUG | -267.46 | -794.35 | -21.21 | 54.86 | -270.71 | -760.03 | -35.10 | 73.43 |
| GUUAUUCCGCAGUCCGAUGGCAGCAGGCUC | -265.69 | -773.17 | -26.01 | 61.38 | -269.26 | -745.42 | -38.18 | 78.15 |
| UCAGUAGGCGUGACGCAGAGCUGGCGAUGG | -275.30 | -799.98 | -27.31 | 62.16 | -278.81 | -772.61 | -39.30 | 78.15 |
| CGCGCCACGUGUGAUCUACAGCCGUUCGGC | -276.12 | -824.24 | -20.61 | 53.51 | -278.31 | -791.91 | -32.81 | 69.18 |
| GACCUGACGUGGACCGCUCCUGGGCGUGGU | -259.06 | -765.26 | -21.83 | 56.31 | -262.26 | -733.85 | -34.76 | 74.25 |
| GCCCCUCCACUGGCCGACGGCAGCAGGCUC | -291.48 | -835.02 | -32.62 | 67.33 | -295.50 | -810.53 | -44.24 | 82.19 |
| CGCCGCUGCCGACUGGAGGAGCGCGGGACG | -297.15 | -843.64 | -35.62 | 70.50 | -301.95 | -819.33 | -47.96 | 86.15 |

[a] All duplexes consist of the denoted RNA strand and its complementary DNA strand.

**Table S5.** Corrected Thermodynamic Parameters for RNA/PSDNA-PSMOE Duplexes

| RNA Sequence[a] | MMPBSA | | | | MMGBSA | | | |
|---|---|---|---|---|---|---|---|---|
| | $\Delta H°$ (kcal/mol) | $\Delta S°$ (cal/mol/K) | $\Delta G°_{37}$ (kcal/mol) | $T_m$ (°C) | $\Delta H°$ (kcal/mol) | $\Delta S°$ (cal/mol/K) | $\Delta G°_{37}$ (kcal/mol) | $T_m$ (°C) |
| CCGG | -35.42 | -105.91 | -2.59 | 5.82 | -35.46 | -103.34 | -3.43 | 11.94 |
| CGCG | -34.49 | -98.55 | -3.93 | 15.17 | -34.66 | -97.76 | -4.35 | 18.54 |
| GCGC | -34.93 | -103.06 | -2.98 | 8.27 | -35.06 | -100.83 | -3.80 | 14.47 |
| AGCCG | -38.12 | -109.39 | -4.21 | 19.08 | -38.39 | -107.87 | -4.95 | 24.59 |
| CACAG | -39.62 | -116.13 | -3.61 | 15.62 | -39.60 | -114.60 | -4.08 | 18.79 |
| CGUGC | -46.64 | -137.07 | -4.14 | 21.78 | -46.60 | -135.06 | -4.74 | 25.37 |
| GCACG | -43.22 | -125.19 | -4.41 | 22.34 | -43.24 | -124.04 | -4.79 | 24.84 |
| CGGCU | -45.14 | -130.92 | -4.56 | 23.89 | -45.20 | -129.46 | -5.07 | 27.17 |
| GGUGG | -46.22 | -135.06 | -4.35 | 22.90 | -46.41 | -132.46 | -5.35 | 29.17 |
| CCGCGG | -54.86 | -153.03 | -7.43 | 42.01 | -55.25 | -152.69 | -7.92 | 44.85 |
| CGAUCG | -49.39 | -139.08 | -6.28 | 35.28 | -49.47 | -139.41 | -6.25 | 35.11 |
| CGCGCG | -53.53 | -146.04 | -8.26 | 47.22 | -53.93 | -147.27 | -8.28 | 47.23 |
| CGGCCG | -53.35 | -152.31 | -6.13 | 34.58 | -53.53 | -151.16 | -6.66 | 37.65 |
| CGUACG | -49.55 | -140.42 | -6.02 | 33.71 | -49.70 | -139.97 | -6.31 | 35.52 |
| GACGUC | -49.59 | -142.33 | -5.47 | 30.38 | -49.73 | -141.07 | -5.99 | 33.57 |
| GCAUGC | -48.74 | -139.33 | -5.55 | 30.78 | -48.92 | -138.19 | -6.08 | 34.06 |
| GCCGGC | -52.42 | -144.03 | -7.77 | 44.37 | -53.02 | -143.80 | -8.44 | 48.46 |
| GCGAGC | -55.66 | -159.74 | -6.14 | 34.70 | -55.71 | -158.69 | -6.52 | 36.79 |
| GCGCGC | -53.31 | -151.02 | -6.49 | 36.66 | -53.63 | -149.82 | -7.18 | 40.70 |
| GCUAGC | -52.70 | -151.41 | -5.76 | 32.40 | -52.67 | -150.72 | -5.95 | 33.46 |
| GGAUCC | -48.88 | -139.14 | -5.74 | 31.95 | -49.08 | -138.16 | -6.25 | 35.13 |
| GGCGCC | -54.92 | -152.18 | -7.74 | 43.86 | -55.39 | -151.96 | -8.28 | 46.98 |
| GGGACC | -52.24 | -147.50 | -6.52 | 36.78 | -52.65 | -146.15 | -7.34 | 41.73 |
| GUGAAC | -47.32 | -138.36 | -4.43 | 23.66 | -47.34 | -136.40 | -5.06 | 27.52 |
| UCAUGA | -40.47 | -120.77 | -3.03 | 12.20 | -40.47 | -118.09 | -3.86 | 17.67 |
| UGAUCA | -43.37 | -128.68 | -3.49 | 16.53 | -43.25 | -126.66 | -3.99 | 19.67 |
| CCCGGG | -51.22 | -136.54 | -8.89 | 51.85 | -51.98 | -137.59 | -9.33 | 54.52 |
| ACCGCA | -49.09 | -139.31 | -5.91 | 32.98 | -49.29 | -138.60 | -6.32 | 35.56 |
| AGUUGC | -50.29 | -148.43 | -4.27 | 23.54 | -50.20 | -146.11 | -4.91 | 27.15 |
| AUGCGC | -49.26 | -143.06 | -4.91 | 27.01 | -49.31 | -141.35 | -5.49 | 30.48 |
| CCAACG | -46.42 | -130.06 | -6.10 | 34.02 | -46.58 | -130.33 | -6.18 | 34.54 |
| CGGACG | -53.27 | -149.19 | -7.03 | 39.78 | -53.52 | -149.19 | -7.27 | 41.20 |
| CGGUGC | -52.51 | -148.01 | -6.62 | 37.42 | -52.89 | -146.88 | -7.36 | 41.82 |
| CGUGCC | -50.89 | -141.96 | -6.88 | 39.03 | -51.29 | -141.50 | -7.43 | 42.40 |
| CGUUGA | -47.61 | -136.61 | -5.27 | 28.86 | -47.70 | -135.66 | -5.64 | 31.21 |
| CGUUGC | -51.47 | -148.38 | -5.48 | 30.66 | -51.51 | -147.20 | -5.88 | 32.98 |
| CGUUGU | -48.23 | -141.38 | -4.40 | 23.78 | -48.15 | -139.67 | -4.85 | 26.43 |
| GCCUGC | -52.57 | -148.08 | -6.66 | 37.64 | -52.81 | -147.66 | -7.03 | 39.83 |
| UGCGCA | -44.02 | -124.78 | -5.34 | 28.71 | -44.35 | -123.61 | -6.03 | 33.39 |
| UGUUGC | -46.77 | -138.90 | -3.71 | 19.21 | -46.69 | -136.36 | -4.42 | 23.44 |
| CAAUCG | -48.88 | -139.54 | -5.62 | 31.21 | -48.76 | -139.81 | -5.42 | 29.95 |
| CACGGC | -54.39 | -152.29 | -7.18 | 40.59 | -54.69 | -152.01 | -7.57 | 42.88 |
| CGAUUG | -42.92 | -128.73 | -3.01 | 13.40 | -42.74 | -126.35 | -3.57 | 16.79 |
| GCACCG | -50.08 | -140.00 | -6.68 | 37.82 | -50.42 | -139.68 | -7.12 | 40.52 |
| GGCACG | -53.14 | -148.46 | -7.12 | 40.34 | -53.45 | -148.35 | -7.46 | 42.34 |
| CCCAGGG | -57.28 | -156.96 | -8.62 | 48.60 | -57.85 | -157.22 | -9.11 | 51.33 |
| CAAAAAG | -55.60 | -160.27 | -5.92 | 33.50 | -55.26 | -160.64 | -5.46 | 31.00 |
| AAUACCG | -59.43 | -171.57 | -6.24 | 35.37 | -59.27 | -170.85 | -6.31 | 35.70 |
| AGCCGUG | -59.99 | -168.39 | -7.79 | 43.52 | -60.38 | -167.61 | -8.42 | 46.86 |
| AGCUUCA | -55.93 | -165.11 | -4.75 | 27.28 | -55.69 | -163.07 | -5.14 | 29.32 |
| GGACUUA | -57.47 | -169.00 | -5.08 | 29.26 | -57.31 | -166.85 | -5.59 | 31.85 |
| ACGUAUG | -56.80 | -163.90 | -5.99 | 33.94 | -56.75 | -162.83 | -6.27 | 35.46 |
| CACGGCU | -61.70 | -173.60 | -7.89 | 43.84 | -62.02 | -172.90 | -8.43 | 46.64 |

| | | | | | | | | |
|---|---|---|---|---|---|---|---|---|
| CAUACGU | -53.63 | -158.38 | -4.53 | 25.73 | -53.51 | -155.92 | -5.18 | 29.21 |
| UAAGUCC | -56.51 | -165.21 | -5.29 | 30.22 | -56.34 | -163.63 | -5.61 | 31.88 |
| UGAAGCU | -57.67 | -168.12 | -5.55 | 31.68 | -57.44 | -166.95 | -5.69 | 32.40 |
| AAAAAAAA | -58.44 | -173.68 | -4.60 | 26.97 | -57.80 | -172.73 | -4.25 | 25.09 |
| UAGAUCUA | -63.49 | -187.99 | -5.21 | 30.56 | -63.26 | -185.11 | -5.88 | 33.71 |
| UCUAUAGA | -62.48 | -184.11 | -5.41 | 31.40 | -62.19 | -182.05 | -5.75 | 33.02 |
| CAAAAAAG | -63.50 | -181.18 | -7.34 | 40.85 | -63.11 | -182.41 | -6.56 | 37.01 |
| CAAGCUUG | -63.00 | -177.46 | -7.99 | 44.22 | -63.10 | -177.65 | -8.03 | 44.41 |
| CAUCGAUG | -66.78 | -190.94 | -7.58 | 41.84 | -66.74 | -190.33 | -7.74 | 42.57 |
| CGAUAUCG | -64.71 | -183.14 | -7.94 | 43.75 | -64.82 | -182.87 | -8.13 | 44.69 |
| CGUCGACG | -70.19 | -199.30 | -8.41 | 45.38 | -70.47 | -197.88 | -9.13 | 48.73 |
| GAAGCUUC | -64.63 | -189.92 | -5.75 | 33.17 | -64.37 | -187.80 | -6.15 | 35.07 |
| GAUCGAUC | -62.76 | -183.67 | -5.82 | 33.41 | -62.74 | -181.01 | -6.63 | 37.35 |
| GAUGCAUC | -65.61 | -188.80 | -7.08 | 39.51 | -65.66 | -187.23 | -7.62 | 42.11 |
| GGAAUUCC | -64.47 | -183.29 | -7.65 | 42.34 | -64.65 | -182.30 | -8.13 | 44.74 |
| GGACGUCC | -68.57 | -191.30 | -9.27 | 49.77 | -69.18 | -190.12 | -10.25 | 54.47 |
| GGAGCUCC | -69.49 | -193.46 | -9.52 | 50.80 | -69.92 | -193.28 | -10.01 | 53.08 |
| GGUAUACC | -65.23 | -186.32 | -7.47 | 41.42 | -65.58 | -184.15 | -8.49 | 46.43 |
| GUACGUAC | -65.56 | -190.13 | -6.62 | 37.27 | -65.73 | -187.27 | -7.68 | 42.38 |
| GUAGCUAC | -62.75 | -178.71 | -7.35 | 40.98 | -62.99 | -177.31 | -8.03 | 44.40 |
| GUUGCAAC | -64.14 | -182.87 | -7.45 | 41.39 | -64.33 | -181.57 | -8.04 | 44.33 |
| AAGCGUAG | -66.88 | -189.69 | -8.07 | 44.18 | -67.08 | -188.76 | -8.57 | 46.57 |
| AAUCCAGU | -64.22 | -184.82 | -6.93 | 38.80 | -64.18 | -183.73 | -7.22 | 40.23 |
| ACAUAUGU | -61.18 | -180.68 | -5.17 | 30.14 | -60.92 | -178.38 | -5.62 | 32.29 |
| ACCUAGUC | -64.13 | -181.58 | -7.84 | 43.30 | -64.35 | -180.77 | -8.31 | 45.67 |
| ACGACCUC | -70.88 | -205.54 | -7.16 | 39.66 | -70.86 | -203.33 | -7.83 | 42.66 |
| AGAGAGAG | -65.39 | -183.08 | -8.64 | 47.19 | -65.66 | -183.09 | -8.90 | 48.47 |
| AGAGCUCU | -69.87 | -200.47 | -7.72 | 42.24 | -69.76 | -199.77 | -7.83 | 42.76 |
| AGCGUAAG | -61.93 | -174.25 | -7.92 | 43.96 | -62.29 | -173.37 | -8.55 | 47.25 |
| AGUCCUGA | -63.51 | -182.61 | -6.90 | 38.69 | -63.67 | -180.68 | -7.66 | 42.48 |
| AUGCGCAU | -59.86 | -175.34 | -5.50 | 31.63 | -60.10 | -171.56 | -6.92 | 38.88 |
| CACGGCUC | -70.40 | -196.25 | -9.56 | 50.83 | -70.87 | -195.81 | -10.17 | 53.64 |
| CCAUAUGG | -61.15 | -170.67 | -8.24 | 45.80 | -61.55 | -170.31 | -8.76 | 48.51 |
| CGCGUAUA | -64.29 | -187.83 | -6.06 | 34.62 | -64.19 | -185.59 | -6.65 | 37.46 |
| CGCUGUAA | -66.36 | -188.70 | -7.87 | 43.24 | -66.47 | -187.99 | -8.19 | 44.83 |
| CGUCGUCC | -66.92 | -187.86 | -8.69 | 47.19 | -67.42 | -186.63 | -9.56 | 51.45 |
| CUAGUGGA | -61.58 | -171.44 | -8.43 | 46.72 | -62.06 | -170.90 | -9.08 | 50.16 |
| CUCACGGC | -72.28 | -201.60 | -9.78 | 51.47 | -72.65 | -201.58 | -10.15 | 53.14 |
| CUGAGUCC | -68.33 | -193.75 | -8.26 | 44.93 | -68.52 | -192.88 | -8.72 | 47.11 |
| GAAUAUUC | -59.95 | -178.19 | -4.71 | 27.75 | -59.58 | -175.89 | -5.06 | 29.38 |
| GACUAGUC | -65.50 | -187.26 | -7.45 | 41.30 | -65.64 | -185.92 | -8.01 | 44.00 |
| GAGUACUC | -67.70 | -196.90 | -6.66 | 37.48 | -67.74 | -194.33 | -7.49 | 41.34 |
| GAUUAAUC | -61.41 | -182.08 | -4.96 | 29.15 | -61.09 | -179.62 | -5.41 | 31.29 |
| GCAUAUGC | -62.15 | -177.60 | -7.10 | 39.71 | -62.36 | -176.04 | -7.79 | 43.25 |
| GCCAGUUA | -65.66 | -188.18 | -7.32 | 40.66 | -65.81 | -186.53 | -7.99 | 43.89 |
| GGACCUCG | -72.79 | -203.55 | -9.69 | 50.93 | -73.09 | -203.58 | -9.98 | 52.22 |
| GGAGCACG | -68.93 | -189.50 | -10.18 | 54.20 | -69.50 | -189.87 | -10.64 | 56.35 |
| GGUGCCAA | -69.25 | -191.24 | -9.97 | 53.06 | -69.91 | -190.84 | -10.75 | 56.76 |
| GUCGAACA | -65.82 | -188.32 | -7.44 | 41.22 | -65.90 | -187.18 | -7.87 | 43.30 |
| GUCUAGAC | -64.93 | -185.18 | -7.53 | 41.69 | -65.12 | -183.84 | -8.14 | 44.70 |
| UAGGCCUA | -64.71 | -187.21 | -6.68 | 37.57 | -64.81 | -184.97 | -7.47 | 41.44 |
| UAUGCAUA | -60.78 | -181.43 | -4.54 | 27.02 | -60.47 | -178.37 | -5.18 | 30.09 |
| UGAGCUCA | -67.05 | -192.10 | -7.50 | 41.41 | -67.02 | -191.26 | -7.73 | 42.53 |
| AAUGUCGC | -66.40 | -193.35 | -6.46 | 36.55 | -66.36 | -191.03 | -7.14 | 39.76 |
| ACUGGAUU | -65.34 | -191.39 | -6.01 | 34.40 | -65.21 | -188.98 | -6.62 | 37.30 |
| CAACAGCA | -63.92 | -179.75 | -8.20 | 45.16 | -64.11 | -179.69 | -8.41 | 46.21 |
| CUACGCUU | -60.14 | -171.80 | -6.88 | 38.67 | -60.37 | -170.20 | -7.61 | 42.50 |
| CUUACGCU | -64.46 | -181.77 | -8.11 | 44.64 | -64.60 | -181.68 | -8.28 | 45.48 |

| | | | | | | | | |
|---|---|---|---|---|---|---|---|---|
| GACUAGGU | -65.51 | -188.09 | -7.20 | 40.06 | -65.63 | -186.41 | -7.84 | 43.19 |
| GAGCCGUG | -70.13 | -194.80 | -9.74 | 51.72 | -70.64 | -194.47 | -10.36 | 54.62 |
| GAGGUCGU | -68.22 | -195.73 | -7.55 | 41.55 | -68.41 | -193.74 | -8.35 | 45.35 |
| GCCGUGAG | -66.68 | -182.22 | -10.19 | 54.89 | -67.36 | -182.60 | -10.76 | 57.61 |
| GCGACAUU | -66.80 | -196.92 | -5.75 | 33.30 | -66.83 | -193.07 | -6.98 | 38.98 |
| UAACUGGC | -67.44 | -191.86 | -7.96 | 43.59 | -67.62 | -190.76 | -8.48 | 46.09 |
| UCCACUAG | -62.70 | -177.08 | -7.80 | 43.28 | -62.90 | -176.56 | -8.17 | 45.14 |
| UGCUGUUG | -60.53 | -172.15 | -7.16 | 40.12 | -60.82 | -170.66 | -7.91 | 44.08 |
| UGUUCGAC | -65.64 | -189.34 | -6.94 | 38.83 | -65.76 | -187.23 | -7.72 | 42.58 |
| UUACAGCG | -64.89 | -183.39 | -8.03 | 44.22 | -65.05 | -182.97 | -8.33 | 45.70 |
| UUGGCACC | -62.71 | -176.22 | -8.09 | 44.75 | -63.29 | -174.49 | -9.20 | 50.53 |
| CAAAAAAAG | -71.47 | -205.26 | -7.83 | 42.63 | -71.00 | -206.01 | -7.14 | 39.55 |
| CAAACAAAG | -71.26 | -201.79 | -8.71 | 46.64 | -71.19 | -202.23 | -8.50 | 45.66 |
| CAAAUAAAG | -69.54 | -199.07 | -7.83 | 42.77 | -69.28 | -199.32 | -7.49 | 41.20 |
| CAAAGAAAG | -71.24 | -200.81 | -8.98 | 47.93 | -71.15 | -201.76 | -8.60 | 46.15 |
| AAAAAAAAA | -63.88 | -187.73 | -5.69 | 32.82 | -63.28 | -187.20 | -5.25 | 30.72 |
| AUAACUGGC | -71.70 | -204.41 | -8.33 | 44.87 | -71.91 | -202.89 | -9.02 | 47.96 |
| AUCUAUCCG | -76.85 | -220.86 | -8.38 | 44.51 | -76.74 | -219.78 | -8.61 | 45.49 |
| CGCUGUUAC | -75.03 | -212.63 | -9.11 | 47.90 | -75.30 | -211.37 | -9.78 | 50.83 |
| GCCAGUUAA | -68.42 | -192.28 | -8.81 | 47.55 | -68.84 | -191.21 | -9.57 | 51.17 |
| CAACAGCAA | -72.86 | -208.38 | -8.26 | 44.42 | -72.87 | -207.39 | -8.58 | 45.85 |
| CAACAGCAU | -71.34 | -204.04 | -8.09 | 43.78 | -71.46 | -202.62 | -8.64 | 46.31 |
| CGCUGUUAG | -72.79 | -207.06 | -8.60 | 45.93 | -73.01 | -205.68 | -9.25 | 48.84 |
| CUAACAGCG | -77.91 | -218.22 | -10.26 | 52.45 | -78.02 | -218.93 | -10.16 | 51.98 |
| GUAACAGCG | -76.64 | -214.96 | -10.00 | 51.56 | -76.91 | -214.80 | -10.32 | 52.94 |
| UUAACUGGC | -70.56 | -204.41 | -7.19 | 39.79 | -70.64 | -201.88 | -8.06 | 43.71 |
| AUGAGCUCAU | -78.50 | -228.25 | -7.74 | 41.71 | -78.56 | -225.07 | -8.79 | 46.03 |
| GCGAAUUCGC | -83.51 | -238.25 | -9.65 | 48.90 | -83.72 | -236.44 | -10.42 | 51.96 |
| GCGAAAAGCG | -87.73 | -246.06 | -11.45 | 55.27 | -87.94 | -246.23 | -11.61 | 55.87 |
| AUCAAUCAUA | -76.12 | -221.12 | -7.57 | 41.16 | -75.86 | -219.57 | -7.79 | 42.10 |
| UUGUAGUCAU | -73.30 | -213.95 | -6.97 | 38.74 | -73.51 | -209.93 | -8.43 | 45.08 |
| GAAAUGAAAG | -80.74 | -230.58 | -9.26 | 47.71 | -80.62 | -230.19 | -9.26 | 47.71 |
| CCAACUUCUU | -73.98 | -212.83 | -8.01 | 43.17 | -74.05 | -210.93 | -8.67 | 46.07 |
| AUCGUCUGGA | -83.76 | -237.10 | -10.25 | 51.28 | -84.05 | -235.82 | -10.94 | 54.04 |
| AGCGUAAGUC | -82.57 | -233.49 | -10.19 | 51.24 | -83.10 | -231.31 | -11.39 | 56.12 |
| CGAUCUGCGA | -85.39 | -242.69 | -10.16 | 50.62 | -85.65 | -241.12 | -10.90 | 53.53 |
| UGGCGAGCAC | -79.94 | -226.68 | -9.68 | 49.56 | -80.71 | -223.15 | -11.53 | 57.35 |
| GAUGCGCUCG | -87.43 | -243.02 | -12.09 | 57.92 | -87.94 | -242.96 | -12.62 | 59.92 |
| GGGACCGCCU | -89.30 | -246.81 | -12.79 | 60.23 | -90.25 | -245.52 | -14.14 | 65.40 |
| AAAAAAAAAA | -73.04 | -214.13 | -6.66 | 37.40 | -72.34 | -213.70 | -6.09 | 35.01 |
| CGGCAAGCGC | -85.22 | -235.89 | -12.09 | 58.51 | -86.07 | -234.76 | -13.29 | 63.30 |
| UAGGUUAUAA | -77.33 | -224.11 | -7.85 | 42.25 | -77.30 | -221.74 | -8.56 | 45.21 |
| CGUACACAUGC | -87.26 | -246.80 | -10.75 | 52.63 | -87.96 | -243.79 | -12.39 | 58.99 |
| CCAUUGCUACC | -89.58 | -250.78 | -11.84 | 56.39 | -90.13 | -249.70 | -12.72 | 59.72 |
| CCAUCGCUACC | -87.94 | -242.81 | -12.67 | 60.14 | -88.79 | -242.08 | -13.74 | 64.27 |
| ACGUAUUAUGC | -86.62 | -249.45 | -9.29 | 47.07 | -86.81 | -246.53 | -10.38 | 51.26 |
| GCAUAAUACGU | -92.05 | -267.16 | -9.23 | 46.22 | -91.85 | -264.74 | -9.78 | 48.24 |
| CGCGAAUUCGCG | -106.28 | -293.88 | -15.18 | 64.32 | -106.89 | -294.46 | -15.61 | 65.63 |
| CUGACAAGUGUC | -92.13 | -263.49 | -10.45 | 50.62 | -92.76 | -259.29 | -12.39 | 57.75 |
| UUUUAUAAUAAA | -83.45 | -242.73 | -8.21 | 43.22 | -83.21 | -240.66 | -8.60 | 44.77 |
| AUUGGAUACAAA | -91.56 | -264.48 | -9.57 | 47.52 | -91.80 | -260.77 | -10.96 | 52.59 |
| CAACCAACCAAC | -101.97 | -285.26 | -13.54 | 59.75 | -102.47 | -284.60 | -14.25 | 62.11 |
| CUUCCUUCCUUC | -103.12 | -291.06 | -12.89 | 57.24 | -103.20 | -290.99 | -12.99 | 57.57 |
| GGAACAAGAUGC | -99.75 | -279.89 | -12.98 | 58.29 | -100.37 | -278.21 | -14.13 | 62.24 |
| GGAACUUGAUGC | -98.22 | -275.67 | -12.76 | 57.86 | -98.86 | -273.87 | -13.96 | 62.06 |
| AAUGGAUUACAA | -86.28 | -246.75 | -9.79 | 49.03 | -86.80 | -243.23 | -11.40 | 55.27 |
| GUCAGGAAUCUG | -97.15 | -275.61 | -11.71 | 54.32 | -97.63 | -273.08 | -12.98 | 58.77 |
| UUGUAAUCCAUU | -91.45 | -266.64 | -8.79 | 44.71 | -91.55 | -262.32 | -10.23 | 49.91 |

| | | | | | | | | |
|---|---|---|---|---|---|---|---|---|
| UUUGUAUCCAAU | -94.02 | -273.08 | -9.37 | 46.52 | -94.01 | -269.66 | -10.42 | 50.24 |
| CGCAUGGGUACGC | -112.64 | -315.99 | -14.69 | 61.06 | -113.81 | -312.00 | -17.09 | 68.56 |
| AGCCUAACUCAGC | -106.82 | -300.53 | -13.65 | 59.00 | -107.43 | -298.55 | -14.88 | 62.99 |
| CAUAUGGCCAUAUG | -115.46 | -328.42 | -13.65 | 57.24 | -115.96 | -325.17 | -15.16 | 61.79 |
| GUAUACCGGUAUAC | -113.76 | -324.62 | -13.13 | 55.95 | -114.50 | -319.82 | -15.36 | 62.76 |
| ACAUUAUUAUUACA | -97.11 | -284.46 | -8.93 | 44.72 | -97.14 | -279.54 | -10.48 | 50.01 |
| UACUAACAUUAACUA | -112.37 | -333.77 | -8.90 | 43.53 | -112.22 | -326.46 | -11.02 | 49.77 |
| AUACUUACUGAUUAG | -116.12 | -342.17 | -10.05 | 46.55 | -115.95 | -336.08 | -11.76 | 51.50 |
| GUACACUGUCUUAUA | -114.84 | -333.52 | -11.46 | 50.75 | -115.15 | -327.76 | -13.55 | 56.98 |
| GUAUGAGAGACUUUA | -119.48 | -344.48 | -12.69 | 53.71 | -120.00 | -338.81 | -14.97 | 60.30 |
| UUCUACCUAUGUGAU | -113.51 | -323.38 | -13.26 | 56.40 | -114.21 | -319.02 | -15.31 | 62.67 |
| AGUAGUAAUCACACC | -120.89 | -348.76 | -12.78 | 53.76 | -121.57 | -342.26 | -15.47 | 61.45 |
| AUCGUCUCGGUAUAA | -124.26 | -357.91 | -13.30 | 54.73 | -124.65 | -352.84 | -15.27 | 60.23 |
| ACGACAGGUUUACCA | -116.61 | -328.32 | -14.83 | 60.61 | -117.82 | -323.57 | -17.51 | 68.72 |
| CUUUCAUGUCCGCAU | -118.37 | -337.93 | -13.61 | 56.58 | -119.08 | -333.10 | -15.81 | 63.07 |
| UGGAUGUGUGAACAC | -121.07 | -346.09 | -13.78 | 56.61 | -122.23 | -339.00 | -17.14 | 66.33 |
| ACCCCGCAAUACAUG | -117.95 | -335.08 | -14.07 | 58.04 | -118.90 | -329.99 | -16.61 | 65.56 |
| GCAGUGGAUGUGAGA | -110.67 | -331.92 | -7.77 | 40.38 | -111.44 | -319.09 | -12.52 | 54.47 |
| GGUCCUUACUUGGUG | -125.95 | -358.99 | -14.66 | 58.26 | -126.71 | -354.07 | -16.95 | 64.63 |
| CGCCUCAUGCUCAUC | -122.20 | -348.14 | -14.28 | 57.85 | -122.75 | -344.34 | -16.01 | 62.79 |
| AAAUAGCCGGGCCGC | -136.70 | -382.00 | -18.28 | 66.02 | -137.90 | -378.75 | -20.48 | 71.76 |
| CCAGCCAGUCUCUCC | -129.30 | -360.51 | -17.54 | 65.72 | -130.39 | -358.05 | -19.39 | 70.78 |
| GACGACAAGACCGCG | -135.22 | -378.08 | -18.01 | 65.62 | -136.25 | -375.41 | -19.87 | 70.51 |
| CAGCCUCGUCGCAGC | -134.05 | -375.32 | -17.70 | 65.05 | -135.10 | -372.31 | -19.69 | 70.31 |
| CUCGCGGUCGAAGCG | -135.91 | -384.68 | -16.65 | 61.81 | -137.03 | -379.27 | -19.46 | 69.16 |
| GCGUCGGUCCGGGCU | -125.20 | -356.69 | -14.63 | 58.29 | -126.56 | -349.17 | -18.32 | 68.70 |
| CAACUUGAUAUUAUUA | -125.74 | -371.69 | -10.52 | 47.01 | -125.38 | -365.19 | -12.17 | 51.47 |
| CAUAUUGGCCAAUAUG | -132.50 | -378.10 | -15.29 | 58.80 | -133.09 | -373.59 | -17.28 | 64.10 |
| GUAUAACCGGUUAUAC | -129.00 | -372.19 | -13.62 | 54.88 | -129.76 | -365.00 | -16.61 | 62.96 |
| CGCGUACGCGUACGCG | -138.18 | -382.69 | -19.55 | 69.10 | -139.76 | -379.43 | -22.14 | 75.83 |
| CAACUUGAUAUUAAUA | -120.41 | -354.09 | -10.65 | 47.83 | -120.55 | -346.68 | -13.08 | 54.67 |
| UAUGUAUAUUUUGUAAUCAG | -157.25 | -466.58 | -12.61 | 49.33 | -157.37 | -454.93 | -16.34 | 57.46 |
| UUCAAGUUAAACAUUCUAUC | -156.12 | -454.47 | -15.24 | 55.16 | -156.61 | -445.56 | -18.48 | 62.47 |
| UGAUUCUACCUAUGUGAUUU | -152.91 | -457.14 | -11.20 | 46.61 | -153.01 | -444.15 | -15.33 | 55.76 |
| GAGAUUGUUUCCCUUUCAAA | -159.45 | -468.48 | -14.22 | 52.57 | -159.98 | -457.09 | -18.28 | 61.43 |
| AUGCAAUGCUACAUAUUCGC | -159.64 | -463.61 | -15.92 | 56.23 | -160.52 | -453.33 | -19.98 | 65.21 |
| CCACUAUACCAUCUAUGUAC | -141.88 | -417.20 | -12.54 | 50.58 | -142.75 | -405.19 | -17.14 | 61.75 |
| CCAUCAUUGUGUCUACCUCA | -154.99 | -452.44 | -14.73 | 54.18 | -155.97 | -440.77 | -19.32 | 64.56 |
| CGGGACCAACUAAAGGAAAU | -168.97 | -478.87 | -20.52 | 64.84 | -170.52 | -471.03 | -24.50 | 73.37 |
| UAGUGGCGAUUAGAUUCUGC | -151.72 | -459.63 | -9.23 | 42.48 | -152.15 | -442.29 | -15.04 | 55.23 |
| AGCUGCAGUGGAUGUGAGAA | -163.42 | -462.88 | -19.93 | 64.55 | -165.31 | -453.69 | -24.67 | 75.06 |
| UACUUCCAGUGCUCAGCGUA | -161.59 | -466.87 | -16.86 | 58.02 | -162.89 | -455.79 | -21.59 | 68.45 |
| CAGUGAGACAGCAAUGGUCG | -170.79 | -491.48 | -18.43 | 60.07 | -172.15 | -480.76 | -23.12 | 69.91 |
| CGAGCUUAUCCCUAUCCCUC | -173.94 | -496.97 | -19.88 | 62.63 | -174.98 | -489.41 | -23.26 | 69.63 |
| CGUACUAGCGUUGGUCAUGG | -164.08 | -468.80 | -18.75 | 61.81 | -165.80 | -458.36 | -23.71 | 72.69 |
| AAGGCGAGUCAGGCUCAGUG | -170.30 | -481.50 | -21.03 | 65.71 | -172.09 | -473.13 | -25.42 | 75.08 |
| ACCGACGACGCUGAUCCGAU | -167.09 | -472.24 | -20.69 | 65.57 | -168.98 | -463.53 | -25.29 | 75.56 |
| AGCAGUCCGCCACACCCUGA | -164.42 | -466.04 | -19.95 | 64.40 | -166.36 | -456.48 | -24.85 | 75.22 |
| CAGCCUCGUUCGCACAGCCC | -179.52 | -501.86 | -23.94 | 70.15 | -181.45 | -495.60 | -27.82 | 78.05 |
| GUGGUGGGCCGUGCGCUCUG | -171.46 | -484.12 | -21.39 | 66.26 | -174.07 | -472.44 | -27.62 | 79.58 |
| GUCCACGCCCGGUGCGACGG | -177.50 | -501.55 | -22.02 | 66.49 | -179.53 | -492.22 | -26.95 | 76.63 |
| GAUAUAGCAAAAUUCUAAGUUAAUA | -199.88 | -591.36 | -16.55 | 53.22 | -200.56 | -574.79 | -22.37 | 63.44 |
| AUAACUUUACGUGUGUGACCUAUUA | -204.96 | -599.87 | -19.00 | 56.93 | -206.57 | -582.03 | -26.14 | 69.36 |
| GUUCUAUACUCUUGAAGUUGAUUAC | -198.37 | -588.75 | -15.86 | 52.16 | -199.03 | -571.47 | -21.88 | 62.76 |
| CCCUGCACUUUAACUGAAUUGUUUA | -199.57 | -584.00 | -18.53 | 56.69 | -200.57 | -569.27 | -24.09 | 66.61 |
| UAACCAUACUGAAUACCUUUUGACG | -169.27 | -508.51 | -11.63 | 46.48 | -170.29 | -488.80 | -18.76 | 60.84 |
| UCCACACGGUAGUAAAAUUAGGCUU | -200.79 | -589.93 | -17.91 | 55.48 | -202.07 | -572.63 | -24.56 | 67.22 |
| UUCCAAAAGGAGUUAUGAGUUGCGA | -192.19 | -559.52 | -18.73 | 57.87 | -193.69 | -544.30 | -24.96 | 69.46 |

| | | | | | | | | |
|---|---|---|---|---|---|---|---|---|
| AAUAUCUCUCAUGCGCCAAGCUACA | -198.89 | -579.89 | -19.12 | 57.81 | -200.43 | -563.79 | -25.65 | 69.55 |
| UAGUAUAUCGCAGCAUCAUACAGGC | -205.51 | -609.44 | -16.59 | 52.80 | -206.59 | -589.99 | -23.70 | 64.95 |
| UGGAUUCUACUCAACCUUAGUCUGG | -211.50 | -612.61 | -21.59 | 60.62 | -213.22 | -597.09 | -28.12 | 71.78 |
| CGGAAUCCAUGUUACUUCGGCUAUC | -199.45 | -585.14 | -18.06 | 55.87 | -201.15 | -566.54 | -25.52 | 69.17 |
| CUGGUCUGGAUCUGAGAACUUCAGG | -208.48 | -604.25 | -21.16 | 60.25 | -210.31 | -588.14 | -27.99 | 72.08 |
| ACAGCGAAUGGACCUACGUGGCCUU | -196.10 | -573.32 | -18.37 | 56.77 | -198.66 | -552.02 | -27.53 | 73.51 |
| AGCAAGUCGAGCAGGGCCUACGUUU | -210.11 | -599.54 | -24.26 | 65.42 | -212.91 | -583.71 | -31.97 | 78.91 |
| GCGAGCGACAGGUUACUUGGCUGAU | -203.35 | -583.35 | -22.52 | 63.30 | -206.10 | -566.38 | -30.52 | 77.69 |
| AAAGGUGUCGCGGAGAGUCGUGCUG | -207.95 | -601.76 | -21.41 | 60.74 | -210.22 | -584.19 | -29.12 | 74.18 |
| AUGGGUGGGAGCCUCGGUAGCAGCC | -208.76 | -598.18 | -23.32 | 63.97 | -211.58 | -581.08 | -31.44 | 78.23 |
| CAGUGGGCUCCUGGGCGUGCUGGUC | -227.12 | -644.66 | -27.28 | 68.02 | -230.12 | -629.27 | -35.05 | 80.71 |
| GCCAACUCCGUCGCCGUUCGUGCGC | -205.23 | -595.41 | -20.65 | 59.77 | -207.54 | -577.01 | -28.67 | 73.87 |
| ACGGGUCCCCGCACCGCACCGCCAG | -190.22 | -538.63 | -23.25 | 66.72 | -193.82 | -521.66 | -32.11 | 83.99 |
| UUAUGUAUUAAGUUAUAUAGUAGUAGUAGU | -230.51 | -693.73 | -15.46 | 49.34 | -232.40 | -663.82 | -26.61 | 66.18 |
| AUUGAUAUCCUUUUCUAUUCAUCUUUCAUU | -224.02 | -694.61 | -8.69 | 39.87 | -224.67 | -660.77 | -19.83 | 56.36 |
| AAAGUACAUCAACAUAGAGAAUUGCAUUUC | -237.17 | -710.22 | -17.00 | 51.17 | -238.87 | -682.24 | -27.38 | 66.50 |
| CUUAAGAUAUGAGAACUUCAACUAAUGUGU | -235.12 | -714.03 | -13.77 | 46.70 | -236.68 | -681.93 | -25.28 | 63.53 |
| CUCAACUUGCGGUAAAUAAAUCGCUUAAUC | -228.02 | -685.27 | -15.58 | 49.67 | -229.93 | -655.93 | -26.59 | 66.49 |
| UAUUGAGAACAAGUGUCCGAUUAGCAGAAA | -246.92 | -732.41 | -19.87 | 54.56 | -249.22 | -704.35 | -30.87 | 70.41 |
| GUCAUACGACUGAGUGCAACAUUGUUCAAA | -232.17 | -686.85 | -19.24 | 54.81 | -234.83 | -659.18 | -30.48 | 72.06 |
| AACCUGCAACAUGGAGUUUUUGUCUCAUGC | -234.98 | -695.98 | -19.23 | 54.56 | -237.13 | -669.99 | -29.44 | 70.00 |
| CCGUGCGGUGUGUACGUUUUAUUCAUCAUA | -247.86 | -732.18 | -20.88 | 55.91 | -250.77 | -702.83 | -32.89 | 73.27 |
| GUUCACGUCCGAAAGCUCGAAAAAGGAUAC | -222.36 | -684.71 | -10.10 | 41.91 | -223.90 | -649.44 | -22.57 | 60.78 |
| AGUCUGGUCUGGAUCUGAGAACUUCAGGCU | -254.12 | -740.03 | -24.71 | 60.74 | -257.17 | -714.85 | -35.57 | 76.31 |
| UCGGAGAAAUCACUGAGCUGCCUGAGAAGA | -234.98 | -709.29 | -15.10 | 48.59 | -236.94 | -677.60 | -26.88 | 65.99 |
| CUUCAACGGAUCAGGUAGGACUGUGGUGGG | -249.31 | -769.87 | -10.65 | 42.06 | -252.11 | -724.40 | -27.55 | 65.05 |
| ACGCCCACAGGAUUAGGCUGGCCCACAUUG | -249.88 | -717.71 | -27.38 | 65.08 | -254.17 | -692.14 | -39.60 | 83.23 |
| GUUAUUCCGCAGUCCGAUGGCAGCAGGCUC | -256.67 | -744.49 | -25.88 | 62.13 | -259.99 | -719.47 | -36.95 | 77.93 |
| UCAGUAGGCGUGACGCAGAGCUGGCGAUGG | -263.69 | -760.13 | -28.05 | 64.40 | -267.64 | -734.33 | -40.00 | 81.16 |
| CGCGCCACGUGUGAUCUACAGCCGUUCGGC | -245.83 | -732.11 | -18.88 | 53.24 | -248.99 | -698.86 | -32.34 | 72.70 |
| GACCUGACGUGGACCGCUCCUGGGCGUGGU | -262.34 | -740.86 | -32.67 | 71.16 | -266.95 | -719.76 | -43.82 | 87.19 |
| GCCCCUCCACUGGCCGACGGCAGCAGGCUC | -255.50 | -742.38 | -25.37 | 61.53 | -259.81 | -712.37 | -38.98 | 81.09 |
| CGCCGCUGCCGACUGGAGGAGCGCGGGACG | -268.04 | -783.73 | -25.09 | 59.91 | -272.25 | -751.19 | -39.38 | 79.39 |

[a] All duplexes consist of the denoted RNA strand and its complementary DNA strand.

**Table S6.** Corrected Thermodynamic Parameters for RNA/PSMOE-PSDNA Duplexes

| RNA Sequence[a] | MMPBSA | | | | MMGBSA | | | |
|---|---|---|---|---|---|---|---|---|
| | $\Delta H°$ (kcal/mol) | $\Delta S°$ (cal/mol/K) | $\Delta G°_{37}$ (kcal/mol) | $T_m$ (°C) | $\Delta H°$ (kcal/mol) | $\Delta S°$ (cal/mol/K) | $\Delta G°_{37}$ (kcal/mol) | $T_m$ (°C) |
| CCGG | -36.10 | -110.12 | -1.97 | 2.10 | -36.01 | -107.04 | -2.83 | 7.97 |
| CGCG | -35.55 | -105.90 | -2.72 | 6.85 | -35.54 | -103.74 | -3.38 | 11.63 |
| GCGC | -33.38 | -97.12 | -3.27 | 9.32 | -33.60 | -95.30 | -4.06 | 15.61 |
| AGCCG | -44.90 | -131.23 | -4.22 | 21.67 | -44.92 | -129.45 | -4.79 | 25.28 |
| CACAG | -40.64 | -120.00 | -3.44 | 14.94 | -40.56 | -118.26 | -3.90 | 18.02 |
| CGUGC | -39.73 | -114.93 | -4.10 | 19.01 | -39.95 | -113.12 | -4.88 | 24.58 |
| GCACG | -43.61 | -127.48 | -4.09 | 20.41 | -43.66 | -125.62 | -4.71 | 24.49 |
| CGGCU | -43.54 | -124.40 | -4.98 | 26.20 | -43.79 | -123.07 | -5.64 | 30.69 |
| GGUGG | -42.88 | -120.45 | -5.54 | 29.89 | -43.34 | -119.25 | -6.37 | 35.72 |
| CCGCGG | -55.40 | -152.13 | -8.24 | 46.73 | -55.80 | -152.92 | -8.39 | 47.58 |
| CGAUCG | -54.34 | -160.54 | -4.57 | 26.09 | -54.07 | -158.68 | -4.88 | 27.68 |
| CGCGCG | -53.25 | -149.32 | -6.96 | 39.41 | -53.63 | -148.64 | -7.55 | 42.85 |
| CGGCCG | -58.78 | -165.63 | -7.43 | 41.70 | -59.05 | -165.04 | -7.89 | 44.15 |
| CGUACG | -55.88 | -164.49 | -4.89 | 28.02 | -55.69 | -162.48 | -5.32 | 30.29 |
| GACGUC | -50.99 | -147.95 | -5.12 | 28.55 | -51.05 | -146.19 | -5.73 | 32.10 |
| GCAUGC | -52.45 | -150.48 | -5.80 | 32.59 | -52.46 | -149.71 | -6.05 | 34.06 |
| GCCGGC | -54.84 | -154.57 | -6.92 | 39.11 | -55.31 | -153.08 | -7.85 | 44.45 |
| GCGAGC | -55.21 | -155.68 | -6.95 | 39.23 | -55.37 | -155.54 | -7.15 | 40.39 |
| GCGCGC | -50.02 | -136.12 | -7.83 | 45.12 | -50.62 | -136.49 | -8.31 | 48.14 |
| GCUAGC | -52.18 | -151.56 | -5.19 | 29.12 | -52.13 | -150.16 | -5.58 | 31.31 |
| GGAUCC | -52.23 | -151.12 | -5.38 | 30.19 | -52.32 | -149.39 | -6.01 | 33.79 |
| GGCGCC | -50.52 | -143.87 | -5.92 | 33.15 | -50.90 | -142.04 | -6.87 | 38.93 |
| GGGACC | -56.99 | -162.55 | -6.60 | 37.27 | -57.23 | -161.14 | -7.28 | 40.99 |
| GUGAAC | -48.66 | -141.03 | -4.94 | 27.04 | -48.70 | -139.53 | -5.44 | 30.10 |
| UCAUGA | -49.97 | -149.52 | -3.62 | 19.80 | -49.66 | -147.24 | -4.01 | 21.91 |
| UGAUCA | -41.74 | -125.10 | -2.96 | 12.46 | -41.70 | -122.22 | -3.82 | 17.92 |
| CCCGGG | -51.59 | -143.74 | -7.04 | 39.93 | -52.00 | -143.37 | -7.55 | 43.09 |
| ACCGCA | -45.29 | -128.86 | -5.35 | 28.97 | -45.60 | -127.53 | -6.07 | 33.74 |
| AGUUGC | -49.79 | -146.48 | -4.38 | 24.06 | -49.74 | -144.30 | -5.01 | 27.65 |
| AUGCGC | -45.13 | -129.54 | -4.97 | 26.51 | -45.31 | -128.20 | -5.57 | 30.42 |
| CCAACG | -53.16 | -153.31 | -5.63 | 31.72 | -53.09 | -152.48 | -5.82 | 32.79 |
| CGGACG | -58.72 | -169.39 | -6.20 | 35.15 | -58.72 | -168.06 | -6.62 | 37.33 |
| CGGUGC | -53.27 | -150.71 | -6.55 | 36.97 | -53.56 | -149.74 | -7.14 | 40.45 |
| CGUGCC | -52.56 | -151.90 | -5.47 | 30.73 | -52.69 | -150.04 | -6.18 | 34.82 |
| CGUUGA | -46.55 | -137.25 | -4.01 | 20.92 | -46.46 | -135.31 | -4.51 | 23.97 |
| CGUUGC | -50.54 | -145.52 | -5.43 | 30.27 | -50.66 | -144.08 | -6.00 | 33.63 |
| CGUUGU | -45.79 | -135.30 | -3.85 | 19.73 | -45.77 | -132.95 | -4.55 | 24.04 |
| GCCUGC | -54.04 | -152.41 | -6.80 | 38.39 | -54.23 | -152.14 | -7.07 | 39.96 |
| UGCGCA | -46.27 | -132.86 | -5.08 | 27.47 | -46.57 | -130.92 | -5.99 | 33.29 |
| UGUUGC | -49.01 | -149.03 | -2.81 | 15.00 | -48.66 | -145.81 | -3.46 | 18.48 |
| CAAUCG | -54.22 | -159.72 | -4.71 | 26.77 | -53.81 | -158.72 | -4.61 | 26.18 |
| CACGGC | -54.86 | -153.53 | -7.26 | 41.05 | -55.20 | -153.11 | -7.74 | 43.82 |
| CGAUUG | -49.63 | -147.57 | -3.89 | 21.19 | -49.37 | -145.54 | -4.26 | 23.22 |
| GCACCG | -55.36 | -159.80 | -5.82 | 32.92 | -55.40 | -158.34 | -6.32 | 35.67 |
| GGCACG | -48.53 | -138.09 | -5.73 | 31.82 | -48.88 | -136.54 | -6.55 | 36.99 |
| CCCAGGG | -61.14 | -164.42 | -10.17 | 56.48 | -61.89 | -165.55 | -10.57 | 58.52 |
| CAAAAAG | -54.06 | -154.13 | -6.28 | 35.42 | -53.81 | -154.93 | -5.79 | 32.64 |
| AAUACCG | -59.16 | -174.51 | -5.07 | 29.39 | -58.90 | -172.44 | -5.44 | 31.25 |
| AGCCGUG | -54.12 | -155.71 | -5.85 | 33.02 | -54.36 | -153.71 | -6.71 | 37.88 |
| AGCUUCA | -56.94 | -165.59 | -5.61 | 31.91 | -56.81 | -164.22 | -5.90 | 33.47 |
| GGACUUA | -57.17 | -164.83 | -6.07 | 34.38 | -57.18 | -163.55 | -6.48 | 36.57 |
| ACGUAUG | -53.80 | -159.88 | -4.24 | 24.19 | -53.71 | -156.80 | -5.10 | 28.84 |
| CACGGCU | -62.81 | -177.43 | -7.81 | 43.31 | -63.17 | -176.21 | -8.55 | 47.10 |

| | | | | | | | | |
|---|---|---|---|---|---|---|---|---|
| CAUACGU | -60.17 | -174.00 | -6.23 | 35.34 | -60.07 | -172.88 | -6.48 | 36.59 |
| UAAGUCC | -57.94 | -167.27 | -6.09 | 34.54 | -57.91 | -166.08 | -6.42 | 36.29 |
| UGAAGCU | -56.96 | -163.74 | -6.20 | 35.08 | -56.97 | -162.71 | -6.53 | 36.86 |
| AAAAAAAA | -59.19 | -174.52 | -5.09 | 29.50 | -58.61 | -173.93 | -4.69 | 27.42 |
| UAGAUCUA | -61.21 | -178.15 | -5.98 | 34.10 | -61.06 | -176.59 | -6.32 | 35.82 |
| UCUAUAGA | -60.92 | -176.20 | -6.30 | 35.68 | -60.85 | -174.90 | -6.63 | 37.36 |
| CAAAAAAG | -62.00 | -176.69 | -7.23 | 40.39 | -61.69 | -177.69 | -6.61 | 37.25 |
| CAAGCUUG | -71.48 | -205.31 | -7.83 | 42.61 | -71.19 | -205.25 | -7.56 | 41.41 |
| CAUCGAUG | -63.94 | -178.86 | -8.50 | 46.70 | -64.16 | -179.14 | -8.63 | 47.35 |
| CGAUAUCG | -72.03 | -210.24 | -6.86 | 38.27 | -71.79 | -208.32 | -7.21 | 39.81 |
| CGUCGACG | -73.23 | -207.53 | -8.89 | 47.19 | -73.43 | -206.60 | -9.39 | 49.41 |
| GAAGCUUC | -67.62 | -198.78 | -6.00 | 34.45 | -67.35 | -196.53 | -6.42 | 36.36 |
| GAUCGAUC | -70.52 | -201.03 | -8.20 | 44.37 | -70.58 | -200.21 | -8.51 | 45.82 |
| GAUGCAUC | -67.89 | -196.33 | -7.02 | 39.14 | -67.84 | -194.66 | -7.49 | 41.33 |
| GGAAUUCC | -68.39 | -196.90 | -7.35 | 40.61 | -68.39 | -195.39 | -7.82 | 42.84 |
| GGACGUCC | -69.36 | -193.63 | -9.33 | 49.92 | -69.92 | -192.64 | -10.20 | 54.02 |
| GGAGCUCC | -70.98 | -200.92 | -8.69 | 46.60 | -71.28 | -199.70 | -9.38 | 49.75 |
| GGUAUACC | -63.76 | -182.99 | -7.03 | 39.32 | -64.16 | -180.17 | -8.31 | 45.69 |
| GUACGUAC | -66.59 | -190.46 | -7.54 | 41.66 | -66.81 | -188.66 | -8.33 | 45.42 |
| GUAGCUAC | -65.44 | -188.74 | -6.93 | 38.76 | -65.51 | -186.88 | -7.57 | 41.87 |
| GUUGCAAC | -64.73 | -186.67 | -6.86 | 38.47 | -64.86 | -184.59 | -7.64 | 42.26 |
| AAGCGUAG | -67.95 | -193.18 | -8.06 | 44.01 | -67.98 | -192.79 | -8.21 | 44.73 |
| AAUCCAGU | -62.75 | -179.36 | -7.14 | 39.93 | -62.82 | -178.36 | -7.53 | 41.89 |
| ACAUAUGU | -65.21 | -191.47 | -5.85 | 33.68 | -64.95 | -189.41 | -6.23 | 35.44 |
| ACCUAGUC | -65.53 | -187.30 | -7.47 | 41.37 | -65.60 | -186.28 | -7.85 | 43.25 |
| ACGACCUC | -72.26 | -205.30 | -8.62 | 46.08 | -72.37 | -204.55 | -8.96 | 47.63 |
| AGAGAGAG | -67.79 | -194.10 | -7.62 | 41.94 | -67.77 | -193.30 | -7.84 | 42.98 |
| AGAGCUCU | -70.02 | -202.60 | -7.21 | 39.92 | -69.87 | -201.21 | -7.50 | 41.22 |
| AGCGUAAG | -66.94 | -193.40 | -6.99 | 39.01 | -67.02 | -191.31 | -7.71 | 42.43 |
| AGUCCUGA | -64.63 | -188.33 | -6.25 | 35.52 | -64.65 | -185.76 | -7.07 | 39.46 |
| AUGCGCAU | -67.04 | -192.42 | -7.39 | 40.90 | -67.09 | -191.10 | -7.85 | 43.06 |
| CACGGCUC | -72.53 | -202.32 | -9.81 | 51.57 | -72.93 | -202.15 | -10.27 | 53.60 |
| CCAUAUGG | -66.25 | -186.64 | -8.39 | 45.80 | -66.48 | -186.20 | -8.76 | 47.62 |
| CGCGUAUA | -66.27 | -193.49 | -6.29 | 35.73 | -66.13 | -191.35 | -6.82 | 38.21 |
| CGCUGUAA | -66.21 | -188.66 | -7.72 | 42.55 | -66.41 | -187.32 | -8.34 | 45.56 |
| CGUCGUCC | -70.06 | -196.11 | -9.26 | 49.44 | -70.40 | -195.80 | -9.70 | 51.50 |
| CUAGUGGA | -70.39 | -200.33 | -8.28 | 44.79 | -70.57 | -199.11 | -8.85 | 47.40 |
| CUCACGGC | -67.86 | -190.07 | -8.94 | 48.26 | -68.37 | -188.96 | -9.79 | 52.37 |
| CUGAGUCC | -68.32 | -190.98 | -9.12 | 49.06 | -68.68 | -190.72 | -9.56 | 51.15 |
| GAAUAUUC | -59.56 | -177.94 | -4.40 | 26.15 | -59.12 | -175.50 | -4.72 | 27.65 |
| GACUAGUC | -66.61 | -193.66 | -6.57 | 37.07 | -66.57 | -191.46 | -7.22 | 40.12 |
| GAGUACUC | -69.68 | -200.88 | -7.41 | 40.83 | -69.65 | -199.38 | -7.84 | 42.82 |
| GAUUAAUC | -61.70 | -180.46 | -5.76 | 33.04 | -61.45 | -178.93 | -5.98 | 34.12 |
| GCAUAUGC | -67.78 | -194.21 | -7.57 | 41.70 | -67.82 | -193.01 | -7.99 | 43.67 |
| GCCAGUUA | -65.61 | -186.22 | -7.88 | 43.40 | -65.80 | -185.31 | -8.36 | 45.71 |
| GGACCUCG | -69.90 | -196.40 | -9.01 | 48.27 | -70.26 | -195.63 | -9.62 | 51.10 |
| GGAGCACG | -71.88 | -204.27 | -8.56 | 45.87 | -72.15 | -202.83 | -9.27 | 49.11 |
| GGUGCCAA | -68.46 | -196.21 | -7.64 | 41.97 | -68.85 | -193.45 | -8.89 | 47.85 |
| GUCGAACA | -66.15 | -192.37 | -6.52 | 36.81 | -66.32 | -189.27 | -7.65 | 42.18 |
| GUCUAGAC | -64.67 | -185.57 | -7.14 | 39.83 | -64.82 | -183.87 | -7.82 | 43.14 |
| UAGGCCUA | -70.85 | -201.66 | -8.33 | 44.96 | -71.02 | -200.48 | -8.87 | 47.44 |
| UAUGCAUA | -58.44 | -171.33 | -5.33 | 30.63 | -58.42 | -168.78 | -6.10 | 34.57 |
| UGAGCUCA | -67.07 | -193.54 | -7.08 | 39.41 | -67.06 | -191.94 | -7.56 | 41.71 |
| AAUGUCGC | -65.25 | -186.38 | -7.48 | 41.43 | -65.41 | -185.07 | -8.03 | 44.16 |
| ACUGGAUU | -66.76 | -193.03 | -6.92 | 38.69 | -66.77 | -191.15 | -7.52 | 41.51 |
| CAACAGCA | -68.39 | -194.52 | -8.09 | 44.10 | -68.41 | -194.11 | -8.24 | 44.81 |
| CUACGCUU | -67.86 | -192.93 | -8.05 | 43.95 | -67.91 | -192.42 | -8.26 | 44.99 |
| CUUACGCU | -70.68 | -202.66 | -7.85 | 42.77 | -70.66 | -201.58 | -8.17 | 44.23 |

| | | | | | | | | |
|---|---|---|---|---|---|---|---|---|
| GACUAGGU | -69.13 | -196.68 | -8.16 | 44.36 | -69.32 | -195.49 | -8.72 | 46.98 |
| GAGCCGUG | -68.71 | -192.07 | -9.17 | 49.25 | -69.16 | -191.45 | -9.81 | 52.28 |
| GAGGUCGU | -73.69 | -209.42 | -8.77 | 46.57 | -73.90 | -208.19 | -9.36 | 49.20 |
| GCCGUGAG | -69.36 | -195.30 | -8.82 | 47.43 | -69.84 | -193.82 | -9.75 | 51.86 |
| GCGACAUU | -65.21 | -192.02 | -5.68 | 32.88 | -65.47 | -187.38 | -7.38 | 40.93 |
| UAACUGGC | -67.88 | -194.41 | -7.61 | 41.89 | -67.95 | -193.14 | -8.08 | 44.09 |
| UCCACUAG | -65.86 | -186.36 | -8.09 | 44.39 | -66.02 | -185.88 | -8.40 | 45.88 |
| UGCUGUUG | -64.00 | -183.70 | -7.05 | 39.41 | -64.10 | -182.20 | -7.61 | 42.19 |
| UGUUCGAC | -66.64 | -194.71 | -6.28 | 35.70 | -66.68 | -191.71 | -7.25 | 40.22 |
| UUACAGCG | -67.85 | -197.01 | -6.78 | 38.02 | -67.83 | -194.85 | -7.42 | 41.01 |
| UUGGCACC | -60.43 | -175.61 | -5.99 | 34.10 | -60.78 | -172.01 | -7.46 | 41.67 |
| CAAAAAAAG | -71.55 | -204.57 | -8.14 | 43.98 | -71.16 | -205.44 | -7.47 | 41.03 |
| CAAACAAAG | -75.70 | -215.02 | -9.04 | 47.49 | -75.48 | -215.73 | -8.60 | 45.61 |
| CAAAUAAAG | -72.94 | -209.50 | -8.00 | 43.22 | -72.59 | -209.69 | -7.59 | 41.44 |
| CAAAGAAAG | -74.88 | -213.27 | -8.77 | 46.41 | -74.67 | -213.68 | -8.43 | 44.97 |
| AAAAAAAAA | -65.66 | -192.25 | -6.07 | 34.69 | -65.05 | -192.01 | -5.53 | 32.16 |
| AUAACUGGC | -75.93 | -216.69 | -8.76 | 46.23 | -76.06 | -215.35 | -9.30 | 48.58 |
| AUCUAUCCG | -72.09 | -208.27 | -7.53 | 41.23 | -72.02 | -206.66 | -7.96 | 43.13 |
| CGCUGUUAC | -77.32 | -223.80 | -7.94 | 42.62 | -77.20 | -221.97 | -8.39 | 44.53 |
| GCCAGUUAA | -73.44 | -206.92 | -9.29 | 48.97 | -73.86 | -205.63 | -10.12 | 52.68 |
| CAACAGCAA | -71.64 | -202.72 | -8.79 | 46.97 | -71.85 | -201.94 | -9.25 | 49.05 |
| CAACAGCAU | -75.31 | -214.60 | -8.78 | 46.42 | -75.40 | -213.60 | -9.18 | 48.16 |
| CGCUGUUAG | -72.95 | -207.95 | -8.48 | 45.38 | -73.18 | -206.29 | -9.23 | 48.75 |
| CUAACAGCG | -75.48 | -212.57 | -9.59 | 49.95 | -75.67 | -212.38 | -9.83 | 51.00 |
| GUAACAGCG | -76.50 | -217.79 | -8.98 | 47.14 | -76.70 | -216.38 | -9.63 | 49.90 |
| UUAACUGGC | -69.28 | -195.07 | -8.80 | 47.39 | -69.70 | -193.86 | -9.60 | 51.14 |
| AUGAGCUCAU | -86.56 | -250.51 | -8.90 | 45.59 | -86.40 | -248.51 | -9.36 | 47.37 |
| GCGAAUUCGC | -86.48 | -242.82 | -11.21 | 54.59 | -86.79 | -242.45 | -11.63 | 56.21 |
| GCGAAAAGCG | -86.75 | -244.63 | -10.91 | 53.35 | -86.93 | -244.27 | -11.21 | 54.49 |
| AUCAAUCAUA | -70.40 | -213.14 | -4.32 | 27.44 | -70.10 | -207.78 | -5.68 | 33.16 |
| UUGUAGUCAU | -79.05 | -230.01 | -7.75 | 41.70 | -79.10 | -226.74 | -8.81 | 46.08 |
| GAAAUGAAAG | -83.48 | -240.32 | -8.98 | 46.25 | -83.20 | -239.63 | -8.91 | 46.01 |
| CCAACUUCUU | -84.30 | -239.04 | -10.20 | 50.97 | -84.21 | -239.27 | -10.04 | 50.34 |
| AUCGUCUGGA | -80.92 | -235.16 | -8.02 | 42.66 | -81.08 | -231.48 | -9.32 | 47.90 |
| AGCGUAAGUC | -85.96 | -247.60 | -9.21 | 46.83 | -86.19 | -244.51 | -10.39 | 51.39 |
| CGAUCUGCGA | -92.86 | -264.50 | -10.86 | 52.03 | -92.96 | -263.18 | -11.38 | 53.91 |
| UGGCGAGCAC | -83.95 | -232.05 | -12.02 | 58.53 | -84.81 | -231.00 | -13.20 | 63.33 |
| GAUGCGCUCG | -86.49 | -245.39 | -10.42 | 51.46 | -86.96 | -243.03 | -11.62 | 56.13 |
| GGGACCGCCU | -89.97 | -247.74 | -13.17 | 61.57 | -90.99 | -246.60 | -14.54 | 66.79 |
| AAAAAAAAAA | -74.12 | -216.77 | -6.92 | 38.50 | -73.51 | -216.18 | -6.49 | 36.71 |
| CGGCAAGCGC | -91.55 | -252.38 | -13.31 | 61.65 | -92.26 | -252.49 | -13.99 | 64.14 |
| UAGGUUAUAA | -77.55 | -225.78 | -7.56 | 41.02 | -77.54 | -222.81 | -8.47 | 44.81 |
| CGUACACAUGC | -91.83 | -258.76 | -11.62 | 55.04 | -92.36 | -256.96 | -12.71 | 59.08 |
| CCAUUGCUACC | -88.20 | -245.87 | -11.98 | 57.28 | -88.87 | -244.76 | -12.99 | 61.17 |
| CCAUCGCUACC | -87.51 | -240.79 | -12.86 | 61.04 | -88.38 | -240.33 | -13.87 | 64.96 |
| ACGUAUUAUGC | -86.75 | -257.67 | -6.87 | 38.08 | -86.61 | -252.31 | -8.40 | 43.69 |
| GCAUAAUACGU | -87.71 | -250.12 | -10.17 | 50.28 | -88.01 | -247.86 | -11.17 | 54.13 |
| CGCGAAUUCGCG | -102.98 | -290.77 | -12.84 | 57.10 | -103.67 | -287.91 | -14.42 | 62.40 |
| CUGACAAGUGUC | -105.66 | -299.42 | -12.84 | 56.56 | -106.03 | -297.51 | -13.80 | 59.68 |
| UUUUAUAAUAAA | -89.08 | -262.03 | -7.85 | 41.54 | -88.67 | -259.00 | -8.38 | 43.46 |
| AUUGGAUACAAA | -93.34 | -269.42 | -9.82 | 48.18 | -93.67 | -265.32 | -11.42 | 53.94 |
| CAACCAACCAAC | -102.45 | -287.82 | -13.23 | 58.53 | -102.90 | -286.79 | -13.99 | 61.10 |
| CUUCCUUCCUUC | -109.76 | -313.13 | -12.69 | 55.30 | -109.58 | -312.55 | -12.69 | 55.34 |
| GGAACAAGAUGC | -99.46 | -280.96 | -12.37 | 56.18 | -100.02 | -278.70 | -13.62 | 60.52 |
| GGAACUUGAUGC | -101.00 | -287.45 | -11.89 | 54.24 | -101.41 | -284.77 | -13.13 | 58.43 |
| AAUGGAUUACAA | -89.13 | -255.12 | -10.04 | 49.57 | -89.57 | -251.77 | -11.52 | 55.15 |
| GUCAGGAAUCUG | -95.26 | -267.26 | -12.41 | 57.26 | -96.00 | -265.06 | -13.84 | 62.39 |
| UUGUAAUCCAUU | -94.93 | -279.59 | -8.25 | 42.59 | -94.79 | -274.75 | -9.62 | 47.30 |

| | | | | | | | | |
|---|---|---|---|---|---|---|---|---|
| UUUGUAUCCAAU | -87.94 | -258.32 | -7.87 | 41.64 | -88.13 | -252.87 | -9.74 | 48.57 |
| CGCAUGGGUACGC | -111.96 | -307.91 | -16.51 | 67.18 | -113.34 | -306.00 | -18.48 | 73.41 |
| AGCCUAACUCAGC | -107.93 | -307.35 | -12.65 | 55.49 | -108.18 | -305.14 | -13.59 | 58.50 |
| CAUAUGGCCAUAUG | -109.31 | -308.10 | -13.80 | 58.95 | -110.16 | -304.82 | -15.66 | 64.88 |
| GUAUACCGGUAUAC | -118.01 | -339.97 | -12.62 | 53.73 | -118.49 | -334.68 | -14.74 | 59.93 |
| ACAUUAUUAUUACA | -108.93 | -318.05 | -10.34 | 48.08 | -108.71 | -314.08 | -11.34 | 51.22 |
| UACUAACAUUAACUA | -116.88 | -344.12 | -10.21 | 46.92 | -116.77 | -337.84 | -12.05 | 52.22 |
| AUACUUACUGAUUAG | -118.82 | -347.54 | -11.09 | 49.22 | -118.84 | -341.67 | -12.93 | 54.50 |
| GUACACUGUCUUAUA | -122.25 | -352.60 | -12.95 | 54.04 | -122.81 | -346.60 | -15.37 | 60.89 |
| GUAUGAGAGACUUUA | -118.43 | -343.36 | -11.99 | 51.83 | -118.92 | -336.89 | -14.49 | 59.09 |
| UUCUACCUAUGUGAU | -117.22 | -339.50 | -11.97 | 51.95 | -117.51 | -334.15 | -13.93 | 57.69 |
| AGUAGUAAUCACACC | -114.23 | -329.83 | -11.98 | 52.39 | -115.24 | -321.89 | -15.45 | 62.88 |
| AUCGUCUCGGUAUAA | -120.82 | -345.97 | -13.56 | 56.02 | -121.49 | -340.74 | -15.86 | 62.65 |
| ACGACAGGUUUACCA | -124.41 | -356.06 | -14.03 | 56.74 | -125.35 | -349.64 | -16.96 | 65.00 |
| CUUUCAUGUCCGCAU | -122.01 | -343.97 | -15.38 | 61.11 | -122.79 | -340.92 | -17.11 | 66.07 |
| UGGAUGUGUGAACAC | -119.88 | -347.55 | -12.14 | 52.08 | -120.62 | -339.98 | -15.22 | 60.94 |
| ACCCCGCAAUACAUG | -120.01 | -338.98 | -14.92 | 60.17 | -120.95 | -334.91 | -17.12 | 66.62 |
| GCAGUGGAUGUGAGA | -128.03 | -358.45 | -16.92 | 64.23 | -129.22 | -354.80 | -19.23 | 70.65 |
| GGUCCUUACUUGGUG | -133.69 | -380.40 | -15.76 | 59.86 | -134.34 | -376.18 | -17.72 | 65.03 |
| CGCCUCAUGCUCAUC | -119.40 | -333.94 | -15.88 | 63.19 | -120.59 | -330.39 | -18.16 | 69.96 |
| AAAUAGCCGGGCCGC | -134.88 | -381.04 | -16.76 | 62.29 | -135.85 | -376.73 | -19.06 | 68.36 |
| CCAGCCAGUCUCUCC | -129.39 | -363.37 | -16.75 | 63.43 | -130.35 | -360.17 | -18.70 | 68.78 |
| GACGACAAGACCGCG | -129.62 | -360.70 | -17.80 | 66.38 | -130.94 | -357.55 | -20.10 | 72.70 |
| CAGCCUCGUCGCAGC | -128.41 | -356.72 | -17.83 | 66.76 | -129.84 | -353.36 | -20.30 | 73.64 |
| CUCGCGGUCGAAGCG | -133.57 | -369.09 | -19.15 | 69.22 | -134.79 | -367.74 | -20.79 | 73.54 |
| GCGUCGGUCCGGGCU | -139.14 | -384.99 | -19.80 | 69.53 | -140.86 | -381.31 | -22.66 | 76.93 |
| CAACUUGAUAUUAUUA | -127.69 | -373.76 | -11.82 | 50.27 | -127.58 | -367.86 | -13.54 | 54.89 |
| CAUAUUGGCCAAUAUG | -128.30 | -366.76 | -14.60 | 57.67 | -128.83 | -362.27 | -16.53 | 62.95 |
| GUAUAACCGGUUAUAC | -132.13 | -380.73 | -14.10 | 55.70 | -132.63 | -374.85 | -16.42 | 61.85 |
| CGCGUACGCGUACGCG | -140.99 | -401.68 | -16.47 | 60.37 | -142.44 | -393.50 | -20.46 | 70.45 |
| CAACUUGAUAUUAAUA | -119.73 | -355.11 | -9.65 | 45.15 | -119.58 | -347.53 | -11.85 | 51.28 |
| UAUGUAUAUUUUGUAAUCAG | -150.43 | -446.03 | -12.16 | 48.90 | -150.88 | -433.55 | -16.48 | 58.73 |
| UUCAAGUUAAACAUUCUAUC | -155.56 | -468.80 | -10.23 | 44.41 | -155.34 | -455.19 | -14.23 | 53.02 |
| UGAUUCUACCUAUGUGAUUU | -161.21 | -471.35 | -15.09 | 54.23 | -161.83 | -460.58 | -19.05 | 62.85 |
| GAGAUUGUUUCCCUUUCAAA | -151.58 | -436.50 | -16.26 | 58.13 | -152.89 | -426.48 | -20.68 | 68.47 |
| AUGCAAUGCUACAUAUUCGC | -163.15 | -483.92 | -13.14 | 49.94 | -163.50 | -470.87 | -17.53 | 59.22 |
| CCACUAUACCAUCUAUGUAC | -154.29 | -444.74 | -16.42 | 58.10 | -155.56 | -434.60 | -20.84 | 68.25 |
| CCAUCAUUGUGUCUACCUCA | -164.75 | -488.23 | -13.40 | 50.35 | -166.33 | -469.77 | -20.70 | 65.73 |
| CGGGACCAACUAAAGGAAAU | -169.64 | -486.06 | -18.96 | 61.37 | -171.05 | -476.23 | -23.42 | 70.82 |
| UAGUGGCGAUUAGAUUCUGC | -169.24 | -486.89 | -18.31 | 60.05 | -170.63 | -476.18 | -23.02 | 70.02 |
| AGCUGCAGUGGAUGUGAGAA | -162.26 | -465.80 | -17.86 | 60.13 | -164.14 | -453.64 | -23.51 | 72.62 |
| UACUUCCAGUGCUCAGCGUA | -169.60 | -484.49 | -19.41 | 62.34 | -170.85 | -476.12 | -23.25 | 70.48 |
| CAGUGAGACAGCAAUGGUCG | -171.00 | -488.32 | -19.62 | 62.55 | -172.37 | -479.42 | -23.75 | 71.25 |
| CGAGCUUAUCCCUAUCCCUC | -168.71 | -487.58 | -17.56 | 58.53 | -169.84 | -476.92 | -21.99 | 67.91 |
| CGUACUAGCGUUGGUCAUGG | -170.41 | -484.89 | -20.09 | 63.65 | -172.20 | -475.00 | -24.95 | 73.98 |
| AAGGCGAGUCAGGCUCAGUG | -176.40 | -502.10 | -20.75 | 64.03 | -178.01 | -492.84 | -25.23 | 73.24 |
| ACCGACGACGCUGAUCCGAU | -171.88 | -488.57 | -20.43 | 64.12 | -173.36 | -480.34 | -24.45 | 72.60 |
| AGCAGUCCGCCACACCCUGA | -167.10 | -476.20 | -19.48 | 62.89 | -169.06 | -465.26 | -24.82 | 74.47 |
| CAGCCUCGUUCGCACAGCCC | -165.69 | -474.92 | -18.47 | 60.93 | -167.90 | -461.54 | -24.82 | 74.76 |
| GUGGUGGGCCGUGCGCUCUG | -171.89 | -475.29 | -24.55 | 73.16 | -174.84 | -467.02 | -30.07 | 85.08 |
| GUCCACGCCCGGUGCGACGG | -175.62 | -500.47 | -20.48 | 63.60 | -177.19 | -491.13 | -24.94 | 72.79 |
| GAUAUAGCAAAAUUCUAAGUUAAUA | -199.02 | -593.11 | -15.16 | 50.90 | -199.58 | -575.02 | -21.32 | 61.67 |
| AUAACUUUACGUGUGUGACCUAUUA | -165.70 | -515.06 | -6.03 | 35.92 | -166.24 | -489.26 | -14.57 | 52.62 |
| GUUCUAUACUCUUGAAGUUGAUUAC | -196.75 | -577.64 | -17.68 | 55.48 | -198.11 | -560.49 | -24.36 | 67.51 |
| CCCUGCACUUUAACUGAAUUGUUUA | -186.49 | -564.52 | -11.48 | 45.32 | -187.39 | -541.59 | -19.50 | 59.91 |
| UAACCAUACUGAAUACCUUUUGACG | -200.14 | -595.90 | -15.41 | 51.25 | -200.88 | -577.18 | -21.96 | 62.65 |
| UCCACACGGUAGUAAAAUUAGGCUU | -213.24 | -624.79 | -19.56 | 57.03 | -214.45 | -607.93 | -26.00 | 67.80 |
| UUCCAAAAGGAGUUAUGAGUUGCGA | -183.82 | -544.34 | -15.08 | 51.97 | -185.44 | -524.53 | -22.83 | 66.74 |

| | | | | | | | | |
|---|---|---|---|---|---|---|---|---|
| AAUAUCUCUCAUGCGCCAAGCUACA | -202.78 | -595.12 | -18.29 | 55.94 | -204.35 | -576.74 | -25.56 | 68.69 |
| UAGUAUAUCGCAGCAUCAUACAGGC | -194.64 | -585.14 | -13.24 | 47.93 | -195.92 | -561.68 | -21.80 | 63.05 |
| UGGAUUCUACUCAACCUUAGUCUGG | -203.73 | -594.19 | -19.53 | 57.99 | -205.33 | -577.48 | -26.32 | 69.91 |
| CGGAAUCCAUGUUACUUCGGCUAUC | -189.48 | -564.98 | -14.34 | 50.18 | -191.13 | -542.70 | -22.89 | 65.87 |
| CUGGUCUGGAUCUGAGAACUUCAGG | -209.55 | -617.82 | -18.03 | 54.85 | -211.37 | -596.54 | -26.45 | 69.10 |
| ACAGCGAAUGGACCUACGUGGCCUU | -214.41 | -605.64 | -26.66 | 68.97 | -217.36 | -592.08 | -33.81 | 81.35 |
| AGCAAGUCGAGCAGGGCCUACGUUU | -222.58 | -634.60 | -25.86 | 66.34 | -224.98 | -620.58 | -32.60 | 77.49 |
| GCGAGCGACAGGUUACUUGGCUGAU | -214.96 | -623.11 | -21.79 | 60.55 | -217.34 | -604.25 | -30.02 | 74.42 |
| AAAGGUGUCGCGGAGAGUCGUGCUG | -201.96 | -605.41 | -14.28 | 49.23 | -203.20 | -582.33 | -22.67 | 63.61 |
| AUGGGUGGGAGCCUCGGUAGCAGCC | -221.72 | -629.83 | -26.48 | 67.50 | -224.94 | -613.31 | -34.82 | 81.45 |
| CAGUGGGCUCCUGGGCGUGCUGGUC | -223.88 | -637.77 | -26.17 | 66.66 | -227.34 | -619.25 | -35.37 | 81.90 |
| GCCAACUCCGUCGCCGUUCGUGCGC | -205.46 | -602.69 | -18.63 | 56.25 | -207.68 | -581.45 | -27.43 | 71.54 |
| ACGGGUCCCCGCACCGCACCGCCAG | -221.43 | -618.76 | -29.62 | 72.94 | -225.43 | -603.85 | -38.23 | 87.58 |
| UUAUGUAUUAAGUUAUAUAGUAGUAGUAGU | -216.98 | -682.00 | -5.57 | 35.48 | -217.41 | -645.56 | -17.29 | 53.00 |
| AUUGAUAUCCUUUUCUAUUCAUCUUUCAUU | -232.01 | -707.80 | -12.59 | 45.16 | -233.13 | -676.42 | -23.44 | 61.09 |
| AAAGUACAUCAACAUAGAGAAUUGCAUUUC | -226.26 | -684.34 | -14.12 | 47.61 | -227.49 | -656.09 | -24.10 | 62.80 |
| CUUAAGAUAUGAGAACUUCAACUAAUGUGU | -233.79 | -686.62 | -20.94 | 57.22 | -235.98 | -663.43 | -30.32 | 71.61 |
| CUCAACUUGCGGUAAAUAAAUCGCUUAAUC | -224.59 | -704.75 | -6.12 | 36.29 | -224.75 | -668.89 | -17.39 | 52.60 |
| UAUUGAGAACAAGUGUCCGAUUAGCAGAAA | -249.69 | -731.81 | -22.83 | 58.50 | -252.15 | -707.16 | -32.93 | 73.11 |
| GUCAUACGACUGAGUGCAACAUUGUUCAAA | -237.33 | -702.18 | -19.65 | 55.00 | -240.34 | -672.55 | -31.85 | 73.36 |
| AACCUGCAACAUGGAGUUUUUGUCUCAUGC | -237.39 | -719.44 | -14.36 | 47.43 | -238.82 | -688.42 | -25.41 | 63.46 |
| CCGUGCGGUGUGUACGUUUUAUUCAUCAUA | -245.82 | -722.24 | -21.92 | 57.56 | -248.61 | -695.49 | -33.00 | 73.80 |
| GUUCACGUCCGAAAGCUCGAAAAAGGAUAC | -244.38 | -729.68 | -18.18 | 52.37 | -246.52 | -700.16 | -29.47 | 68.66 |
| AGUCUGGUCUGGAUCUGAGAACUUCAGGCU | -240.73 | -736.29 | -12.48 | 44.71 | -242.34 | -700.77 | -25.10 | 62.58 |
| UCGGAGAAAUCACUGAGCUGCCUGAGAAGA | -240.86 | -717.45 | -18.45 | 52.99 | -243.66 | -686.13 | -30.96 | 71.40 |
| CUUCAACGGAUCAGGUAGGACUGUGGUGGG | -266.13 | -768.32 | -27.95 | 63.99 | -269.42 | -744.79 | -38.54 | 78.65 |
| ACGCCCACAGGAUUAGGCUGGCCCACAUUG | -262.12 | -760.44 | -26.39 | 62.26 | -265.12 | -736.53 | -36.80 | 76.81 |
| GUUAUUCCGCAGUCCGAUGGCAGCAGGCUC | -246.99 | -750.27 | -14.40 | 47.06 | -248.87 | -715.32 | -27.12 | 64.82 |
| UCAGUAGGCGUGACGCAGAGCUGGCGAUGG | -253.60 | -726.96 | -28.25 | 65.88 | -257.94 | -701.79 | -40.39 | 83.70 |
| CGCGCCACGUGUGAUCUACAGCCGUUCGGC | -260.01 | -757.69 | -25.13 | 60.73 | -263.45 | -730.27 | -37.07 | 77.50 |
| GACCUGACGUGGACCGCUCCUGGGCGUGGU | -244.62 | -719.30 | -21.63 | 57.25 | -248.02 | -689.60 | -34.24 | 75.85 |
| GCCCCUCCACUGGCCGACGGCAGCAGGCUC | -271.10 | -782.00 | -28.68 | 64.43 | -275.03 | -755.76 | -40.75 | 80.90 |
| CGCCGCUGCCGACUGGAGGAGCGCGGGACG | -248.47 | -745.40 | -17.40 | 51.04 | -250.98 | -712.16 | -30.21 | 69.15 |

[a] All duplexes consist of the denoted RNA strand and its complementary DNA strand.
